# fSuSiE enables fine-mapping of QTLs from genome-scale molecular profiles

**DOI:** 10.1101/2025.08.17.670732

**Authors:** William R. P. Denault, Hao Sun, Peter Carbonetto, Anjing Liu, Philip L. De Jager, David Bennett, The Alzheimer’s Disease Functional Genomics Consortium, Gao Wang, Matthew Stephens

**Author notes:** Equally significant contributions (William R. P. Denault, Hao Sun). Correspondence: William R.P. Denault; Gao Wang,; Matthew Stephens.

## Abstract

Molecular quantitative trait locus (QTL) studies seek to identify the causal variants affecting molecular traits like DNA methylation and histone modifications. However, existing fine-mapping tools are not well suited to high-dimensional molecular traits, and so analyses of these traits typically proceed by considering each variant and each molecular measurement independently, ignoring the LD among variants and the spatial correlation in effects between nearby sites. This severely limits accuracy in identifying causal variants and quantifying their molecular trait effects. Here, we introduce fSuSiE (“functional Sum of Single Effects”), a fine-mapping method that addresses these challenges by explicitly modeling the spatial structure of genetic effects on molecular traits. fSuSiE integrates wavelet-based functional regression with the computationally efficient “Sum of Single Effects” framework to simultaneously finemap causal variants and identify the molecular traits they affect. In simulations, fSuSiE identified causal variants and affected CpGs more accurately than methods that ignore spatial structure. In applications to DNA methylation and histone acetylation (H3K9ac) data from the ROSMAP study of the dorsolateral prefrontal cortex, fSuSiE achieved dramatically higher resolution than existing methods (e.g., identifying 6,355 single-variant methylation credible sets compared to only 328 from an existing approach). Applied to Alzheimer’s disease (AD) risk loci, fSuSiE identified potential causal variants colocalizing with AD GWAS signals for established genes, including *CASS4* and *CR1/CR2*, suggesting specific potential regulatory mechanisms underlying these AD risk loci.

## Introduction

Molecular quantitative trait locus (QTL) mapping [1] aims to identify genetic variants associated with a wide variety of molecular traits measured across the genome, including RNA expression [2, 3], protein expression [4, 5], DNA methylation [6–11], chromatin accessibility [12, 13], transcription-factor binding [14, 15] and histone modification [16, 17]. A critical next step, known as *fine-mapping*, is to pinpoint the individual causal variants, determine how many act independently, and estimate their effects on the (typically large number of) molecular traits [18–22]. While sophisticated fine-mapping methods exist to handle these challenges for univariate or low-dimensional traits such as the total expression level of a gene [23–39], they are ill-suited to many molecular traits such as DNA methylation that are inherently high-dimensional. For example, a two-step approach consisting of univariate association testing followed by univariate-trait fine-mapping has been used at scale (e.g., for fine-mapping expression QTLs and splice-junction QTLs [40]), but this approach will not work well for spatially structured molecular traits. The existing methods fail to leverage the fact that causal SNPs may often affect nearby univariate molecular traits in a coordinated or spatially dependent way [41, 42]. (For example, methylation QTLs frequently affect multiple CpGs within regions spanning several kb [7, 8, 10, 43–49].) This not only limits power to detect such effects, but also limits understanding of the effects of genetic variants on molecular traits. Furthermore, since most genetic effects on complex disease risk are likely mediated through effects on molecular traits, these limitations of existing analysis tools also hinder progress in understanding the genetics of complex disease.

To address these limitations, we introduce *functional SuSiE* (fSuSiE), a new method for fine-mapping high-dimensional, spatially structured molecular traits, drawing on ideas from functional data analysis [44, 50–54]. fSuSiE models the effects of each putative causal SNP as a function that varies along the genome so as to capture spatially dependent changes in the molecular trait [44–47, 50– 53, 55–60]. It overcomes the high computational burden of fine-mapping high-dimensional molecular traits by building on the “Sum of Single Effects” (SuSiE) model [24, 25], which decomposes the complex multi-SNP computations into a series of simpler single-SNP computations. The resulting method is computationally practical for fine-mapping molecular measurements made across moderately large genetic regions containing hundreds or thousands of trait measurements and thousands of genetic variants.

We validate fSuSiE’s performance using simulated methylation QTL datasets, where it simultaneously identifies causal variants and affected CpGs more accurately than methods ignoring spatial structure. To demonstrate its potential for biological discovery, we applied fSuSiE genome-wide to DNA methylation (DNAm) and H3K9ac data [61] from the ROSMAP study [62], analyzing over 400,000 CpG sites and 90,000 histone peaks. Applied to Alzheimer’s disease risk loci, fSuSiE pinpointed putative causal variants that colocalized with AD GWAS signals for established genes including *CASS4* and *CR1/CR2*, suggesting biologically coherent regulatory mechanisms underlying genetic risk. While we showcase fSuSiE using DNAm and H3K9ac, it is widely applicable to any trait that varies along the genome, such as other histone marks, chromatin accessibility, expression level variation among exons and introns, and other traits measured by high-throughput sequencing technologies.

## Results

### Overview of fSuSiE

fSuSiE takes as input an *N* ×*T* matrix, **Y**, containing molecular trait data on *N* samples at *T* genomic locations, and an *N* ×*J* matrix, **X** of genotypes for the same *N* samples at *J* genetic variants, typically SNPs (Fig. 1a–b). It then attempts to identify which SNPs affect which molecular traits by fitting a multivariate linear model,

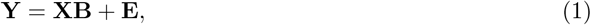

where **B** is a *J*×*T* matrix of effects whose element *b*_*jt*_ represents the effect of SNP *j* on molecular trait *t*, and **E** is a matrix of residuals. Like most fine-mapping methods, fSuSiE assumes that only a few genetic variants affect the trait: that is, most rows of **B** are zero. It does so by modeling **B** using a “sum of single effects” (SuSiE) model [24, 25], **B** = **B**^(1)^ + ···+ **B**^(*L*)^, where each matrix **B**^(*l*)^ is assumed to capture the effect of a single SNP, and so has exactly one non-zero row. (One or more **B**^(*l*)^ may be entirely zero if the data support fewer than *L* effects.)

**Fig. 1.**
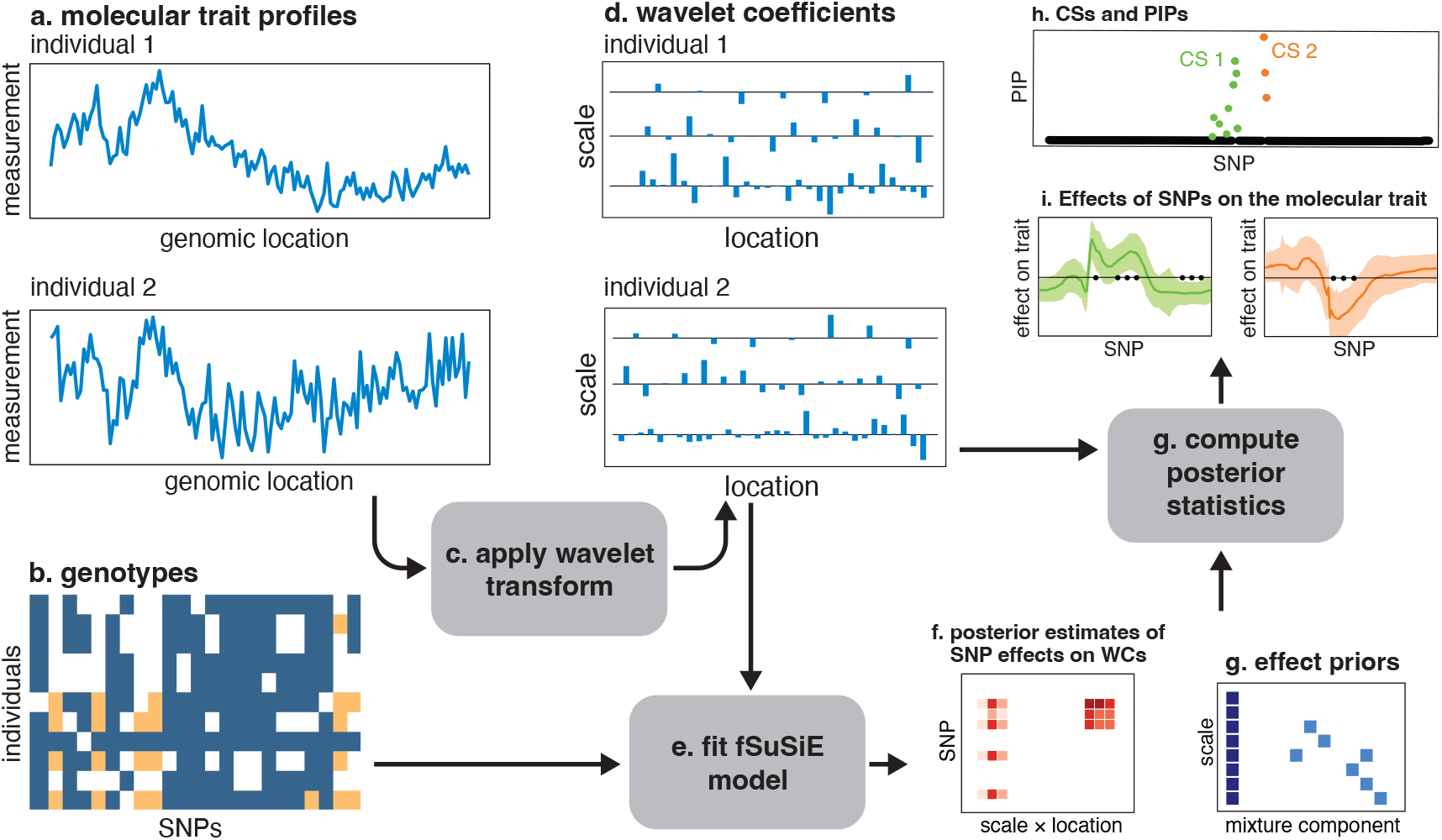
Overview of fine-mapping high-dimensional, spatially structured molecular traits using fSuSiE. The inputs to fSuSiE are an *N* ×*T* matrix, **Y**, containing *N* samples of the molecular trait measured at *T* locations (a), and an *N*× *J* genotype matrix, **X** (b). First, the discrete wavelet transform (DWT) is applied to **Y** (c), resulting in an *N* ×*T* matrix of wavelet coefficients (WCs), **D** (d). The fSuSiE model is fit to **X, D** (e); this model fitting includes estimating the effects of the SNPs on the WCs—a *J* ×*T* matrix, **F** (f)—and fitting the adaptive shrinkage priors for the effects **F** (g). Key fSuSiE outputs include: credible sets (h), each of which is intended to capture a distinct causal SNP; a posterior inclusion probability (PIP) for each SNP giving the probability that the SNP is causal for at least one trait (h); and posterior estimates of the SNP effects on **Y**, credible bands, and estimates of affected locations (i). These quantities are defined in the Online Methods.

fSuSiE uses a wavelet-based linear model [63, 64] for the effects of each causal SNP—that is, for the non-zero row in each **B**^(*l*)^. Wavelets have been used in several genomics applications to model temporally or spatially structured data [44, 50, 53, 63, 65]. Here, we use wavelets to model spatial structure in the effect of each SNP on the molecular trait. Specifically, fSuSiE applies the discrete wavelet transform (DWT) to (1), then uses flexible sparse prior distributions to shrink the DWT-transformed effects. This approach is motivated by the observation that *spatial structure* in the original signal (Fig. 1a) often corresponds to *sparse structure* in the wavelet-transformed signal (Fig. 1b). We implemented two different priors: a simple “independent shrinkage” (IS) prior that applies the same prior to all wavelet-transformed effects; and a more flexible “shrinkage-per-scale” (SPS) prior that uses a different prior for each wavelet scale. The SPS prior is more flexible and is therefore expected to perform better, but at an increased computational cost.

The key analysis questions answered by fSuSiE are: (i) which SNPs are causally affecting the observed molecular phenotypes? (ii) which molecular phenotypes are affected by each causal SNP? In terms of the model above, question (i) asks which rows of **B** (or rows of **B**^(*l*)^) are non-zero, whereas question (ii) relates to the values of the effect estimates in these non-zero rows. To answer (i), fSuSiE outputs a posterior inclusion probability (PIP) for each SNP and a “credible set” (CS) [20, 24] for each non-zero **B**^(*l*)^. The PIP for SNP *j* is the probability that the *j*th row of **B** is non-zero. The CS for each **B**^(*l*)^ is a subset of (highly correlated) SNPs that has high probability (e.g., *>*0.95) of containing the non-zero row. Within each CS, we refer to the SNP with the highest PIP as the “sentinel SNP”. To answer (ii), fSuSiE outputs, for each sentinel SNP, an estimate of its effect on each molecular trait, along with an assessment of its uncertainty: a pointwise credible band that is designed to have high probability (e.g.,*>*0.95) of containing the effect at each location. If the credible band for a particular molecular phenotype excludes zero then we say that the CS containing that sentinel SNP is”significant” for that molecular phenotype.

Fig. 1 summarizes the overall fSuSiE pipeline. Fig. 2 illustrates fSuSiE in a simulated example, contrasting it with a simple univariate QTL mapping (SNP-trait association tests) as well as a two-step univariate fine-mapping (SNP-trait association tests followed by fine-mapping the associated traits). This example illustrates how fSuSiE directly addresses the key questions of which SNPs are impacting the molecular traits (D), and which traits are affected by each causal SNP (E), whereas the other analyses do not. Further details of the fSuSiE implementation, and guidance on applying fSuSiE to genome-scale molecular trait data, are given in the Methods and Supplementary Note.

**Fig. 2.**
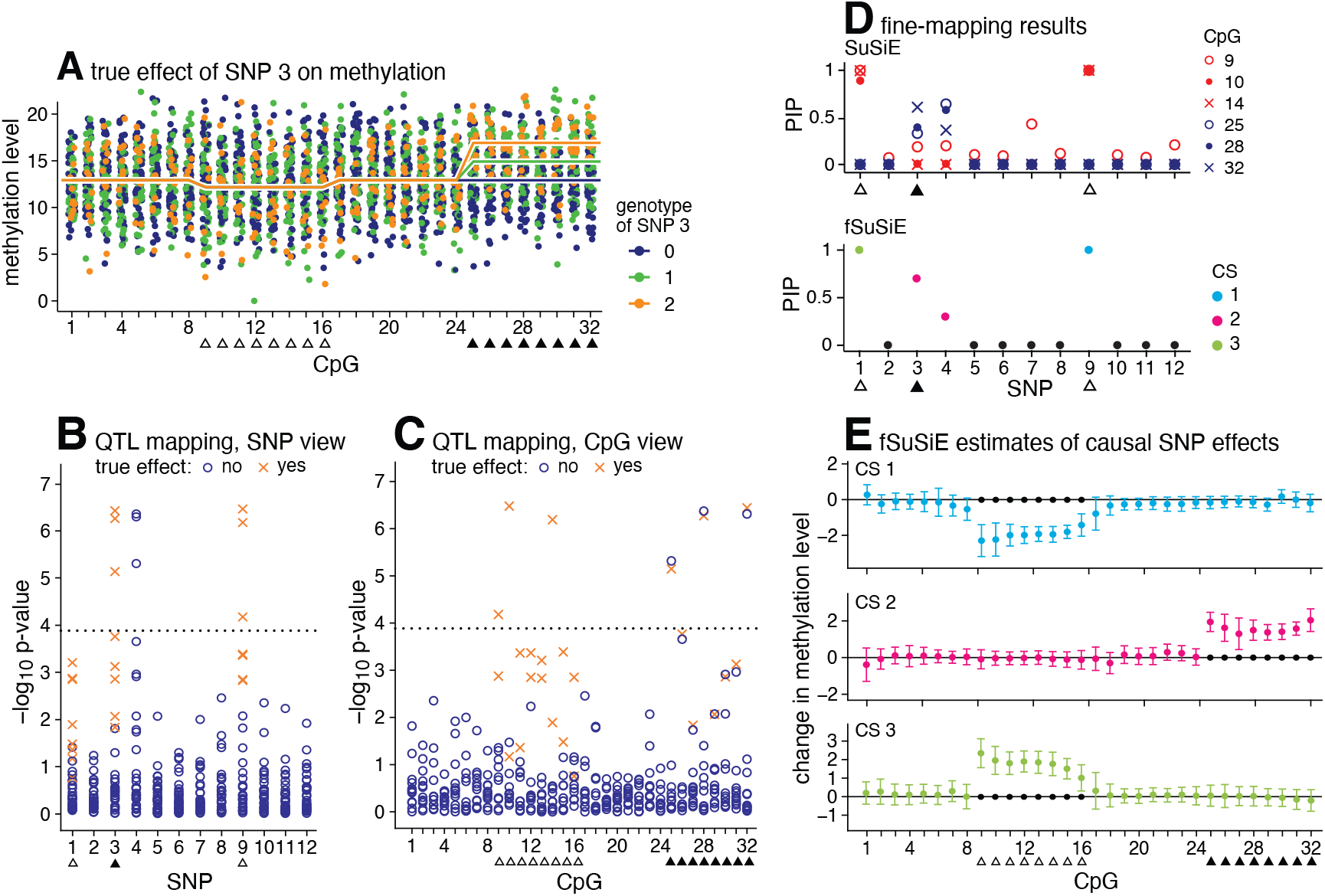
Fine-mapping with fSuSiE: an illustration. In this simulated example, the methylation levels **Y** of *N* = 100 samples are measured at 32 CpGs within a locus of interest (A). The methylation levels in two “clusters” of 16 CpGs are affected by 3 of 12 candidate SNPs. The effect of one of the causal SNPs is depicted in A. (The three curves show the mean methylation levels at the three SNP genotypes.) To map methylation QTLs, one might perform an association test for each SNP-CpG pair [10, 61, 66], resulting in 32 × 12 = 384 association tests. (The *p*-values are from two-sided *t*-tests, and are not corrected for multiple testing.) Examining the associations by SNP (B) or by CpG (C), the association tests find signals corresponding to 2 of the 3 causal SNPs, and the affected CpGs appear to “cluster”. (Perhaps all three causal SNPs would be identified with a more careful multiple testing correction; here we used Bonferroni.) The 3 causal SNPs and corresponding 8 affected CpGs are indicated by △ or ▲. (SNPs 1 and 9 affect the same CpGs.) From these same data, fSuSiE provides a more parsimonious fine-mapping: fSuSiE identifies 3 95% CSs, each containing a causal SNP, and expresses uncertainty in which SNP is causal by the PIPs (D–bottom). (Note that SNPs 3 and 4 are strongly correlated.) fSuSiE also provides 95% credible bands for each CS summarizing the CpGs that are affected by each causal SNP (E). A credible band that does not include zero indicates that the CpG is affected (•). One could instead analyze each of the 6 associated CpGs separately using a univariate fine-mapping method (e.g., SuSiE). However, this results in a less parsimonious fine-mapping result that is difficult to make sense of: first, it produces multiple fine-mapping signals, and so it is unclear which one should be used (D–top); second, it does not provide a summary similar to E indicating which CpGs are affected by each causal SNP (E). This example is also an R “vignette” available at https://stephenslab.github.io/fsusieR/articles/methyl_demo.html.

### Performance assessment in simulated methylation data sets

To validate fSuSiE we used simulated molecular QTL data sets where we could assess accuracy of the estimates by comparing to the true values. We assessed fSuSiE’s ability to perform the two key tasks highlighted above: (i) correctly identify the causal SNPs that affect the molecular trait at one or more locations; and (ii) correctly identify the molecular locations affected by the causal SNPs.

We simulated data sets containing both SNP genotypes and CpG methylation levels, in which one or more SNPs affected the methylation levels at one or more CpGs. We simulated genotypes with realistic patterns of linkage disequilibrium (LD) using sim1000G [67], with genotype data from the 1000 Genomes Phase 3 whole-genome sequencing [68] as the input source. The simulations varied in size of genomic region (median: ~1 Mb), number of SNPs (1,500–4,000 SNPs), and LD patterns. Consistent with real patterns of LD, simulated SNPs were often very strongly correlated with each other, and sometimes perfectly correlated, and therefore one should not expect 100% accuracy in identifying the causal SNPs, even in the simulation settings that were most favorable to fSuSiE.

We simulated realistic whole-genome bisulfite sequencing (WGBS) methylation levels using WGBSSuite [69], with different types of spatial structure in the methylation changes that might be motivated by different epigenetic mechanisms: constant changes in methylation across contiguous windows, with abrupt changes between windows (“WGBS block”); and changes that varied smoothly within the locus (“WGBS decay”). We also benchmarked fSuSiE in three other simulation settings in which the data conformed more closely to fSuSiE’s modeling assumptions (“wavelet simulations”), but also allowed for some departures to assess more realistic situations in which the variance due to non-genetic effects (i.e., “noise”) was spatially structured. We varied the number of CpGs (*T*), the number of SNPs (*J*), the number of causal SNPs, and the total amount of variation explained by the SNPs. Detailed descriptions of the simulation procedures are given in the Online Methods.

As far as we are aware, fSuSiE is the first available tool to perform statistical fine-mapping of spatially structured traits. Among existing approaches, perhaps the most obvious competitors are approaches that first reduce the high-dimensional trait to a single dimension, e.g. using the top principal component (PC), and then apply a univariate fine-mapping method (e.g., [70, 71]). Here we implemented a version of this approach, “SuSiE-topPC”, by running SuSiE on the top PC computed from the column-centered **Y**. This type of approach has important limitations compared with fSuSiE. First, in real applications the top PC may primarily capture non-genetic effects such as experimental batch effects or population structure [72, 73]. Second, it only identifies causal SNPs that impact some aspect of the high-dimensional trait, without identifying which specific molecular locations (CpGs) are affected by these SNPs. Nonetheless, this existing approach may work well for detecting causal SNPs that have the biggest effect on a locus, and serves as a helpful benchmark against which to compare the ability of fSuSiE to identify causal SNPs.

Both fSuSiE and SuSiE-topPC produce posterior inclusion probabilities (PIPs) that assess the probability of each SNP being a causal SNP, and 95% credible sets (CSs) that are designed to contain (with probability *>*0.95) at least one causal SNP. In fine-mapping applications, due to high LD, it may often be impossible to confidently identify the causal SNP, but it may nonetheless be possible to identify a small number of SNPs, in high LD with one another (which we call high “purity”), that likely contains a causal SNP. That is, while no single SNP may have a PIP near 1, there may be a small, high purity, 95% CS. For this reason CSs, are perhaps the most relevant fine-mapping output; however comparing PIPs is also informative, and so we assessed both PIPs and CSs here.

Comparing PIPs (Fig. 3A), both variants of fSuSiE (IS prior and SPS prior) achieved higher power than SuSiE-topPC at the same false discovery rate in all simulation scenarios. Comparing CSs (Fig. 3B, C, Supplementary Fig. 1), fSuSiE consistently achieved better power, with smaller and higher purity CSs, while maintaining coverage close to the target value (95%). Notably, fSuSiE maintained coverage close to the target value even in situations (“fractional noise”, “GP noise”) where fSuSiE’s assumptions about the spatial dependence might be expected to be violated (Fig. 3B, Supplementary Figures 5–8). This is partly explained by the decorrelating, or “whitening”, effect of wavelets on spatially structured signals [63, 74, 75]. As expected, the performance of fSuSiE also tended to improve as the density of the molecular trait measurements increased (i.e., more CpGs), and as the SNPs produced larger changes in methylation (Supplementary Figures 3–12). All methods produced well calibrated PIPs (Supplementary Fig. 2). Interestingly, the performance gains over SuSiE-topPC were among the greatest in the WGBS and GP noise simulations that did not conform to fSuSiE’s modeling assumptions. Consistent with this, in the simpler wavelet simulations the top PC captures the genetic component of variation well, whereas in the WGBS block and WGBS decay simulations the genetic effect is distributed across many CpGs and the top PC mixes genetic with non-genetic variation.

**Fig. 3.**
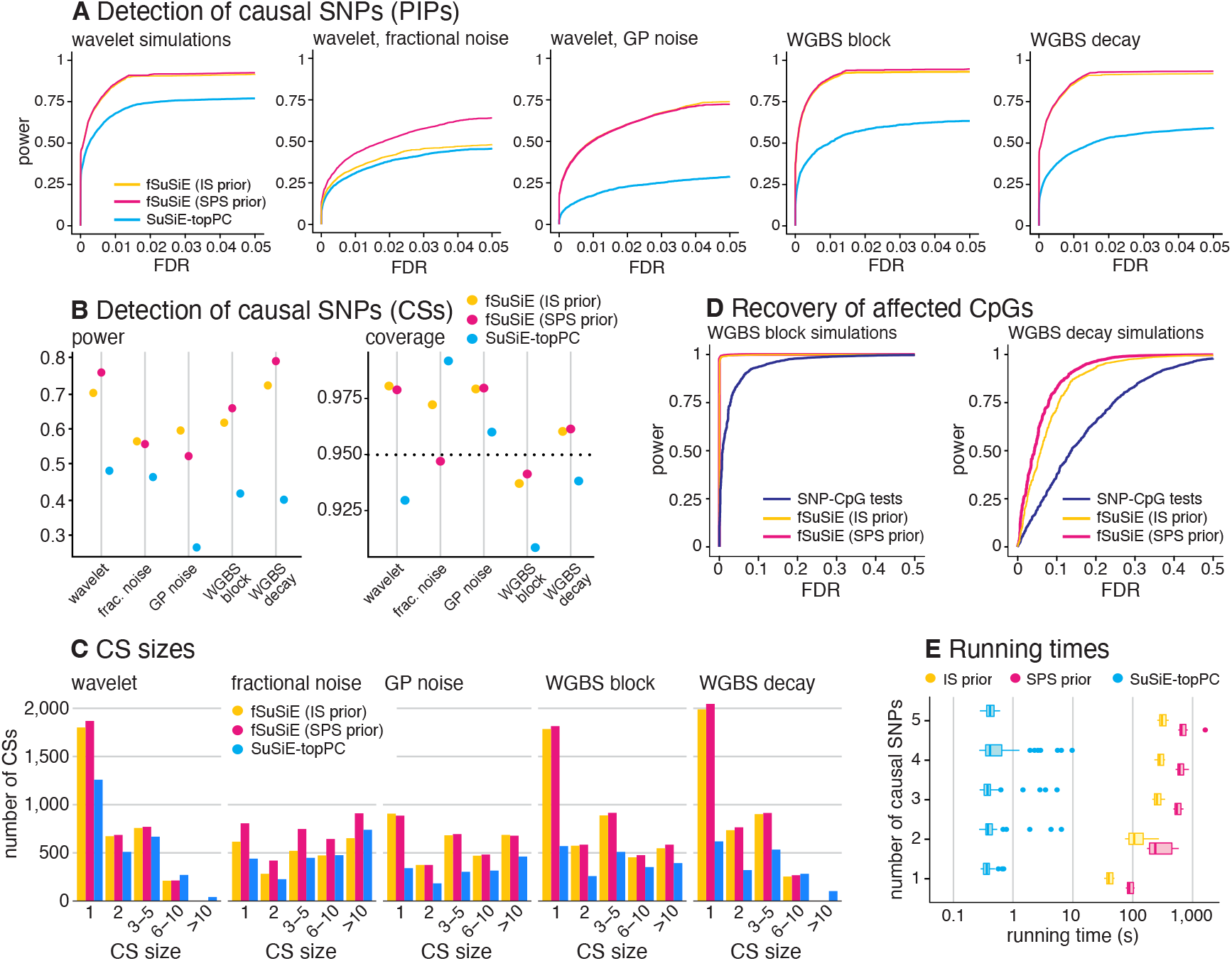
Performance assessment in simulated data sets. Performance in simulated data sets was evaluated in two separate tasks: recovery of the causal SNPs (A–C), and identification of the affected molecular trait locations (D). In A–C, we simulated data sets under five scenarios. In each of the five scenarios, *n =* 300 methylation data sets were simulated with *N* = 100 samples, *T* = 128 locations (CpGs), *J* = 1,500–4,000 SNPs, among which 1–20 were causal, and 10% total variance explained. In D, data sets were simulated in two scenarios with the same settings as above, but with *J* = 1 SNP, which was causal. Panels A, D show power vs. FDR (i.e., precision-recall with a flipped x-axis) in identifying causal SNPs (A) and affected CpGs (D). In each simulation setting, FDR and power were calculated by varying a measure threshold from 0 to 1 (*n* = 300). In A, the measure was PIP for all methods; in D, it was *α*_max_ for fSuSiE and *p-*value from the SNP-CpG tests. (See Online Methods for a definition of *α*_max_.) Power and FDR were calculated as 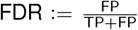 and 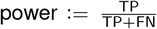, where FP, TP, FN, FN denote, respectively, the number of false positives, true positives, false negatives and true negatives. Note that the yellow and red curves (fSuSiE, IS and SPS priors) closely overlap in some plots and therefore are not always visible. Panels B, C evaluate the 95% CSs returned by fSuSiE and SuSiE-topPC: *power*, the proportion of the true causal SNPs included in at least one CS; *coverage*, the proportion of CSs containing a true causal SNP; and *CS size*, the number of SNPs in a CS. The dotted horizontal line in B shows the target coverage (95%). Panel E summarizes the running times in simulated data sets with *J =* 1,000 SNPs (*n* = 300). The box plot whiskers depict 1.5× the interquartile range, the box bounds represent the upper and lower quartiles (25th and 75th percentiles), the center line represents the median (50th percentile), and points represent outliers. See Supplementary Figures 1–16 for additional results.

One particularly striking result is the increase in the number of single-variant CSs returned by fSuSiE vs. SuSiE-topPC (Fig. 3C). In most cases (7,188 out of 7,422), these single-variant CSs contained the causal SNP, and in many cases single-variant CSs occurred even when there were other SNPs in very high LD with the causal SNP (including in cases where the other SNP differed from the causal SNP at only a single individual). To confirm this striking result, we performed additional simulations to specifically confirm that, given sufficiently strong signals, fSuSiE can indeed accurately produce a single-variant CS even in the presence of very strong LD (see Online Methods, “Additional simulations to assess single-variant CSs” and Supplementary Fig. 13). These results highlight an important general benefit of joint analysis of multiple traits: when a SNP affects multiple traits, joint analysis not only increases power to detect effects, but also substantially improves the fine-mapping resolution by reducing the typical sizes of CSs (see also [33]).

As noted above, a limitation of SuSiE-topPC (and related methods) is that it does not identify which locations (the CpGs) are affected by the causal SNPs identified. One way to address this would be to follow SuSiE-topPC with simple univariate SNP-CpG tests to determine which CpGs are significantly associated with the causal SNPs identified. A benefit of functional regression methods like fSuSiE is that they exploit spatial correlations in effects to increase power to detect such associations. To demonstrate this, we performed simulations with a single (causal) SNP affecting a subset of CpGs, and applied both fSuSiE and standard SNP-CpG tests to detect the associated CpGs. The results showed consistent gains in power and accuracy for fSuSiE, with the power gains being greatest in the more challenging WGBS decay simulations (Fig. 3D, Supplementary Figures 14 and 15). (We did not use the wavelet simulations here because in these simulations the causal SNPs produced changes, mostly small, to all CpGs.) This illustrates the ability of fSuSiE to combine borderline association signals across spatially-proximal affected CpGs to more accurately identify these signals.

Overall, the two variants of fSuSiE—one with the simpler IS prior, the other with a more flexible SPS prior—performed similarly, with a slight power advantage for the SPS prior. The tradeoff is that the SPS prior also increases running time (Fig. 3E, Supplementary Fig. 16). Thus, the choice of prior depends on weighing the benefits of additional discoveries against the additional computation. While both variants of fSuSiE take much longer to run than SuSiE-topPC, they remain practical for data sets of the scale we expect to analyze in molecular QTL studies—that is, data sets with thousands of SNPs and thousands of molecular trait measurements. We took the IS prior to be the default prior in light of its good performance at a favourable computational cost, and in subsequent fSuSiE analyses the IS prior is assumed unless stated otherwise.

### Genome-wide fine-mapping of DNA methylation and histone acetylation QTL

To demonstrate the potential of fSuSiE to generate insights into genetic regulation of molecular trait profiles, and to aid in interpretation of disease-associated genetic variants, we applied fSuSiE to DNA methylation (DNAm) and histone acetylation (H3K9ac) data from the Religious Orders Study/Memory and Aging Project (ROSMAP) with harmonized genotypes from the Alzheimer’s Disease Sequencing Project [62, 76, 77]. Array-based DNA methylation profiles of *n* = 636 donors and H3K9ac ChIP-seq profiles of *n* = 592 donors were obtained from the dorsolateral prefrontal cortex (DLPFC), a brain tissue affected by AD. The molecular trait data for fine-mapping with fSuSiE were the *n* = 636 methylation proportions (“*β* values”) at 413,433 CpGs and the *n* = 592 read counts at 92,401 H3K9ac peaks. (H3K9ac peak calling and other data processing steps are detailed in the Online Methods.) Supplementary Fig. 18 shows the trait-level spatial clustering of high-activity positions in these data at the *CASS4* (methylation) and *CR1*/*CR2* (H3K9ac) fine-mapping regions, illustrating the empirical pattern that motivates fSuSiE. We used fSuSiE to fine-map SNPs affecting methylation (mSNPs) and H3K9ac (haSNPs), as described below. We also performed SuSiE fine-mapping of gene expression and protein abundance (from the same DLPFC samples) to obtain a more complete picture of the genetic regulation in this brain region.

To apply fSuSiE genome-wide, we partitioned the genome into large genomic regions, and then (after filtering out regions containing too few CpGs or peaks) applied fSuSiE to each region separately, using the SNP genotype data and molecular trait data at all SNPs and locations within the region. To minimize regulatory interactions between regions, we defined the regions based on topologically associating domains (TAD) [78]. For brevity we refer to these regions as TADs, although they also contain boundaries between TADs; see Online Methods. In total, we decomposed the genome (autosomal chromosomes only) into 1,329 TADs ranging in size from 2 to 35 Mb (median: 4 Mb) each containing 18–10,035 CpGs (median: 450 CpGs) and 2–827 H3K9ac peaks (median: 116 peaks).

To illustrate the inferences produced by fSuSiE, Fig. 4 presents a detailed example of H3K9ac fine-mapping in a 3.7-Mb TAD on chromosome 1. (See also Supplementary Fig. 19 for a detailed methylation example.) In this TAD, we used fSuSiE to analyze the genotypes at 16,642 SNPs and H3K9ac measurements at 269 peaks for *n* = 592 individuals. fSuSiE produced 4 CSs (Fig. 4A) corresponding to 4 independent signals of genetic effects on histone acetylation levels at one or more peaks (Fig. 4B). One SNP (rs12757179 in CS 1) was identified as a haSNP with high confidence (PIP = 0.99). For the other haSNPs, fSuSiE was less certain about which SNP was causal; this uncertainty is represented by CSs containing several SNPs each. Some of the haSNPs produce coordinated changes in histone acetylation across several sites, where a single variant simultaneously modulates histone acetylation levels at multiple nearby peaks in a spatially-correlated pattern; for example, the CS1 haSNP (rs12757179) affects histone acetylation (in both directions) at 17 H3K9ac peaks (that is, their 95% credible bands did not include zero). Interestingly, all inferred changes occur at peaks near an haSNP. (The CS3 haSNP produces the longest-range changes, but the changes still remain within 200 kb of the SNP.) We also observed this tendency genome-wide in both methylation and H3K9ac (Fig. 5B). Since the fSuSiE model does not take location of SNPs into account, the fact that fSuSiE identified mostly short-range effects provides some independent support that these effects are likely real rather than false positives.

**Fig. 4.**
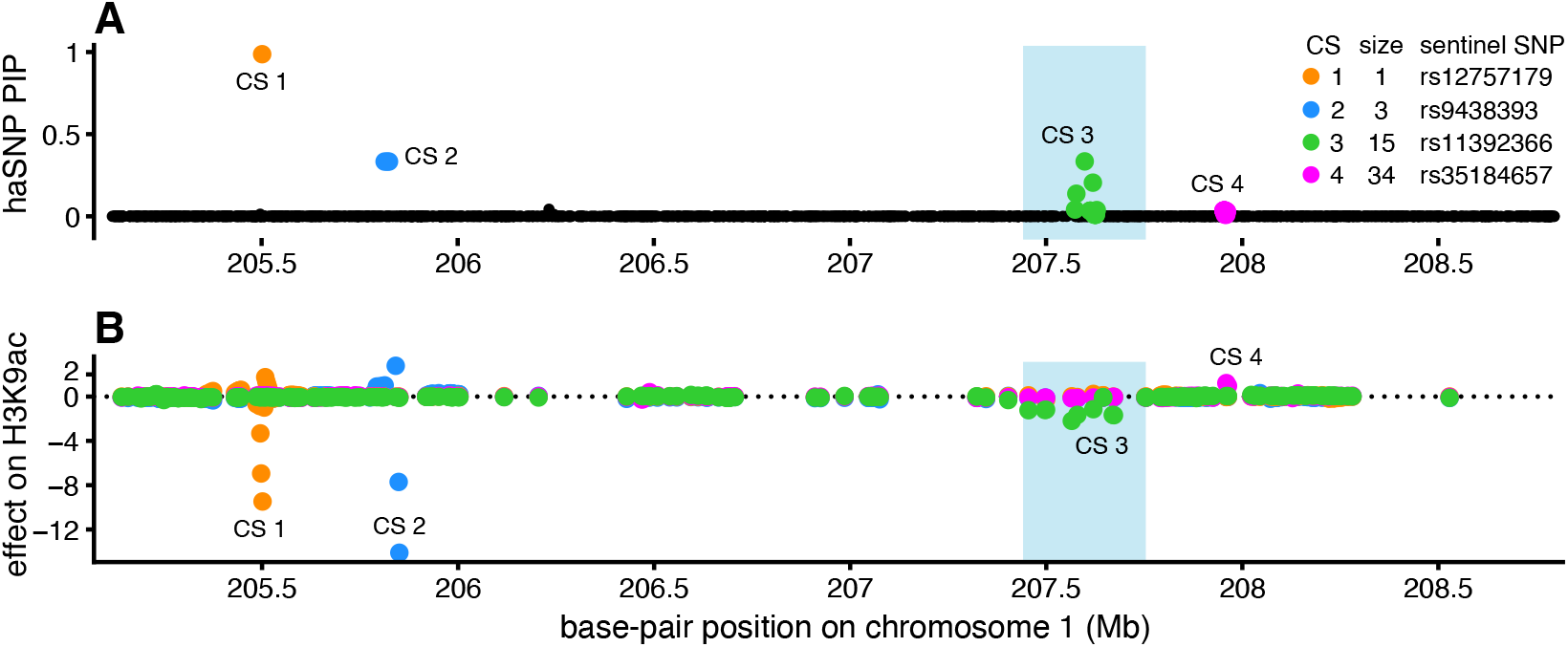
Example fSuSiE fine-mapping of H3K9ac. Panel A shows the posterior inclusion probabilities (PIPs) for all SNPs analyzed in the TAD (chromosome 1, 205.1–208.8 Mb, 16,642 SNPs, 269 H3K9ac peaks). The PIP is an estimate of the probability that the SNP affects H3K9ac levels at one or more peaks within the TAD. The colors indicate the different haSNPs, and the uncertainty in which SNP is the haSNP is represented by the 95% credible set (CS): each SNP in the CS is a candidate haSNP, with probability of being causal given by the PIP. *CS size* is the number of SNPs in the CS; *sentinel SNP* is the SNP with the highest PIP in the CS. Panel B shows the effect of each haSNP on H3K9ac levels at each of the peaks analyzed in the TAD. (When the CS contains more than one SNP, the effect of the sentinel SNP is shown.) Since the effects were estimated at different scales in this analysis, to show the effect estimates in a single plot, we divided the effects by the credible band sizes. Base-pair positions are based on Genome Reference Consortium human genome assembly 38 (hg38). The results within the blue-shaded region are examined in more detail in Fig. 6.

**Fig. 5.**
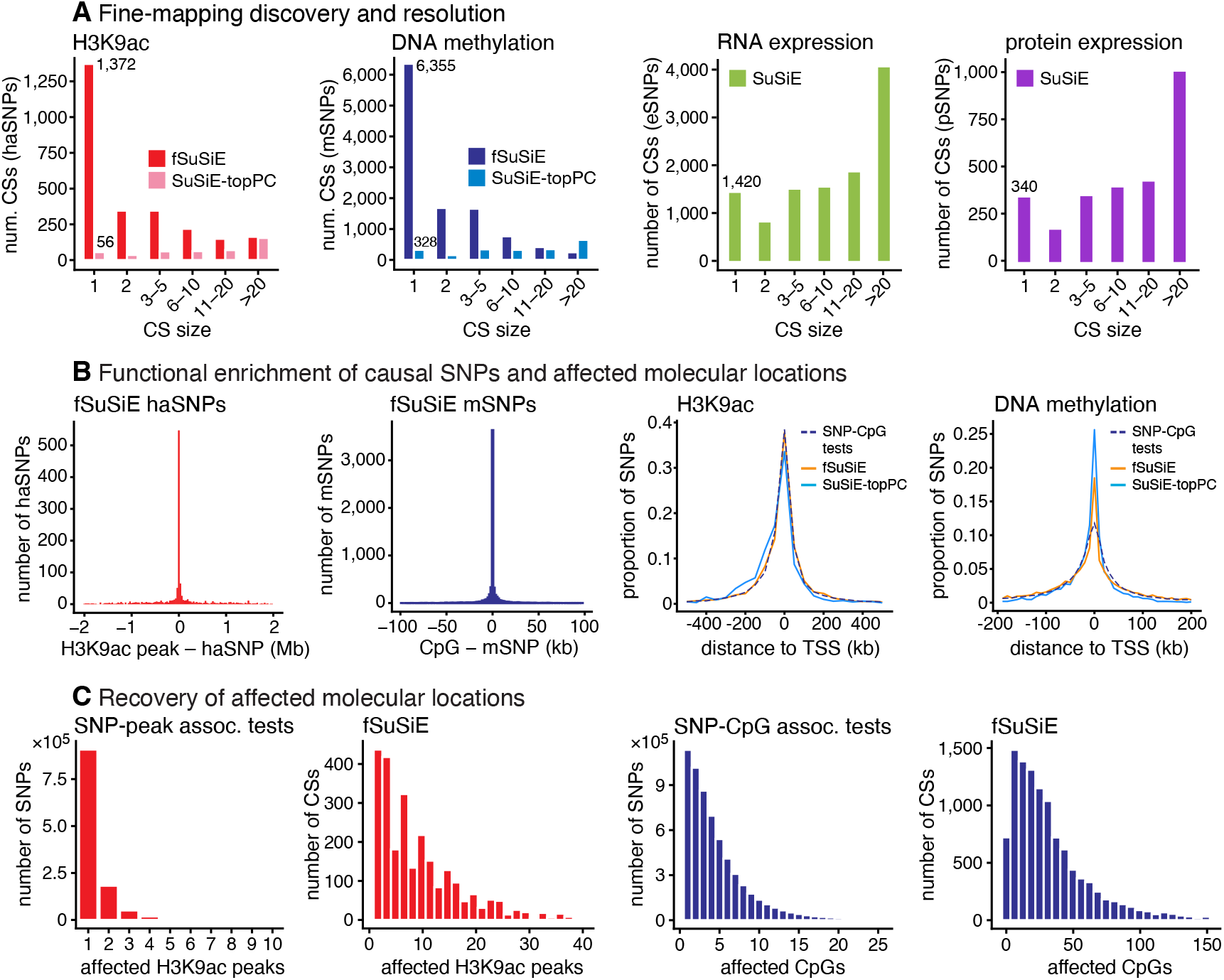
Fine-mapping of mSNPs (*n* = 636) and haSNPs (*n* = 592) in DLPFC. Panel A compares: the number of CSs returned by SuSiE-topPC, fSuSiE (for H3K9ac and DNA methylation) and SuSiE (for RNA and protein expression, also in DLPFC; *n* = 777 and 416, respectively); and the fine-mapping resolution (“CS size” is the number of SNPs included in the 95% CS). In Panel B, functional enrichment of the mSNPs and haSNPs is assessed by distance between fine-mapped SNP and the nearest affected molecular location (for single-variant CSs only), and by distance to the nearest gene transcription start site (TSS). Panel C compares the number of affected molecular locations per SNP from fSuSiE and the association tests. Notes: in C, a small fraction of SNPs and CSs (*<*2%) with a larger number of affected locations were not included in the histograms; in the distance-to-TSS plots, the SuSiE-topPC and fSuSiE SNP counts were weighted by the PIPs; the TSSs are obtained from the annotated gene transcripts in the GENCODE combined Ensembl/HAVANA database [79]; all the CS statistics do not double-count CSs that overlap in different (overlapping) TADs. See Supplementary Fig. 20 for additional supporting results.

Fig. 5 summarizes the genome-wide results from applying fSuSiE to all the methylation and H3K9ac data. Consistent with the increases in power shown in the simulations, fSuSiE identified many more putative causal SNPs than SuSiE-topPC, and at a much higher resolution; for example, fSuSiE identified 1,372 single-variant CSs for H3K9ac and 6,355 single-variant CSs for DNA methylation, compared to just 56 and 328, respectively, from SuSiE-topPC (Fig. 5A; see also Supplementary Fig. 20A, B). Among the SNPs in single-variant CSs, 55 were identified as affecting both histone acetylation and methylation. Furthermore, 50 of the mSNPs and 27 of the haSNPs affect transcript levels of one or more genes. These observations suggest potential coordinated cis-regulatory effects (Supplementary Fig. 21). fSuSiE also produced a much higher proportion of single-variant CSs compared to SuSiE fine-mapping analysis of RNA and protein expression (Fig. 5A), again highlighting the benefits of joint analysis of multiple traits [33]. (Although the increase in single-variant CSs outputted by fSuSiE here is consistent with results in the simulations, and consistent with the expectation that joint analysis will improve fine-mapping resolution, it is also prudent to point out that fSuSiE makes additional modeling assumptions about the molecular trait. Large deviations from these assumptions may cause fSuSiE to be overcondident in its inferences, and potentially produce CSs that are too small. In particular, additional factors such as batch effects could induce correlations between molecular locations if they are not included in the fSuSiE model; see Methods for details.)

The many more additional haSNPs and mSNPs discovered by fSuSiE tend to be located close to the inferred affected peaks/CpGs, and remain concentrated near gene TSSs (Fig. 5B). This provides evidence that the additional effects detected are mostly real signals rather than false positives. More generally, enrichments of haSNPs and mSNPs in predefined functional annotations [80, 81] were largely consistent between fSuSiE and SuSiE-topPC, suggesting that the additional discoveries of fSuSiE maintain similar functional characteristics (Supplementary Fig. 22). One exception was the stronger enrichments of SuSiE-topPC mSNPs in regions involved more directly in gene transcription (TSS, 5’ UTR, promoters and coding regions). This may be because the strongest methylation signals (the top methylation PC) tend to occur near regions of direct transcriptional regulation. In comparison, fSuSiE, by looking beyond the strongest methylation signals, provides a more balanced picture of mSNPs involved in both direct and indirect gene regulation.

As a fine-mapping method, fSuSiE provides a much more parsimonious explanation for observed associations than does conventional (univariate or “SNP-by-SNP”) QTL association testing (Fig. 5C). Conventional QTL testing results in large numbers of SNPs, many in high LD with one other, being associated with a (usually) small number of peaks/CpGs; in contrast, fSuSiE identifies a much smaller number of potentially-causal SNPs associated with (usually) a larger number of peaks/CpGs. In other words, fSuSiE finds many more changes in methylation and histone acetylation that span large parts the fine-mapping regions; some mSNPs are estimated to affect methylation levels at more than 100 CpGs). The vast majority of mSNPs and haSNPs identified by fSuSiE are associated with methylation and histone acetylation at locations very close to the SNP; for 96% mSNPs and 54% haSNPs, the nearest affected location is within 100 kb (Fig. 5B). Additionally, 26.8% of haSNP CSs contain at least one SNP that is within the MACS2-called interval of the affected H3K9ac peak. These results agree with the general expectation that the variants affecting chromatin are often within the regulated chromatin region [82].

### fSuSiE reveals epigenetic regulatory mechanisms colocalizing with Alzheimer’s disease risk loci

Alzheimer’s disease (AD) is a complex neurodegenerative disorder with substantial genetic heritability. Recent large-scale genome-wide association studies have identified hundreds of AD risk loci [83, 84], but the molecular mechanisms underlying most genetic associations remain unclear. Since GWAS variants typically affect gene regulation rather than protein coding sequences, integration of molecular QTL data with GWAS through colocalization analysis offers a powerful approach to identify regulatory mechanisms and target genes underlying disease associations. To demonstrate this, we performed colocalization analysis using COLOC (version 5) [85] between our fSuSiE fine-mapping results for methylation and histone acetylation, and AD GWAS summary statistics from two large-scale meta-analyses [83, 84].

This genome-wide colocalization analysis identified 18 loci colocalizing with AD risk variants in 95% colocalization CSs. For illustration, we focus on 6 of these loci (3 methylation and 3 histone acetylation) with AD variants reaching genome-wide significance (Fig. 6, Supplementary Figures 23 and 24). A targeted SuSiE+COLOC comparison on the fSuSiE-nominated regions recovered 5 of the 18 loci with broadly consistent evidence; the fraction of the remaining 13 fSuSiE-only loci reaching AD *p <* 5 × 10^−8^ (53.8%) is comparable to the shared set (60%), so the fSuSiE-only loci are supported by AD evidence of similar strength. These colocalized variants suggest potential regulatory targets for AD-associated epigenetic changes. In an attempt to pinpoint the specific genes regulated by these epigenetic changes, we integrated our results with eQTL data from the same ROSMAP DLPFC samples. This multi-modal integration provides substantial advantages over eQTL analysis alone: while eQTL colocalization with GWAS can identify genes affected by disease variants, the association itself does not provide mechanistic insights into how the disease variants may operate through regulation. To illustrate how integrating epigenomic QTL data can help address this, we show results for two well-studied AD risk loci, near genes *CASS4* and *CR1*/*CR2*), where the fSuSiE fine-mapping results provide mechanistic insights into disease-associated genetic variants.

**Fig. 6.**
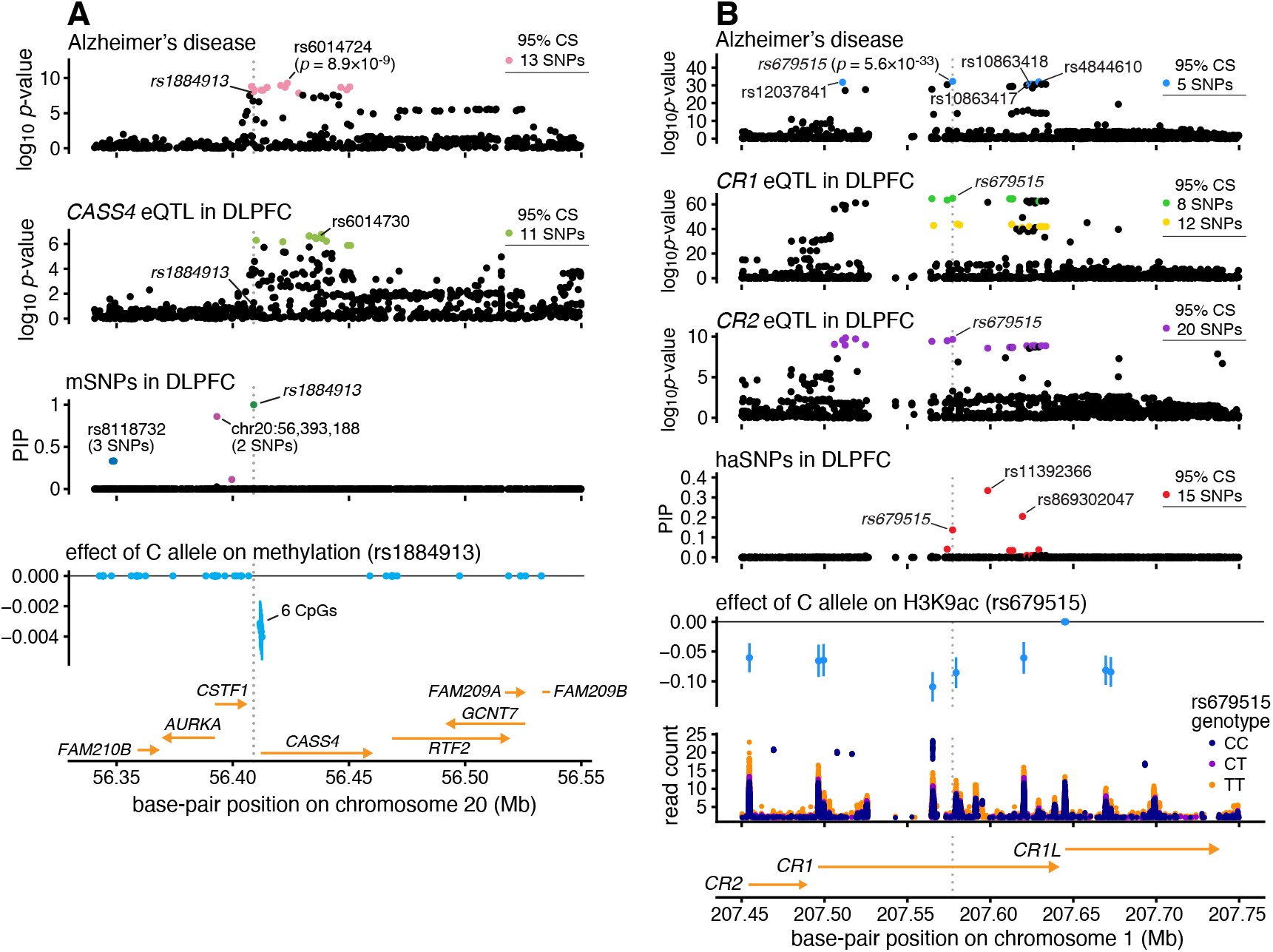
Results of methylation fine-mapping at the *CASS4* AD locus (A) and H3K9ac fine-mapping at the *CR1*/*CR2* AD locus (B). *AD (top) plots:* association *p-*values (two-sided *t*-test, *n* = 455,258 or 788,989) from Alzheimer’s disease (AD) GWAS [83, 84], and 95% CS from SuSiE fine-mapping using the AD GWAS summary statistics [24, 25]. *eQTL plots:* association *p-*values (two-sided *t-*test, *n* = 777) from eQTL analysis and 95% CS from SuSiE fine-mapping. The top SNPs for AD and gene expression are labeled. *PIP plots:* PIPs obtained from fSuSiE fine-mapping of methylation or histone acetylation. *Effect plots:* SNP effects on methylation or histone acetylation estimated by fSuSiE. CpGs or peaks unaffected by the SNP are drawn on the *y* = 0 line. Error bars depict 95% credible bands (*n* = 636). For each CS, the sentinel SNP (the SNP with the highest PIP in the CS) is labeled. If the CS contains more than 1 SNP, the total number of SNPs in the 95% CS is given. For the gene trascript annotations, the arrow indicates the direction of transcription (source: GENCODE Ensembl/HAVANA [79]). *Read count plot (B only):* the “raw” ChIP-seq peaks (average read counts) for each genotype at SNP rs679515. Average read counts less than 2 are not shown.

*CASS4* is a potential AD risk gene supported by association analyses [83, 86, 87] and colocalization of AD SNPs with *CASS4* expression SNPs [87, 88]. Previous colocalization and fine-mapping analyses have suggested rs6014724 and rs17462136 in the 5’ UTR of *CASS4* as putative causal variants [88]. The fSuSiE methylation fine-mapping suggests another potential causal variant: rs1884913 (PIP *>* 0.99, MAF = 23.2%), an mSNP that lies within the CS for AD, and affects methylation levels at 6 CpGs near the *CASS4* TSS (Fig. 6A, Supplementary Fig. 19). Notably, rs1884913 shows moderate correlation with the previously reported variant rs6014724 (*r* = 0.51, *r*^2^ = 0.26) but high correlation with rs17462136 (*r* = 0.94, *r*^2^ = 0.88). This finding, supported by convergent evidence from multiple molecular QTL analyses, suggests that rs1884913 as a potential causal variant may contribute to understanding regulatory mechanisms underlying disease-associated genetic variants, and illustrates how multivariate functional analysis of molecular phenotypes may achieve higher fine-mapping resolution than univariate analysis of complex disease phenotypes by distinguishing between functionally distinct variants (rs17462136 and rs1884913 are separated by 11.3 kb) that remain indistinguishable using GWAS data alone due to high linkage disequilibrium.

The *CR1*/*CR2* AD risk locus [87, 89, 90] has also been intensely investigated [91–94], including in a recent proteome-wide association study that suggested both *CR1* and *CR2* as causal proteins [95]. However, none of these studies have pinpointed the causal variant. Our SuSiE and fSuSiE analyses suggest that the top association for AD, rs679515 (*p* = 5.6 × 10^−33^, MAF = 20.0%) is a solid candidate causal variant: first, rs679515 is the only SNP that overlaps CSs for both *CR1* expression and *CR2* expression; second, this SNP is one of the top SNPs in a CS for H3K9ac, with PIP = 0.14 (Fig. 6B, Supplementary Fig. 19). Furthermore, fSuSiE results suggest rs679515 has moderately long-range effects on histone acetylation, including at H3K9ac peaks near the TSSs of both *CR1* and *CR2* (Fig. 6B). Therefore, the fSuSiE H3K9ac results not only nominates a potential causal variant (rs679515), but also implicate both *CR1* and *CR2* via histone modification associations. (This result is supported by the multi-trait colocalization analysis in another FunGen-xQTL study [96].) We caution that the direction of causality is ambiguous in this example; fSuSiE cannot distinguish between these two scenarios: (i) the SNP affects regulatory activity near both genes; (ii) the SNP affects regulatory activity near one gene which in turn alters activity near the other gene [82].

## Discussion

We have introduced fSuSiE, a method that extends SuSiE to high-dimensional, spatially structured traits by borrowing ideas from functional regression. In simulated and real data sets, we demonstrated the benefits of fSuSiE for fine-mapping high-dimensional molecular traits; most notably, fSuSiE is the only method that can simultaneously estimate the genetic variants affecting the molecular trait and the specific changes to the molecular trait that are produced by the genetic variants. Although fSuSiE is more computationally demanding than conventional QTL association analysis, or than fine-mapping of a univariate trait (e.g., SuSiE-topPC), as we have shown here it remains practical for data sets containing hundreds of samples, thousands of molecular locations, and thousands of SNPs.

The molecular data sets we considered here consist of measurements made at predefined genomic locations (CpGs or peaks called by MACS). However, fSuSiE does not require predefined molecular features; for sequencing data (e.g., RNA-seq, ChIP-seq, WGBS), fSuSiE could instead be applied directly to the base-pair-level sequencing data (or data combined into small bins), in which case the relevant molecular features would be inferred from the fSuSiE effect estimates and credible bands. Applying fSuSiE to high-resolution sequencing data has the potential to quantify gene regulatory effects in greater detail and discover additional gene regulatory effects in regions outside the called peaks, opening new avenues for analyzing and integrating multiome data. In fact, beyond the ROSMAP methylation and histone modification analyses presented here, we have deployed fSuSiE across diverse molecular contexts within the FunGen-xQTL consortium, including integration with methylation data from multiple cohorts and analysis of single-nucleus ATAC-seq data from six major brain cell types for chromatin accessibility using recently published data on ROSMAP individuals [97].

fSuSiE is also a very flexible model framework, and includes additional modeling features we did not highlight in this paper. For example, one could incorporate functional annotations to guide discovery of the causal variants and affected molecular features. Second, fSuSiE is not limited to using wavelet-based functional regression; our implementation also includes hidden Markov model (HMM) alternatives that, while computationally slower, can sometimes offer more refined results for certain data types, and other functional regression approaches might be better suited to certain types of molecular data. A key advantage of embedding our approach within the SuSiE framework is the seamless integration with numerous available downstream analytical methods based on SuSiE. This design choice makes fSuSiE readily available for data integration pipelines, as demonstrated through our colocalization analysis using SuSiE-based COLOC methods [85] in Alzheimer’s disease GWAS applications. Additionally, fSuSiE-estimated effect sizes can be directly incorporated into transcriptome-wide association studies (TWAS) to combine multiple genetic regulatory effects across molecular profile regions with GWAS signals. Note that for other fine-mapping problems in which there are a small number of traits that are not necessarily spatially structured, mvSuSiE [33] remains a good option. (See Supplementary Fig. 17 for an illustration of how univariate SuSiE, mvSuSiE, and fSuSiE behave on a single locus with a spatially structured effect.)

Our work has several limitations representing directions for future research. First, the current methods do not handle missing data: the molecular trait needs to be measured at every location in every sample. When some measurements are missing, they could potentially be imputed provided that the imputation is accurate and/or there are not too many missing measurements. Second, the current framework requires analyzing one molecular context at a time. Future work includes developing joint modeling approaches for multiple molecular contexts to finemap shared effects without requiring separate colocalization analyses. This would be particularly valuable for multiome data where both single-nucleus RNA-seq and single-nucleus ATAC-seq data are available, enabling simultaneous analysis of gene expression and chromatin accessibility as spatially structured molecular profiles. Third, while fSuSiE can handle both normalized and count-based data, our applications focused on normalized molecular phenotypes. Although our method and software implementation includes count-based models specifically designed for high-throughput sequencing data that could be beneficial in other applications, we did not apply these count-based alternatives in the current work.

A final limitation we mention is that fSuSiE relies on individual-level data. fSuSiE could, in principle, be adapted to summary statistics (association test statistics and an LD matrix), analogous to SuSiE-RSS and mvSuSiE-RSS [25, 33] (see the Supplementary Note). However, it is not clear that this would be practical or desirable for high-dimensional molecular traits. Consider that standard summary-statistics sharing for spatially structured data first collapses the signal into called peaks or regions and then reports per-feature summaries, which loses resolution by construction. fSuSiE works directly on the sequencing coverage or on small fixed-width bins (e.g., 50 bp); *T* therefore scales with assay resolution, from the ~850K CpGs of the Illumina EPIC array to the ~28 million CpGs in human whole-genome bisulfite sequencing, pushing *T* into the thousands per window without peak calling. The fSuSiE output is not a *J*× *T* matrix but a compact fine-mapping object that, as demonstrated here, can be consumed directly by SuSiE-based COLOC [85] and related pipelines. We therefore recommend that molecular-QTL studies run fSuSiE on their individual-level data and share the fine-mapping objects for downstream integration.

The fSuSiE methods are implemented in an R package, fsusieR, available on GitHub and distributed via the MIT Open Source license. The R package includes detailed documentation, examples, and guidance on applying the methods in practice.

## Methods

### fSuSiE

#### Wavelet-based functional model

Consider a data set consisting of *N* samples of a molecular trait measured at *T* locations. The measurements are stored in an *N*×*T* matrix **Y**, in which *i*∈{1, …, *N*} indexes samples and *t* ∈ {1, …, *T*} indexes locations. Let ***x*** = (*x*_1_, …, *x*_*N*_)^⊺^ be a covariate of interest (e.g., a genotype) measured in the *N* samples.

We begin with a standard multivariate linear regression model of **Y** given ***x***,

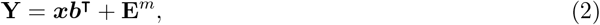

where ***b*** = (*b*_1_, …, *b*_*T*_)^⊺^ is a column vector of regression coefficients (“effects”) to be estimated, and **E**^*m*^ is an *N* ×*T* matrix of residuals (which we will assume are normally distributed). The *m* in the superscript here refers to the fact that the residuals are defined in “measurement space” (as opposed to “wavelet space”, which will be denoted using the *w* superscript).

Transforming **Y** to the wavelet space involves a simple linear transformation, **D** = **YW**^−1^, where **W** is the *T* ×*T* DWT matrix. The resulting transformed data, **D**, also an *N*× *T* matrix, contains the empirical wavelet coefficients (WCs). The DWT matrix **W** is an orthogonal matrix, which is very convenient; for example, statistical quantities on the measurement scale are easily recovered after they have been estimated on the wavelet scale. The orthogonality of **W** also means that **W**^−1^ = **W**^⊺^, and therefore the transformed data can be written as **D** = **YW**^⊺^. (See [63, 75, 98–103] and the Supplementary Note for background on statistical modeling using wavelets.) Applying **D** = **YW**^⊺^ to (2) results in a wavelet space version of the model,

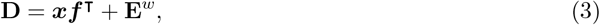

where ***f*** = **W*b*** is the vector of effects in the wavelet space, and **E**^*w*^ = **E**^*m*^**W**^⊺^ is an *N* ×*T* matrix of residuals in the wavelet space.

Next we extend (2) to a multiple linear regression model in which the molecular trait is modeled as a linear combination of *J* covariates of interest (e.g., genotypes measured at *J* SNPs):

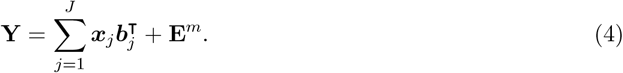

The corresponding multiple regression model in wavelet space is

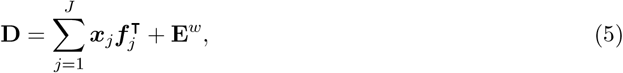

in which ***f***_*j*_ = **W**^⊺^***b***_*j*_. We often write (4) and (5) more compactly as

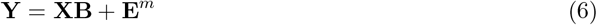

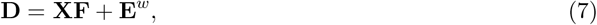

where **X** is the *N*× *J* matrix containing *N* observations about the *J* covariates of interest, **B** is the *J*× *T* matrix of unknown effects, and **F** = **BW**^⊺^ is the *J* ×*T* matrix of unknown effects in the wavelet space. fSusiE is developed based on (7). (The measurement-space model (6) is implied but never directly used in fSuSiE.) In expressions below, we denote elements of **X, Y, B, F, E**^*m*^ and **E**^*w*^ by *x*_*ij*_, *y*_*it*_, *b*_*jt*_, *f*_*jt*_, 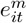and 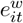, respectively. To simplify notation in the equations below, we drop the “*w*” superscript from **E**^*w*^ when it is clear from the context that these are residuals in the wavelet space.

#### Independence of residuals in wavelet space

fSuSiE follows the common assumption in wavelet regression methods that the residuals in wavelet space 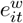 are independent across *t*; that is, **E**^*w*^ is multivariate normal with mean zero and diagonal covariance matrix **∑**, with diagonal entries 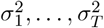. Allowing the variances 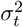 to vary by *t* is important for accommodating non-stationary signals, which frequently occur in practice [63, 74].

The independence of the residuals 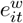 across indices *t*—i.e., across different wavelet scales and locations (see below)—can be motivated by the wavelets’ decorrelating effect on spatially structured signals, often referred to as the “whitening property” of the DWT. (See Fig. 1 in [63], Fig. 4.1 in [75], Figures 1 and 2 in [74], and https://stephenslab.github.io/fsusie-experiments/whitening_demo.html for illustrations of the whitening effect.) It is important to recognize that independence of the residuals 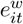 in the wavelet space does *not* imply independence of the residuals 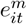 in the measurement space; indeed, assuming independence of the residuals in measurement space would be inappropriate due to the spatial structure. See [50, 63] for additional discussion.

#### Multiple wavelet regression model with an intercept and additional covariates

In practice, we augment the multiple wavelet regression model (7) to include an intercept, and we often include additional covariates (e.g., sex, age, batch effects, genotype PCs):

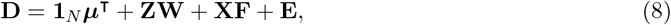

where **1**_*N*_ is a vector of ones of length *N*, **Z** is an *N*× *n*_*z*_ matrix containing data about *n*_*z*_ ≥ 0 additional covariates, ***µ*** = (*µ*_1_, …, *µ*_*T*_)^⊺^ is the (unknown) intercept, and **W** is the *n*_*z*_ ×*T* matrix of (unknown) coefficients corresponding to the additional covariates **Z**. To simplify presentation, we store the intercept ***µ*** in the first row **B**, and set the first column of **Z** to be all ones, so the same model can be written more simply as

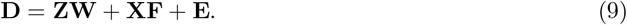

This model can be reduced to the previous model (7) if we remove the linear effects of **Z** on **X** and **D** by making the substitutions 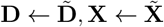, in which we define

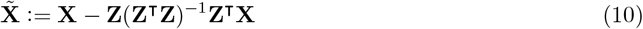

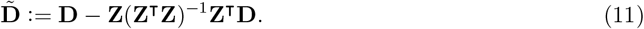

This is an extension the reduction used for univariate linear regression [115]. Note that we have sometimes found it more convenient to remove the linear effects of **Z** from **Y** instead of from **D**, then compute **D** = **YW**, but this is equivalent to (11) because **W** is orthogonal.

#### Standardization

In practice, we scale each column of **Y** to have unit variance before computing the wavelet transform (this is often called “standardization”). This simplifies modeling because it allows us to reasonably assume that the wavelet coefficients have homoskedastic residual variances; consider that 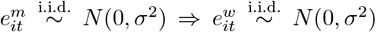. Therefore, in fSuSiE we make the additional modeling assumption that the residual variances do not depend on location; 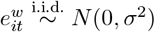.

Also, although not necessary, in practice we also standardize 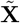 before running fSuSiE. This simplifies many of the underlying statistical computations in fSuSiE. Standardizing 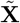 is common practice in QTL mapping analyses, and can be justified in some settings [104] [116]. However, standardizizing 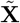 may also lead to issues with low-frequency SNPs; see “Disclaimer on including genetic variants at low minor allele frequencies” below.

#### Single function regression model

The “single function regression” (SFR) model is the functional counterpart of the “single effect regression” (SER) model [24]. The SFR model is a multiple wavelet regression model (7) with the constraint that exactly one of the covariates *j*∈ {1, …, *J*} has a nonzero effect on the wavelet coefficients:

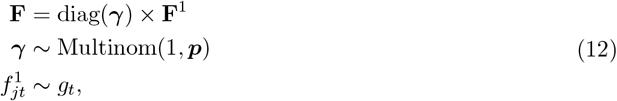

where ***γ*** = (*γ*_1_, …, *γ*_*J*_)^⊺^, **F**^1^ is a *J* ×*T* matrix with entries 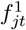, Multinom(*n*, ***p***) denotes the multinomial distribution with sample size *n* and multinomial probabilities ***p*** = (*p*_1_, …, *p*_*J*_), *g*_*t*_ denotes the prior distribution for the effects at location *t* (specified in greater detail below), and diag(***x***) denotes the *N*× *N* diagonal matrix in which the diagonal elements are given by the vector ***x*** = (*x*_1_, …, *x*_*N*_).

Defined in this way, the SFR model has the following properties: (i) ***γ*** is a binary vector in which exactly one of the elements is one and the rest are zeros; (ii) at most one of the rows of the **F** contains nonzero values; and (iii) the 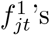 ‘s are *a priori* independent across locations *t* (again motivated by the “whitening” property of the wavelet transform).

Elements of ***p*** are the prior probabilities that each covariate has a nonzero effect on the wavelet coefficients **D**. Unless otherwise stated, the prior probabilities are the same for all covariates (SNPs); *p*_*j*_ = 1*/J, j* = 1, …, *J*.

#### fSuSiE model

The fSuSiE model extends the SFR model to allow at most *R* ≥ 1 nonzero effects. It is the functional counterpart to the Sum of Single Effects (SuSiE) model [24], so we call it the “functional Sum of Single Effects model” (fSuSiE) model. The fSuSiE model is the wavelet regression model (7) together with the following prior on **F**:

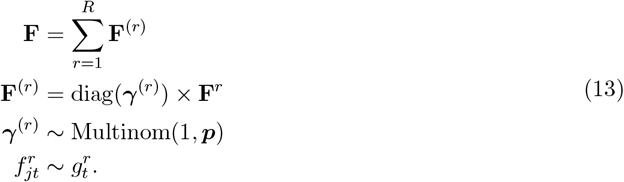

Since each binary vector 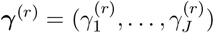 contains exactly one 1, by contruction at most *R* rows of **F** contain nonzeros. The fSuSiE model reduces to the SFR model when *R* = 1. Note that this model allows for a different prior on the effects 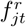 for each *r*; these priors are automatically adapted to the data using the algorithms described below.

#### Scale-dependent mixture priors for fSuSiE

Above, we did not specify the exact form of the priors *g*_*t*_ and 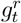 in the SFR and fSuSiE models. To define a prior, we expand on a detail that was previously hidden: the *t* index represents both a wavelet scale and location. Our convention is to use *s* to index scales and *l* to index locations.

Previous wavelet regression approaches have exploited the scale-dependent sparsity of the wavelet coefficients [101, 102] [117, 118]. One of these approaches used a spike-and-slab prior [117, 118]. Building on this spike-and-slab prior as well as our recent work on adaptive shrinkage methods [65, 104] [119], we extend this scale-dependent spike-and-slab prior to a scale-dependent mixture prior, which we refer to as the “shrinkage-per-scale” (SPS) prior:

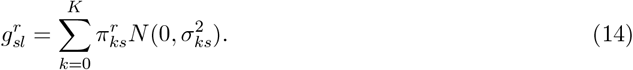

Here, the 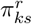 denote the mixture weights 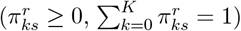 for each single function *r* = 1, …, *R*, and the 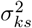 denote the variances of the mixture components. The number of mixture components, *K*, and the component variances 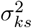 are fixed, user-specified quantities, whereas the mixture weights 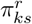 are estimated. Note that this prior depends on the scale *s* but not on the location *l*.

The component variances 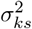 are assumed to be increasing 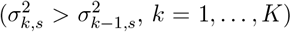, and the first component is assumed to be a Dirac “delta” mass at zero, denoted here by a normal distribution with zero variance, *N* (0, 0). We use procedures similar to those implemented in the ashr R package [104] to automatically determine a suitable *K* as well as the components variances 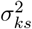.

We also consider a special case of the SPS prior in which the mixture proportions *π*_*ks*_ and variances 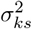 do not depend on the scale, *s* [65]; that is, 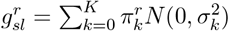. We call this the “independent shrinkage” (IS) prior. While clearly less flexible than the SPS prior, the IS prior reduces the number of parameters to estimate and reduces overall computational effort, and therefore is sometimes more convenient in practice. We empirically assess the benefits of the SPS and IS priors in simulations.

#### Posterior statistics

Here we define the key posterior quantities used in an fSuSiE analysis (Fig. 1). We do not explain here how these quantities are computed; these details are given in the Supplementary Note.

As we briefly explained in the Methods overview, we have three main inference aims:

- *Variable selection:* identify the causal SNPs.
- *Feature annotation:* identify the molecular features and locations that are affected by one or more SNPs.
- *Feature selection:* identify the molecular features and locations that are affected by a given causal SNP.

For variable selection, we compute *credible sets* (CSs): a CS is defined as a subset of {1, …, *J*} containing an effect SNP with high probability [20, 24]. More precisely, a level-*ρ* CS is defined as a set of SNPs that is as small as possible such that it has probability at least *ρ* of containing an effect SNP (a row of **F** containing at least one non-zero). The number of CSs should reflect the number of causal SNPs, and the size of a CS should reflect the number of plausible candidate effects SNPs. We calculate CSs as described in [24].

To determine which SNPs within a CS are the strongest candidates for being an effect SNP, we compute a *posterior inclusion probability* (PIP) for each SNP. The PIP for SNP *j* is defined as the posterior probability that at least one of the entries in the *j*th row of **F** is nonzero:

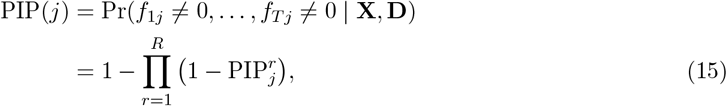

in which 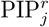 denotes the posterior probability that the *r*th single effect is nonzero for SNP *j*,

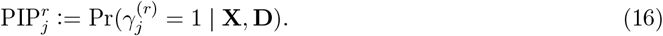

See 2D for an illustration of PIPs and CSs.

The estimates of the SNPs on the molecular features in wavelet space are given by the posterior mean of **F**, 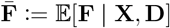. The SNP effects in measurement space are given by the posterior mean of **B**, 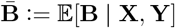. By elementary properties of expectations, this is simply

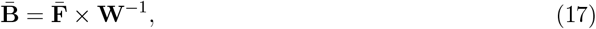

where **W**^−1^ denotes the inverse wavelet transform.

To identify affected features and locations, we compute 1 − *α* pointwise Bayesian credible intervals [105, 106] for elements *f*_*jt*_ and *b*_*jt*_ at selected SNPs *j*. We define “affected” as those elements in which the interval does not include zero. We refer to these intervals as “credible bands.” See Fig. 2E giving an example of the posterior means and the credible bands of *b*_*jt*_ for all locations (CpGs) *t* = 1, …, *T* and for all sentinel SNPs *j*. The affected locations *t* are defined as the locations in which at least one of the credible bands for the sentinel SNPs at *t* does not contain a zero.

#### Disclaimer on including genetic variants at low minor allele frequencies

In examining the results of the fSuSiE analyses, we found that many of the single-variant CSs (i.e., the predicted mSNPs or haSNPs) affected molecular trait locations that were far away (*>*100 kb) from the predicted mSNP or haSNP. These long-range interactions were heavily concentrated at lower minor allele frequencies (MAFs); in particular, we observed a strong excess of SNPs with MAF *<* 5% that these long-range interactions. This suggests that the fSuSiE results for low-frequency SNPs are unreliable. Therefore, *post hoc* we excluded from the fSuSiE results all CSs for which the sentinel SNP had a MAF *<* 5%. For future fSuSiE analyses, the simple recommendation would to exclude SNPs below a certain MAF threshold as a data preprocessing step before running fSuSiE.

More generally, this suggests an issue with low-frequency genetic variants, particularly at smaller sample sizes (the sample size was about 600 in the data sets we analyzed). Although not confirmed empirically, we suspect this is caused by one of two issues (or a combination of both).

The first issue is that fSuSiE is based on Gaussian model of the trait—specifically the residuals are assumed to be normally distributed—and it is possible that low-MAF SNPs are more sensitive to the violations of this assumption, particularly at smaller sample sizes.

The second issue is standardizing 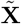, which was done mainly to simplify implementation (see “Standardization” above). It should be noted that the genotypes are commonly standardized in QTL mapping and fine-mapping analyses—either explicitly or implicitly—and is perhaps even more commonly done than not. Standardizing 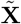 makes the implicit assumption that SNPs at low MAFs have larger effects than SNPs at larger MAFs *a priori* (specifically, the prior effect size is proportional to 1*/*(MAF × (1 −MAF)) [116]). Even if this assumption is not valid, it is not expected to have a large impact when the sample size is large and/or all SNPs have reasonably high MAF. However, we suspect that this assumption was more harmful in our analyses because: (i) the sample sizes are smaller than most fine-mapping analyses of complex traits; (ii) the fact that fSuSiE is modeling the effects of each SNP at many (e.g., thousands) molecular trait locations. If confirmed, the remedy would be to not standardize 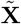.

#### Outline of an fSuSiE analysis

Briefly, the minimal requirements for performing an fSuSiE analysis are:

i. An *N* × *T* matrix, 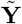, containing *N* molecular trait measurements at *T* locations after the linear effects of selected covariates are removed. The specific steps that were taken to prepare 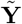 for the molecular data sets analyzed in this paper are detailed below.
ii. An *N* × *J* matrix, 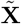, containing the genotype information for the *J* SNPs to be fine-mapped after removing the linear effects of the selected covariates.
iii. *R*^max^, an upper limit on *R*, the number of single functions in the fSuSiE model. Unless specifically mentioned, we set *R*^max^ = 10.
iv. The prior for the wavelet effects. In this paper, we considered two priors: the independent shrinkage (IS) prior and the shrinkage-per-scale (SPS) prior (see “Scale-dependent mixture priors for fSuSiE” for explanations). We used the IS prior unless otherwise mentioned.

An additional input is optional but recommended:

V) The positions (e.g., base-pair positions) *ϕ*_1_, …, *ϕ*_*T*_ ∈ **R** corresponding to the locations 1, …, *T*. If not specified, the positions are assumed to be evenly spaced. Molecular traits are typically not measured at evenly spaced positions, so providing this information will produce more accurate results.

Beyond this, the fSuSiE software includes many other settings and tuning parameters which may be adjusted as needed.

The basic steps of an fSuSiE analysis are as follows:

1. *Compute the wavelet coefficients*. Compute the WCs, 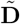, from the molecular trait data, 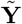. For computational efficiency, 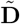 is obtained using the standard discrete wavelet transform (DWT). (In later steps, a second set of WCs are computed using the translation-invariant wavelet transform to improve accuracy.) For all the results presented in this paper, we used the (undecimated) wavelet transform with Daubechies least-asymmetric orthonormal compactly supported wavelets and with 10 vanishing moments (see Chapter 2 of [75]). Note that the fSuSiE software currently supports any wavelet transform that is implemented in the wavethresh R package [75].
2. *Search for a good upper limit on R (optional). R* determines the number of single functions and, correspondingly, the number of causal SNPs. If *R* is too small, fSuSiE may miss some causal SNPs; on the other hand, if *R* is too large, fSuSiE may take a long time to run. We have implemented an *ad hoc* procedure to find a reasonable initial estimate of *R*. By default, this procedure starts by fitting an fSuSiE model with *R* = 3. If all *R* = 3 single functions are kept, *R* is increased by one. This procedure iterates until one or more single functions are pruned, or if the upper limit, *R*^max^, is reached.
3. *Fit fSuSiE model*. We use an iterative algorithm, described below, to fit the fSuSiE model to the data 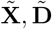. This includes estimating the priors 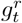 and the residual variance *σ*^2^. During estimation of the priors, some single functions may be pruned, and therefore *R* is an upper bound on the final number of single functions (and CSs).
4. *Compute SNP-level posterior quantities: CSs and PIPs*. Compute CSs and PIPs from the fSuSiE model fitted in the previous step.
5. *Filter credible sets (optional)*. One may filter out the CSs with low “purity” (purity is defined as the smallest absolute correlation among all pairs of SNPs in the CS). This often improves quality or interpretability of the fine-mapping results.
6. *Compute posterior effect estimates and credible bands*. The final step is to compute posterior mean estimates of *b*_*jt*_, and corresponding 1 −*α* pointwise credible bands, *α* ∈ [0, 1], for each location *t* and sentinel SNP *j* (the SNP with the largest PIP in each CS). To address possible inaccuracies with the DWT (see Chapter 9 of [100]), we compute a new WC matrix 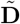 using the translation-invariant wavelet transform (TIWT), also known as the stationary wavelet transform. (In brief, the TIWT modifies the original DWT by applying it to shifted copies of the signal, then the WCs are averaged across the shifted copies.) Since the TIWT greatly increases computational effort, we use this TIWT only in this final step.

All these steps are implemented by the susiF function of the fsusieR R package.

### Molecular QTL simulations

#### Simulation of genotype data

The fine-mapping regions for the simulations were selected uniformly at random from 94 breast cancer loci on autosomal chromosomes reported in [120] (see Supplementary Table 1 of that paper). The median size of a fine-mapping reagion was about 1 Mb.

Similar to [121], we used sim1000G [67] to simulate genotypes of unrelated individuals based on the genotypes from the 1000 Genomes Phase 3 whole-genome sequencing [68]. First, we randomly selected a continent-of-origin label (EUR, AMR, AFR, EAS, SAS), then we simulated SNP genotypes using *N* = 100 individuals chosen uniformly at random from the 1000 Genomes samples with the selected continent-of-origin label.

Within the fine-mapping region, we kept all biallelic SNPs with minor allele frequencies (MAFs) of 5% or greater; that is, SNPs in which the minor allele was observed at least 10 times out of the 2*N* = 200 chromosomes).

For a single simulation, the genotype matrix, **X**, was a matrix with *N* = 100 rows (individuals) and *J* columns (SNPs), in which *J* ranged from approximation 1,500 to 4,000.

#### Simulation of molecular trait data—wavelet simulations

In these simulations, the molecular trait data were simulated from an fSuSiE model, and more precisely, a multiple wavelet regression model with an SPS prior for the causal SNP effects. Simulating the data involved the following steps: the effects of all causal SNPs *j* (on the wavelet scale) were simulated from an SPS prior, 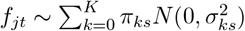, in which the first mixture component was always a “spike” at zero (*σ*_0*s*_ = 0); the effects of all non-causal SNPs *j* were set to zero, *f*_*jt*_ = 0; and then the effects in the measurement space were then obtained as **B** = **FW**^−1^. Similar to [101][117, 118], we simulated molecular trait data with different levels of smoothness: first, we drew a “smoothness parameter”, *µ*, uniformly at random between 0 and 1; then we set the *k* = 0 (“spike”) mixture component at each scale *s* to be *π*_0*s*_ = 1 − 2^−*µs*^. The intuition is that settings of *µ* closer to one produce “smoother” signals (larger effects at the largest scales), while settings of *µ* closer to zero produce effects of similar size across all scales.

Finally, we simulated the molecular trait data, **Y**, according to (6). The noise *e*_*it*_ was simulated in different ways. First, we considered i.i.d. noise, 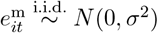. (Note that independent homoskedastic nose in the original space is the same as independent homoskedastic noise in the wavelet space because the wavelet transform is orthogonal; that is, simulating 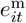 as i.i.d. normal is the same as simulating 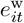 as i.i.d. normal.) Second, we simulated spatially structured noise 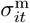using: (i) a Gaussian process (GP) [107] with a squared exponential kernel, 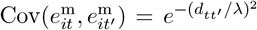 and *λ* = 20, where *d*_*tt*_*′* = |*t*− *t*^′^| denotes the distance between trait locations *t* and *t*^′^ (defined in this way, the distance between two adjacent locations was always equal to 1); or (ii) fractional Gaussian noise [108] with Hurst parameter *H* = 0.75. The GP introduced shorter-range dependencies among molecular trait locations, whereas the fractional Gaussian noise induced longer-range dependencies. In all simulations, the noise was rescaled to achieve the specified proportion of variance in **X** explained by **Y**.

#### Simulation of molecular trait data—WGBS block and WGBS decay simulations

In these simulations, we simulated methylation data from WGBS similar to [69]. The methylation profiles, **Y**, were simulated using the following logistic regression model:

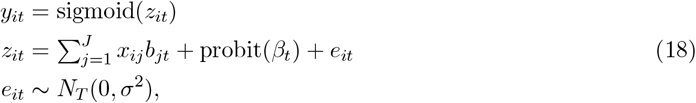

where *β*_*t*_ denotes a baseline methylation level (a “Beta value” [122]) simulated using OmicsSIMLA [123], sigmoid(*x*) = 1*/*(1 + *e*^−*x*^) denotes the sigmoid function, probit(*x*) = Φ^−1^(*x*) denotes the probit link function in which Φ(*x*) is the cumulative density function (CDF) of the standard normal distribution, *N*_*d*_(*x*; ***µ*, ∑**) denotes the multivariate normal distribution on ***x*** ∈ **R**^*d*^ with mean ***µ*** and covariance matrix **∑**, and the other notation was defined above. We transform the Beta values using the probit as this makes the data approximately homoskedastic with a Gaussian noise distribution [122]. The residual variance *σ*^2^ was chosen so that **X** explained a specified total variance in the unobserved **Z**.

The effects ***b***_*j*_ ∈ **R**^*T*^ of the causal SNPs *j* were simulated from a multivariate normal distribution, ***b***_*j*_ ∈ *N*_*T*_ (0, **C**), in which **C** is a *T* × *T* correlation matrix. (The effects of the non-causal SNPs *j* were set to zero, *b*_*jt*_ = 0.) The correlation matrix **C** determines the spatial structure of the methylation changes. We considered two different designs of **C** that resulted in two different sets of WGBS simulations, which we referred to as “WGBS block” and “WGBS decay” in the text.

##### WGBS block simulations

In a “WGBS block” simulation, the fine-mapping region was divided into 5 blocks with the same number of CpGs in each block. The SNP effects were the same within a block: *c*_*tt*_*′* = 1 if CpGs *t, t*^′^ were in the same block; otherwise, *c*_*tt*_*′* = 0.

##### WGBS decay simulations

In a “WGBS decay” simulation, the fine-mapping region was divided into 5 blocks of the same size (same number of CpGs). The SNP effects were similar within a block, but differed more as the CpGs were more distant within the block: *c*_*tt*_*′* = *ρ*^|*t*−*t*^*′*^|^ if CpGs *t, t*^′^ were in the same block, otherwise *c*_*tt*_*′* = 0, with *ρ* = 0.9.

### Methods compared in the simulations

#### fSuSiE

We ran two variants of fSuSiE: one using the IS prior, and another using the SPS prior. fSuSiE was applied as described above (“Outline of an fSuSiE analysis”) by calling the susiF() function from the fsusieR R package with the following settings: L = 20, which sets the upper limit on the number of CSs to X; `prior = “mixture_normal”` for the IS prior or `prior = “mixture_normal_per_scale”` for the SPS prior; and all other options were kept at their defaults.

For a given CpG, we defined *α*_max_ as the largest confidence level *α* for which the corresponding credible band at that CpG did not include zero. For a given *α*_max_ threshold, a CpG was defined as “affected” if *α*_max_ was less than this threshold.

#### SuSiE-topPC

The top PC, denoted here by ***y***, was obtained from the methylation data, **Y**, after centering the columns of **Y**. (Note that the columns of **Y** were not scaled to unit variance.) Then we ran function susie() from the susieR R package [24] on the data **X, *y*** with the following settings: `standardize = TRUE; L = X`, which sets the upper limit on the number of CSs to X; and all other options were kept at their defaults. Note that SuSiE-topPC was not used to identify affected CpGs.

#### SNP-CpG association tests

The SNP-CpG association tests were implemented using the `lm()` function in R. For a given *p-*value threshold, a CpG *t* was defined as “affected” if the (two-sided) *p-*value from the association test for that CpG was less than the threshold.

#### Additional simulations to assess single-variant CSs

In the mSNP fine-mapping and haSNP fine-mapping, fSuSiE reported a large number of CSs with 1 SNP, a surprisingly high number based on our previous experience with fine-mapping other quantitative traits such as RNA and protein expression. Furthermore, we found that some of these single-variant CSs identified a SNP that was in *very* high LD with another SNP that was not in the CS. We wondered whether fSuSiE might be incorrectly generating CSs that were too small and/or incorrectly identifying the causal SNP, and therefore we performed an additional experiment to check this. Specifically, we examined the extreme case of fine-mapping two SNPs, one of which is causal, and the genotypes of the two SNPs differ *in only a single individual*. This is the strongest LD one can observe between two SNPs without being in perfect LD (i.e., a correlation of 1).

In these simulations, we chose the causal SNP at random from European ancestry WGS samples in 1000 Genomes phase 3 (*n* = 503). The second SNP was identical to the first except for a single individual chosen at random; for this individual, we set *x*_2*i*_ ≠ *x*_1*i*_ ∈ {0, 1, 2}. SNP 1 affected molecular locations *t* = 20, …, 30, and had no effect on all other locations (in total, there were 128 locations, *t* = 1, …, 128). The effects of SNP 1, ***b*** = (*b*_1_, …, *b*_128_)^⊺^, were the same for all affected locations, and were varied from simulation to simulation from 0.01 to 1. The molecular trait for individual *i* was simulated as ***y***_*i*_ = *x*_1*i*_***b*** + ***e***_*i*_, with *e*_*it*_ i.i.d. standard normal. We performed 100 simulations at each setting of the effect, 0.01, 0.02, …, 1, for a total of 10,000 simulations.

We compared the results from SuSiE and fSuSiE in these simualtions. For SuSiE, we used as a univariate trait the most associated location; that is, the location *t* with the smallest association *p*-value when testing for association between the location *t* and SNP 1.

In each simulation, we recorded the CS configuration: SNP 1 only; SNP 2 only; or both SNPs 1 and 2. The results of these simulations are summarized in Supplementary Fig. 13. Note that both SuSiE and fSuSiE rarely returned a CS containing the wrong SNP only (SNP 2): out of the 10,000 simulations, SuSiE returned this result in just 8 simulations, and fSuSiE returned this result in just 36 simulations.

### Fine-mapping of molecular traits and Alzheimer’s disease

#### Data sources overview

We analyzed molecular trait data—DNA methylation and histone acetylation (H3K9ac)—in DLPFC donors from the ROSMAP cohort [62]. We also analyzed DLPFC RNA and protein expression data generated by the ADSP Functional Genomics Consortium xQTL Project (FunGen-xQTL) [109, 110]. SNP genotypes were obtained in the DLPFC donors by WGS [110]. Fine-mapping of AD risk loci was performed using data from two recent AD GWASs [83, 84].

#### WGS genotype data

Whole-genome sequencing data for ROSMAP subjects were obtained from the Alzheimer’s Disease Sequencing Project (ADSP) release 4 (R4), comprising a subset of the NIAGADS NG00067 dataset [110]. All sequencing data in ADSP R4 were centrally processed by the Genome Center for Alzheimer’s Disease (GCAD) using the variant calling pipeline and data management tool (VCPA), a standardized pipeline functionally equivalent to the CCDG/TOPMed workflow [111]. VCPA, implemented in Workflow Description Language (WDL) and optimized for the Amazon EC2 cloud environment, accepts sequencing data in FASTQ, BAM, or CRAM formats.

Sequence reads were aligned to GRC human genome assembly 38 (hg38) using BWA-MEM [124], followed by variant calling of single nucleotide variants (SNVs) and short insertion-deletions (indels) using GATK HaplotypeCaller [125], with joint calling workflows for improved alignment quality through local realignment of insertions/deletions and base quality score recalibration using GATK modules. Genotype-level quality control (QC) set each genotype to missing if the read depth (DP) was less than 10 or the genotype quality (GQ) score was less than 20. Variant-level QC flags were applied in the following order: variants in GATK low sequence quality tranches (lacking FILTER “PASS” value above 99.8% VQSR Tranche); monomorphic variants; variants with high missing rate; and variants with high depth. ABHet annotation estimated whether biallelic variants matched expected allelic ratios, with ideal heterozygous variants having values near 0.5. For downstream analysis, variants were retained if they were not assigned any of these QC flags: ABHet between 0.25 and 0.75; missing rate *<*5%; MAF ≥ 1%; and no significant deviation from Hardy-Weinberg equilibrium (*p* ≥ 10^−6^). Additional sample-level quality control steps included: verifying genetic sex; flagging duplicated and related individuals; outliers identified by heterozygosity levels and genotype call rates. The final data set contained approximately 30 million biallelic SNVs and 3.5 million indels.

To control for population stratification, PCA was performed on LD-pruned common variants using PLINK (MAF *>* 5%, *r*^2^ *<* 0.2). We retained the first 15 PCs for use in subsequent analyses.

#### Defining regions for cis-eQTL mapping and fine-mapping of molecular traits

Since methylation and H3K9ac lack gene-centric reference points, we adopted a unified approach using topologically associating domains (TADs) and their boundaries to define regions for QTL mapping and fine-mapping for all molecular traits analyzed. The TADs and their boundaries were derived from combined brain and blood Hi-C data [78]. In total, we defined 1,449 “extended TADs” for our analyses ranging in size from 1 to 17 Mb. For fine-mapping of methylation and H3K9ac, we used these extended TADs as the fine-mapping regions. For QTL mapping and fine-mapping of RNA and protein expression, the analysis region was defined as the extended TAD containing the gene, plus any additional part of the chromosome needed to include ±1 Mb from the gene’s transcription start and end sites.

#### RNA-seq data and analyses of RNA expression

The DLPFC bulk RNA-seq came in three different batches (*n* = 638, 252, 251). Total RNA (≥ 50*µg*) from DLPFC was used for library preparation via poly(A) selection (strand-specific dUTP protocol, *n* = 638) or rRNA depletion (KAPA Stranded RNA-Seq Kit with RiboErase, *n* = 252, RiboGold *n* = 251). Libraries were sequenced at 30–50 million paired-end reads per donor with exact depth varying by batch as described in [61]. Following standard QC via FastQC adapter trimming using fastp [126], reads were aligned to the human reference genome (hg38) using STAR [127] with WASP correction to reduce reference bias [128], with further quality control via Picard [129]. Gene-level RNA expression was quantified with RNA-SeQC [130], removing genes if over 20% samples had TPM expression level of 10% or less. Sample-level RNA QC was performed following from methods outlined by GTEx V8 [36] using three metrics to remove outliers: relative log expression; Mahalanobis distance to hierarchical clustering of samples; and D-statistics quantifying average correlation between pairs of samples. After these QC steps, a total of *n* = 777 samples were retained for subsequent analyses of RNA expression in DLPFC.

After these quality control steps, the transcript abundance (TPM) matrices were quantile-normalized. Technical factors (batch, RNA integrity number, post-mortem interval) and biological covariates (sex, age at death) were included as covariates in QTL analyses, along with the top 15 genotype PCs to account for population stratification. Additional hidden confounders included as covariates were estimated by PCA performed on the quantile-normalized RNA expression matrix, with the number of PCs (34) determined by the Marchenko-Pastur limit [131].

QTL mapping for each gene was performed using TensorQTL [112]. Fine-mapping for each gene was performed using SuSiE [24] after removing the linear effects of aforementioned covariates from both the phenotype and genotype matrices. We took an adaptive approach to determine *L*, the number of CSs, for each gene: starting with *L* = 5, we iteratively increased *L* by 2 if all the single effect regressions (SERs) in the fitted model had a prior variance greater than zero. We continued in this way until at least one SER had a prior variance of zero. This adaptive strategy increased detection of causal SNPs while avoiding an unnecessarily large setting of *L* (which increases computation). Using this adaptive approach, the largest setting of *L* for a gene was 12. We filtered the CSs returned by SuSiE in two ways: we removed a CS if the “purity” [24]—the smallest absolute correlation (Pearson’s *r*) among all pairs of SNPs in the CS—was less than 0.8; and we removed a CS if the sentinel SNP had a MAF less than 5%. (See below for the rationale for filtering based on MAF.)

#### Proteomic data and analyses of protein abundance

DLPFC protein abundance was quantified using selected reaction monitoring (SRM) proteomics [132–135]. Gray matter tissue from DLPFC was homogenized and proteins were extracted, followed by trypsin digestion. Targeted peptides were selected based on prior discovery proteomics experiments, and SRM assays were performed with manual inspection to ensure correct peak assignment and peak boundaries. After QC and filtering for proteins quantified in at least 50% of samples, *n* = 416 subjects with matched genotype and protein abundance data of 7,710 proteins were retained for association analysis and fine-mapping. Peptide relative abundances were (base-2) log-transformed, and centered at the median, then imputation of missing protein levels was performed using grouped empirical Bayes matrix factorization (gEBMF) [136]. QTL mapping and fine-mapping was performed as described for RNA expression (see above).

#### H3K9ac ChIP-seq data, association analysis and fine-mapping

H3K9ac ChIP-seq [113] was performed on DLPFC donors using a well validated H3K9ac antibody (Millipore #06-942). Approximately 50 mg of gray matter tissue was dissected, cross-linked with 1% formaldehyde, and sonicated prior to overnight immunoprecipitation. Purified DNA was used to construct libraries (including end repair, adapter ligation and size selection), and single-end 36-bp reads were generated. Reads were aligned to human genome assembly hg38, and broad peaks were called with MACS [137, 138] (*q <* 0.05, broad cutoff = 0.1). Stringent per-sample quality control criteria were also applied (≥ 15 × 10^6^ unique reads, non-redundant fraction ≥0.3, fraction of reads in peaks ≥ 0.05). A union peak set of 92,401 peaks was established to leverage information across donors. Peak counts were computed for the final set of *n* = 592 donors, then normalized using a limma-voom pipeline [139, 140]. Batch effects for 62 libraries were removed using ComBat [141, 142]. Other technical factors (RNA integrity number, postmortem interval) and biological covariates (sex, age at death), along with the top 15 genotype PCs, were included as covariates the association and fine-mapping analyses. Additional hidden confounders included as covariates in these analyses were estimated from PCA applied to the normalized methylation matrix. The number of hidden confounders, 113, was determined by the Marchenko-Pastur limit based on the input methylation matrix.

SNP-peak association testing was performed using TensorQTL [112]. Significance of the SNP-peak associations was determined by computing *q-*values [143] across all tested variants, separately for each H3K9ac peak.

For fine-mapping, SuSiE-topPC and fSuSiE with the IS prior were applied to the data from each fine-mapping region (the TADs) in the same way as in the simulations, except that we set L = 20. Since fSuSiE requires a minimum of 16 peaks, we did not finemap the TADs that had fewer than than 16 peaks. We also did not consider SNPs with minor allele count (MAC) ≤ 5 (which corresponds to MAF ≤ MAC*/*2*n* = 0.4%).

#### DNA methylation data, association analysis and fine-mapping

DNA methylation was assayed using the Illumina 450K array [61, 114] and processed via the SeSAMe pipeline [144]. SeSAMe applies NOOB background correction, nonlinear dye-bias normalization, and pOOBAH-based probe filtering to generate high-quality methylation proportions (*β* values) [144]. The resulting *β* values were then transformed using a logit transformation. Missing data in the transformed matrix were imputed using flashier, a factor analysis method that estimates latent factors from the observed methylation data and predicts missing entries. As with other molecular datasets, additional hidden confounders were estimated by applying PCA to the imputed methylation matrix, with the number of hidden confounders (38) determined by the Marchenko-Pastur limit. Technical covariates and biological covariates were included as covariates in the association and fine-mapping analyses. Association analysis (TensorQTL) and fine-mapping (SuSiE-topPC, fSuSiE) were performed as described for the H3K9ac ChIP-seq data.

An additional issue specific to the Illumina 450K array is that SNPs within CpG probe sequences can create false methylation signals [145, 146]. Such polymorphisms can interfere with probe hybridization or affect signal detection, particularly when located within the CpG dinucleotide or the adjacent single base extension site, producing false intermediate methylation values often misinterpreted as epigenetic differences. To address this issue, we removed all all SuSiE-topPC and fSuSiE CSs in which one or more of the SNPs overlapped with a CpG probe in the Illumina 450K array (source: `IlluminaHumanMethylation450K.rda` from https://github.com/Yang9704/MethylCallR/ [147]). Reassuringly, most of the CpG probes overlapping the methylation CSs were also included in a list of “suggested probes for removal” from [146] (specifically, these are the CpG probes flagged as “discard” in “Additional file 2”). We also did the same for the SNP-CpG associations, although this removed only a very small number of SNP-CpG associations meeting the *q*-value threshold.

#### AD GWAS fine-mapping and colocalization

To finemap AD risk loci, we constructed an LD reference panel using genotype data from the Alzheimer’s Disease Sequencing Project (ADSP), which consisted of approximately 17,000 WGS individuals of European ancestry. QC on two recent AD GWAS meta-analysis summary statistics [83, 84] was then performed against this reference panel, including allele harmonization and LD mismatch detection using SLALOM [148], which flagged suspicious variants and then removed them. We then ran SuSiE-RSS [24, 25] using the *z-*scores from each of the AD GWAS and the LD from the LD reference panel as input data. To prioritize AD risk loci for further investigation, we ran COLOC (version 5) [85] to assess colocalization of the putative causal AD SNPs with RNA expression, protein expression, DNA methylation and H3K9ac SNPs.

To benchmark fSuSiE+COLOC against a standard per-trait pipeline, we additionally ran univariate SuSiE on each molecular location (each CpG for methylation, each peak for H3K9ac) within the regions nominated by fSuSiE, followed by COLOC v5 applied to the resulting per-location credible sets. Of 18 loci identified by fSuSiE+COLOC at 95% colocalization CSs, 5 were also recovered by SuSiE+COLOC on the same regions and 13 were specific to fSuSiE+COLOC. Of the shared set, 60% (3/5) reached AD GWAS *p <* 5 × 10^−8^ at the lead SNP; of the fSuSiE-only set, 53.8% (7/13), with the largest fSuSiE-only AD GWAS *p*-value being 7.69 × 10^−6^.

#### Excess-of-overlap enrichment analysis

We performed an excess-of-overlap (EOO) enrichment analysis of the haSNPs and mSNPs using a collection of predefined functional annotations from the Baseline-LD v2.2 model among common SNPs (MAF *>* 0.05) in the 1000 Genomes Project [80, 81]. For each molecular trait (H3K9ac, DNA methylation) and method (fSuSiE, SuSiE-topPC), we defined the “positive set” for enrichment analysis using variants within 95% credible sets (CSs) that had PIP *>* 0.5 and complete functional annotations available in Baseline-LD v2.2 common SNPs. This filtering process yielded 8,264 CS variants from a total of 74,065 variants across DNA methylation and H3K9ac credible sets for fSuSiE, and 749 out of 97,595 for SuSiE-topPC. Following [96], we tested whether selected SNPs showed greater overlap with functional annotations than expected by chance, with control set defined as follows: for each selected SNP, 5 SNPs outside the selected set were chosen to match based on LD score and minor allele frequency. For each annotation, enrichment was calculated as the ratio of the proportion of selected SNPs overlapping the annotation to the proportion of control SNPs overlapping the annotation. Statistical significance and confidence intervals were estimated using a jackknife resampling procedure, where the enrichment ratio was recalculated after iteratively removing each chromosome. Mean and standard error of enrichment estimates were computed from jackknife samples; annotations were considered significantly enriched if the 95% jackknife confidence interval for the enrichment ratio did not include 1.

## Data availability

The datasets analyzed, including the ADSP R4 WGS genotype data, are available for application and download from the NIAGADS Data Sharing Service, https://dss.niagads.org. The code implementing the data processing and analysis pipelines for is available at https://statfungen.github.io/xqtl-protocol. Data generated from numerical studies and analysis to prepare figures for the paper is available at https://github.com/stephenslab/fsusie-experiments/. A subset of fine-mapped QTL obtained from our analyses are available for peer review purposes, at https://github.com/stephenslab/fsusie-experiments/blob/main/data/README.md. The complete set of QTL data and QTL-GWAS integration models will be made publicly available at https://synapse.org prior to publication for registered Synapse users, as per the Data Management and Sharing policy of the FunGen-xQTL project. Other data sets used include: 1000 Genomes Phase 3 whole-genome sequencing data, https://ftp.1000genomes.ebi.ac.uk/vol1/ftp/release/20130502/; GENCODE Ensembl/Havana database, https://useast.ensembl.org/info/genome/genebuild/annotation_merge.html; MethylCallR R package, https://github.com/Yang9704/MethylCallR.

## Code availability

The fsusieR R package is available on GitHub at https://github.com/stephenslab/fsusieR (3-clause BSD license). The code used to perform the simulations and generate the manuscript figures is available at https://github.com/stephenslab/fsusie-experiments/. Other software and R packages used in this work include: VCPA 1.1 (http://www.niagads.org/VCPA/); BWA-MEM 0.7.15 (https://github.com/lh3/bwa/); GATK 4.1.1 (https://gatk.broadinstitute.org); FastQC 0.12.1 (http://www.bioinformatics.babraham.ac.uk/projects/fastqc/); fastp 0.23.4 (https://github.com/OpenGene/fastp/); STAR 2.7.11b (https://github.com/alexdobin/STAR/); WASP (https://github.com/bmvdgeijn/WASP/); Picard 3.1.0 (https://broadinstitute.github.io/picard/); RNA-SeQC 2.4.2 (https://github.com/getzlab/rnaseqc/); MACS2 (https://github.com/macs3-project/MACS/); ComBat (https://github.com/epigenelabs/pyComBat/); PLINK 1.9 (https://www.cog-genomics.org/plink/); TensorQTL 1.0.8 (https://github.com/broadinstitute/tensorqtl/); sim1000G 1.40 (https://github.com/adimitromanolakis/sim1000G); WGBSSuite 0.4 (https://github.com/SystemsGeneticsSG/WGBSSuite/); SeqSIMLA (https://seqsimla.sourceforge.net/); susieR 0.14.7 (https://github.com/stephenslab/susieR/); coloc 5.2.3 (https://github.com/chr1swallace/coloc/); ashr 2.2-63 (https://github.com/stephens999/ashr/); mixsqp 0.3.18 (https://github.com/stephenslab/mixsqp/); wavethresh 4.7.2 (https://cran.r-project.org/package=wavethresh); limma 3.56.2 (https://bioconductor.org/packages/limma/); qvalue 2.32.0 (https://github.com/StoreyLab/qvalue/); flashier 1.0.21 (https://github.com/willwerscheid/flashier/); R 4.3.3 (https://www.r-project.org).

## Acknowledgments

We thank Kevin Luo, Xin He, and members of the Wang and Stephens labs for discussions and support, and Yuqi Miao for initial input on simulation study design. We thank the staff at the Research Computing Center at the University of Chicago for providing the high-performance computing resources used to implement the numerical experiments. We thank Angela Helfrich and Mark Bronnimann from Amazon Web Services for providing cloud computing support. We also thank the members of the Alzheimer’s Disease Sequencing Project Functional Genomics Consortium (FunGen-AD) for providing the FunGen-xQTL resource. This work was supported in part by NIH grants R01HG002585 and R35GM153249 (to M.S.), NIH grants R01AG076901 and R01AG086467 (to G.W., H.S., A.L.), U01AF072572 (to P.L.D.) and a grant from the Urbut Family Foundation (to G.W.). This project is supported by the Eric and Wendy Schmidt AI in Science Postdoctoral Fellowship, a Schmidt Sciences, LLC program. Additional support came from the University of Chicago Data Science Institute through the 2024 AI+Science Research Initiative. This research was conducted using data from the Religious Orders Study and the Rush Memory and Aging Project (ROSMAP). We thank the participants and investigators of these studies.

## Author contributions

G.W. and M.S. jointly supervised research. W.D. developed the method and algorithm with input from G.W. and M.S. W.D., G.W. and M.S. conceived and designed the experiments. W.D. implemented the methods comparisons in simulations and performed the statistical analyses. W.D., H.S., P.C., and A.L. conducted data applications and interpreted the results. W.D. and P.C. implemented the fSuSiE software. P.L.D. and D.B. contributed and supervised molecular QTL data production from the ROSMAP cohort. The Alzheimer’s Disease Functional Genomics Consortium contributed additional data resources. W.D., P.C., H.S., G.W. and M.S. wrote and revised the manuscript. All authors critically reviewed the manuscript, suggested revisions as needed, and approved the final version.

## Supplementary text

### Background on wavelet regression

#### Wavelet transform and the concept of scale

To introduce the basic ideas, we consider the discrete wavelet transform (specifically, the unscaled Haar wavelet transform). Then we present the concept of scale (or resolution level), and define notation for the wavelet coefficients. For a more detailed introduction, see Nason[75].

Let ***f*** = (*f*_1_, …, *f*_*T*_) be a time series such that *T* = 2^*S*^. The wavelet transform of *f*_1_, …, *f*_*T*_ is a set of *T* coefficients, denoted by *d*_*s,l*_, with *s* ∈ [0, *S* −1], in which each coefficient summarizes the variation of ***f*** on an interval of [1, *T*]. The index *s* is the “scale” of a wavelet coefficient (WC); WCs at a small *s* (say, 0, 1 or 2) capture variation of ***f*** over large intervals whereas WCs at large *s* capture variation over small intervals of [1, *T*].

To give some intuition for how wavelet coefficients are generated from an observed signal, consider the unscaled Haar wavelet transform, a wavelet transform which involves relatively simple calculations. The single unscaled WC at scale 0 is a simple sum:

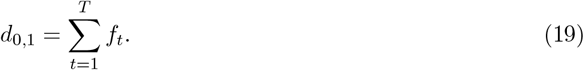

This WC is sometimes called the “height” of ***f***. The single WC at scale 1 is

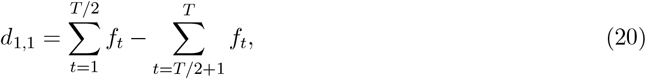

which is the difference in height between left half and right half. The two WCs at scale 2 are

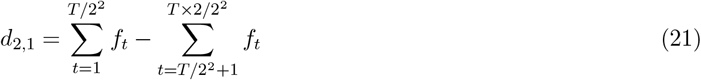

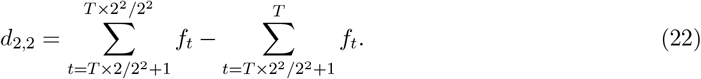

The *d*_2,1_ coefficient is the difference in the height of *g* between the first and second quarters, and *d*_2,2_ is the contrast in the height of *g* between the third and fourth quarters.

The WCs at scales 3 and larger follow a similar logic. By repeating this process at scales *s* = 3 up to *s* = *S* −1, we obtain a total of 2^*S*^ WCs. This process just described can also be equivalently formulated as matrix multiplication with orthogonal projection matrix **W**. This ability to express the wavelet decomposition as a matrix multiplication is a property of all wavelet bases, not just the Haar basis [101].

#### The undecimated wavelet transform

In the presented work, we assume that the data are evenly spaced and have *T* = 2^*S*^ measurements. However, such an assumption is not realistic in practice. The problem of applying wavelets to unevenly-spaced data of length *n* ≠ 2^*S*^ has been addressed by several authors [100] [149] [150] and is referred to as an undecimated/non-decimated wavelet transform or lifting scheme. In our implementation, we used the approach of Kovac and Silverman [150]. The approach of Kovac and Silverman [150] for non-decimated (size ≠ 2^*S*^) and unevenly-spaced data consist on interpolating the observed data on a evenly spaced grid of length 2^*S*^. The interpolated data is then used as our response variable and we apply fSuSiE directly. We provide an additional description of the procedure of Kovac and Silverman [150] is described below.

Suppose, we observe **Y** at *T* ^′^ unevenly spaced time points *τ*_1_, …, *τ*_*T*_ *′* and we assume that *T* ^′^ is not a power of 2. For simplicity, we consider these time points to be between [0, 1]. Kovac and Silverman suggested considering a new set of evenly space-time point, *t*_0_, …, *t*_*T*_ where *T* = 2^*S*^, *S* ∈ ℕ, in which 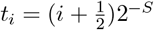 and *S* = *min*{*s* ∈ ℕ, 2^*s*^, ≤ *n*}.

We generate a new matrix of observation **Y**_*inter*_ by interpolating each individual curve on this grid as follows. For a given individual *k* we define **Y**_*k,inter*_(*t*) (the interpolated value of curve of individual k at time point t) as:

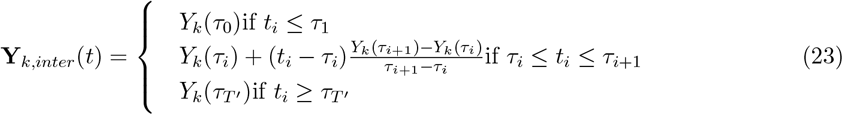

The expression can then be written in terms of the matrix as follows:

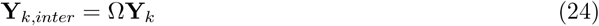

Where Ω is the matrix corresponding to the interpolation procedure described in equation (23). The resulting vector **Y**_*k,inter*_ is of length 2^*S*^ and corresponds to the linear interpolation of **Y**_*k*_ on the evenly spaced time points *t*_0_, …, *t*_*T*_.

### Elementary posterior expressions used to derive fSuSiE computations

To derive our VA approximation of the posterior of model (12), we start by computing some key quantities of some simpler models. These quantities will serve as building blocks for our estimation procedure. Here, we consider that the matrices **Y, D** (response data) and **X** (covariates) are centered in a column-wise fashion. Similarly the corresponding vectors **Y, d** (response data) and ***x*** (covariates) are also assumed to be centered.

#### Simple Bayesian functional regression

The most basic building block in our derivations is a wavelet regression model in which the effects have independent normal priors. We call this the *simple Bayesian functional regression model:*

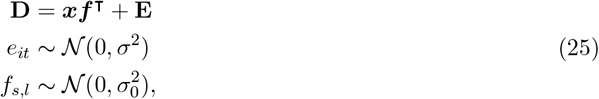

where **D** is a matrix of dimension *N* × *T*, ***x*** is a vector of length *N*, ***f*** is a vector of length *T*, and **E** is a matrix of dimension *N* ×*T* in which the individual entries are denoted by *e*_*it*_. Although not strictly needed to describe this simple functional regression model, here we introduce the indices *s* and *l* for wavelet scale and wavelet location; in this notational convention, each pair (*s, l*) corresponds to an integer from 1 to *T* so that *f*_*s,l*_ is an element of the vector ***f***. This notation will become moree useful below for thee more complex functional regression models.

The maximum-likelihood estimate of ***f*** and its corresponding variance are given by

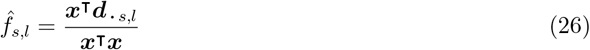

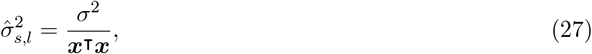

in which ***d***_***·*** *s,l*_ denotes a column of **D**.

The posterior distribution of ***f*** is given by

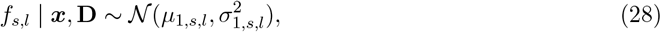

where

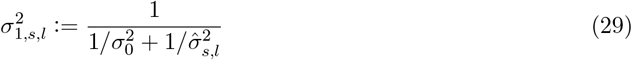

The Bayes factor comparing with the null hypothesis (***f*** = **0**) against the alternative (***f*** ≠ **0**) is

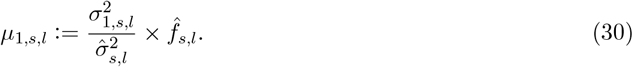

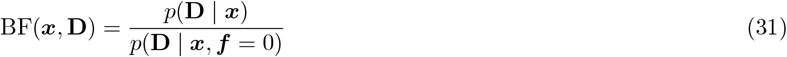

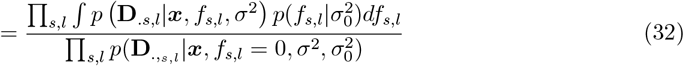

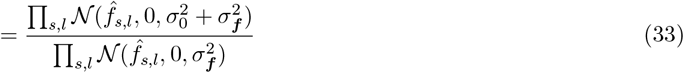

in which 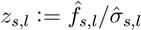.

#### Bayesian simple functional regression with mixture prior

Given the previous derivations, we can now compute the full posterior under the SFR model using a scale-dependent mixture prior. As previously, we start by establishing the basic result for the simple Bayesian functional regression and then extend it to the SFR model. Let’s consider the following simple function regression model with a mixture prior:

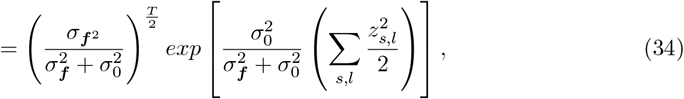

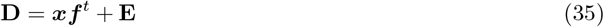

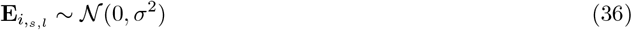

As mentioned in the main text, to lighten the notation we assume that the grid of 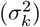 is the same for every scale; in practice, we use a different grid of sigma for each scale. We start by introducing the following latent variables:

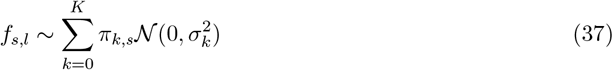

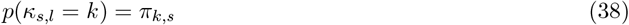

Using results from equation (29) and (30), the posterior distribution of *f*_*s,l*_ given that *κ*_*s,l*_ = *k* is

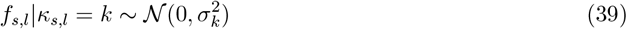

where

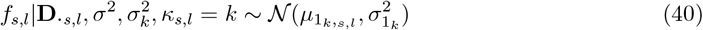

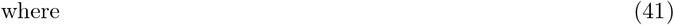

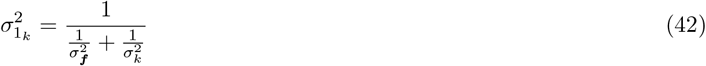

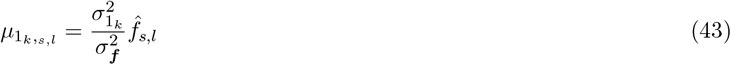

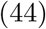

Thus the posterior first and second moment of ***f*** is given by

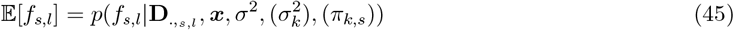

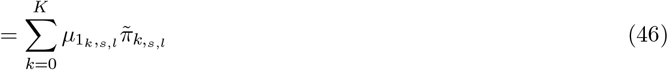

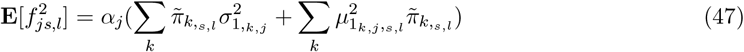

Where 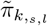 the posterior probability of *κ*_*s,l*_

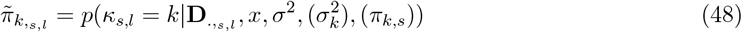

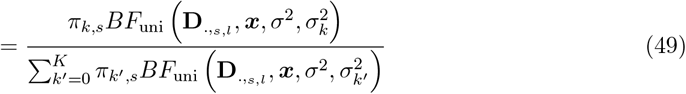

Where *BF*_uni_ is defined as

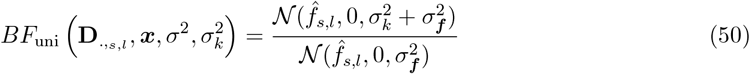

Finally, the Bayes factor for the simple Bayesian functional regression using a mixture prior can be computed as follows:

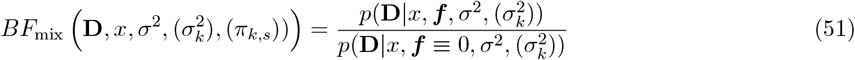

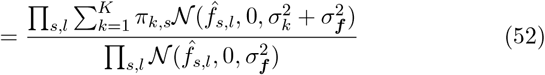

#### Bayesian simple single functional regression

Let’s extend the SFR model with a simple normal prior to a multivariate setting. Here we suppose that only one of the covariate in **X** is affecting the outcome. Our goal is to corner this single covariate by modeling the effect of **X** as follow,

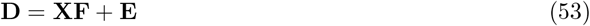

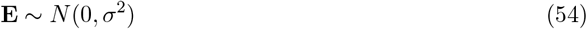

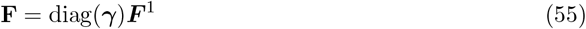

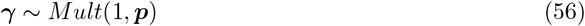

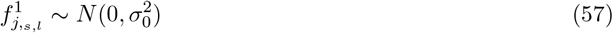

Where **X** is a matrix of dimension *N* ×*J*, **F** is a matrix of length *J* ×*T*, where exactly only one row is different from 0_*T*_, **E** is a matrix of dimension *N* ×*T*, ***f*** is a vector of length 2^*S*^ = *T*, ***p*** is a vector length *J*, with components between 0 and 1 and the sum of its components sums to 1. This model can be fitting using the quantities defined in the previous section as well as the following posterior distribution for ***γ*** and *F*. Following the seminal work of Wang and colleagues [24]:

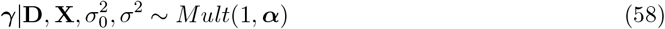

Where the component of the posterior probability of inclusion ***α*** can be computed as following using Bayes rules

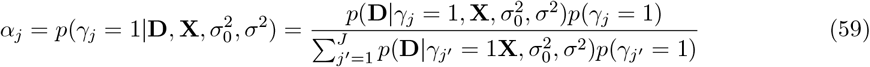

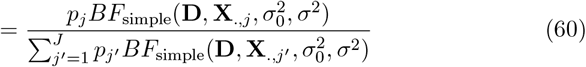

Therefore, the posterior first and second moments of the column of the entries of **F** in model (53) can be computed as follow:

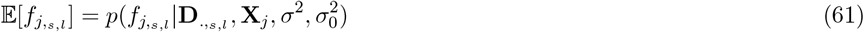

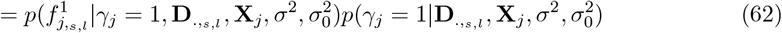

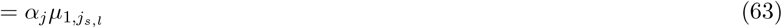

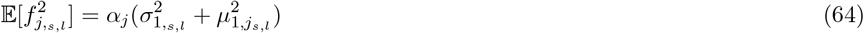

Where 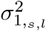 is defined as in (29) and 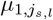 is defined as in (30).

#### Bayesian single functional regression with mixture prior

Finally let’s consider model (12) with mixture prior (written below for convenience):

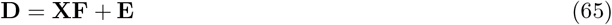

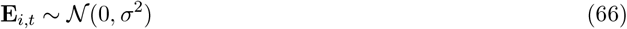

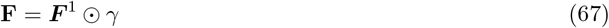

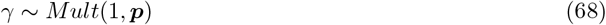

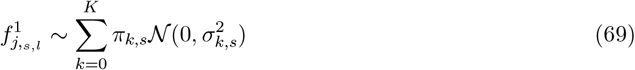

Fitting (12) requires computing the following quantities:

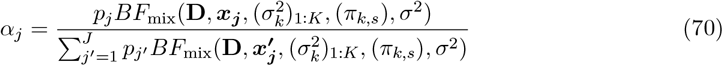

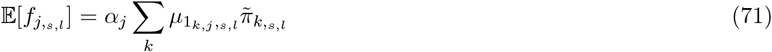

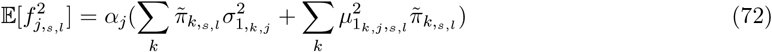

Which is straightforward to compute using results from previous section, see Bayesian simple functional regression with mixture prior.

#### *Estimating the hyperparameters* (*π*_*k,s*_)

So far for any given SFR we assume that the hyperparameters (*σ*^2^, 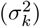, (*π*_*k,s*_)) are knwon. We estimate *σ*^2^ using the residuals (see section 13) and the grid 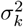is selected in an automatic fashion (see [104]) prior to the estimation procedure. We estimate the remaining *K* ×*S* hyperparameters (*π*_*k,s*_) in a an empirical Bayes way, using an EM algorithm that we detail below. First, notice that the log-likelihood of an SFR can be written as

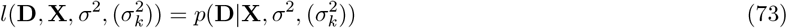

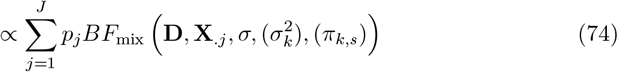

To lighten the notation let’s denote 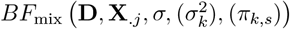 as ***BF***_*j*_ ((*π*_*k,s*_)). Let 𝒳 be a random variable generated from the following mixture:

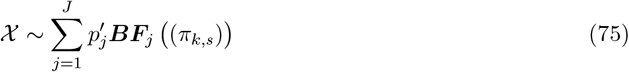

Where 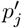 is the proportion of the component *j* in the mixture. Let’s now introduce the latent variable *Z* with value in ⟬1 : *J* ⟭, such that *X*| *Z* = *j* ~ ***BF*** ((*π*_*k,s*_)) and 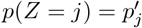

Then the complete log-likelihood can be written.

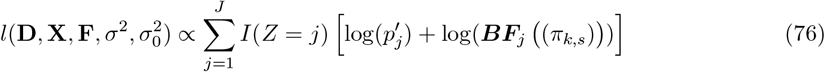

Knowing *Z*|*X*, we can maximize the expected log-likelihood

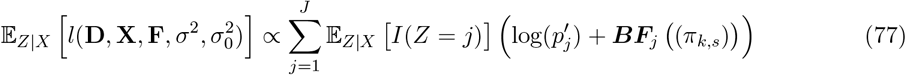

#### E step

Thus E step can be computed as

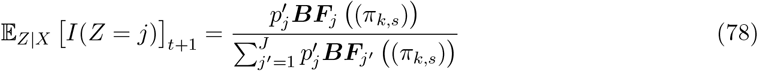

We note *ζ*_*j,t*+1_ = **E**_*Z*|*X*_ [*I*(*Z* = *j*)] *t*+1

#### M step

Thus the complete log-likelihood can be written as

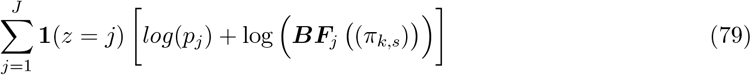

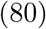

Thus the expected value of the complete log-likelihood is proportional (with respect to (*π*_*k,s*_)) to

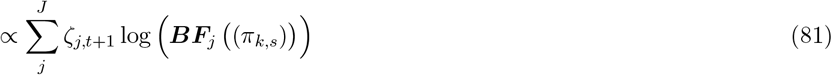

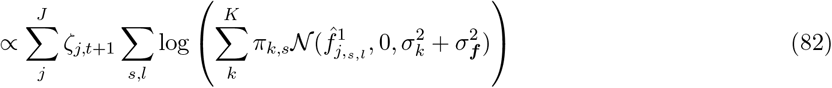

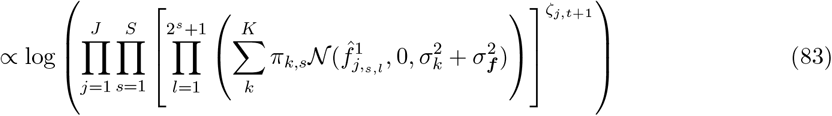

Equation (83) can be maximised using a weighted EM algorithm to optimise over on set of (*π*_*k,s*_)_1:*K*_ (here s is fixed) so corresponding S independent weighted maximization problems

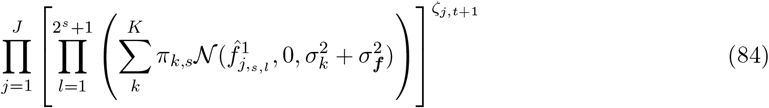

Given that the only unknown quantities are the mixture proportions, these problems are convex and can be solved efficiently; we use the *mixsqp R* package to solve these problems [151].

Note that, in practice when the number of SNPs and/or number of wavelet coefficient is large (*J >* 10, 000, *T >* 1000), optimizing (84) may require to allocate a *J*× (*T* ×*K*) matrix which can lead to requesting very large amount of memory. In practice, we alleviate this problem by optimizing (84) only using SNPs with a large value *ζ*_*j,t*+1_ (e.g. using the 5000 SNPs which have the *ζ*_*j,t*+1_ largest values) and limit the number of wavelet coefficient to a maximum of 1024. These parameters can be changed by the user, but may require a large amount of memory available.

### Algorithms for fitting the fSuSiE model

In the line of the work of [24], we show that given **F**^(1)^, …, **F**^(*R*−1)^ are known, estimating effect *R*, (i.e., **F**^(*R*)^) from model (13) corresponds to fitting an SFR model using as response **R** = **D** − **XF**_*R*−1_ where 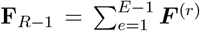 (see Appendix section 13). This suggests that model (13) can be fitted using an iterative algorithm that estimates each element of 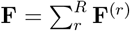 one at a time using an SFR, while holding the other constant. In the vein of [24], we use an Iterative Bayesian stepwise selection (IBSS) to fit the *fSuSiE* model. We detail our IBSS method in Algorithm 1. Similar to the IBSS in the univariate case, our algorithm is simple and has a complexity *O*(*NJTL*) per outer-loop iteration. In particular, the IBSS described in Algorithm 1 is a coordinate ascent algorithm that optimizes a VA of the posterior distribution of **F**^(1)^, …, **F**^(*R*)^ under the *fSuSiE* model (13). The key idea of VA is to find an approximation of the posterior distribution 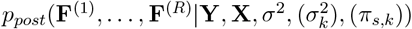 that is simpler to compute than the actual posterior. This approximation is generally obtained by imposing a certain form of the approximation of the posterior; noted:

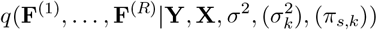

that makes the problem more tractable. As in most VA [152], we fit the fSuSiE by minimizing the Kullback-Leibler (*D*_*KL*_) divergence between *q* and *p*_*post*_.

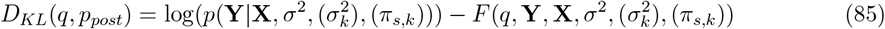

Albeit the *D*_*KL*_(*q, p*_*post*_) is in general intractable, it can be written as a sum of two terms in which only one depends on *q* (see equation (85)). Thus minimizing the Kullback-Leibler divergence in terms of *q* is equivalent to minimizing the second term of (85), which is easier to compute. The function *F* is referred to as the “evidence lower bound” (ELBO). Thus minimizing (85) corresponds to the following maximization problem.

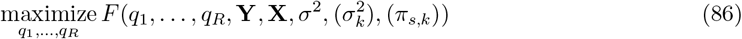

To make (86) a more tractable problem, we seek to minimize the ELBO given that *q* can be factorized as follow:

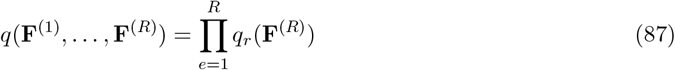

Equation (87) imposes independence between the posterior of distribution of **F**^(1)^, …, **F**^(*R*)^. As noted by Wang and colleagues [24] this factorization allows each posterior distribution *q*_*r*_ to select which are the likely covariates that contain effect *e*. While the maximization problem (86) can be challenging, optimizing (86) for one *q*_*r*_ at a time is straightforward. In particular, optimizing (86) for one *q*_*r*_ corresponds to fitting a SFR model (see the proposition below).

#### Proposition 1.

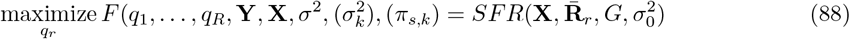

*Where* 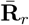 *is the expected value of the residual when removing all the effects except effect r. More formally*

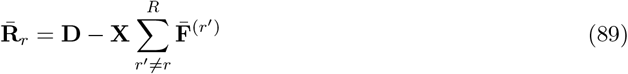

*Where* 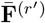 *are the expected value of* **F**^(*r′*)^*with respect to q*_*r*_*′*

Formal arguments to prove Proposition 1 are provided in Appendix section 13. Informally Proposition 1 states that the approximate posterior distribution of a given effect can be obtained by subtracting all the fitted effects to the response apart from the one we are trying to estimate. It follows the IBSS described in Algorithm 1 is a coordinate ascend algorithm that maximizes the ELBO for *q* belonging to the class of distribution defined in (87). Thus providing a simple iterative algorithm to solve (86).

#### Algorithm 1

Basic Iterative Bayesian stepwise selection for functional data

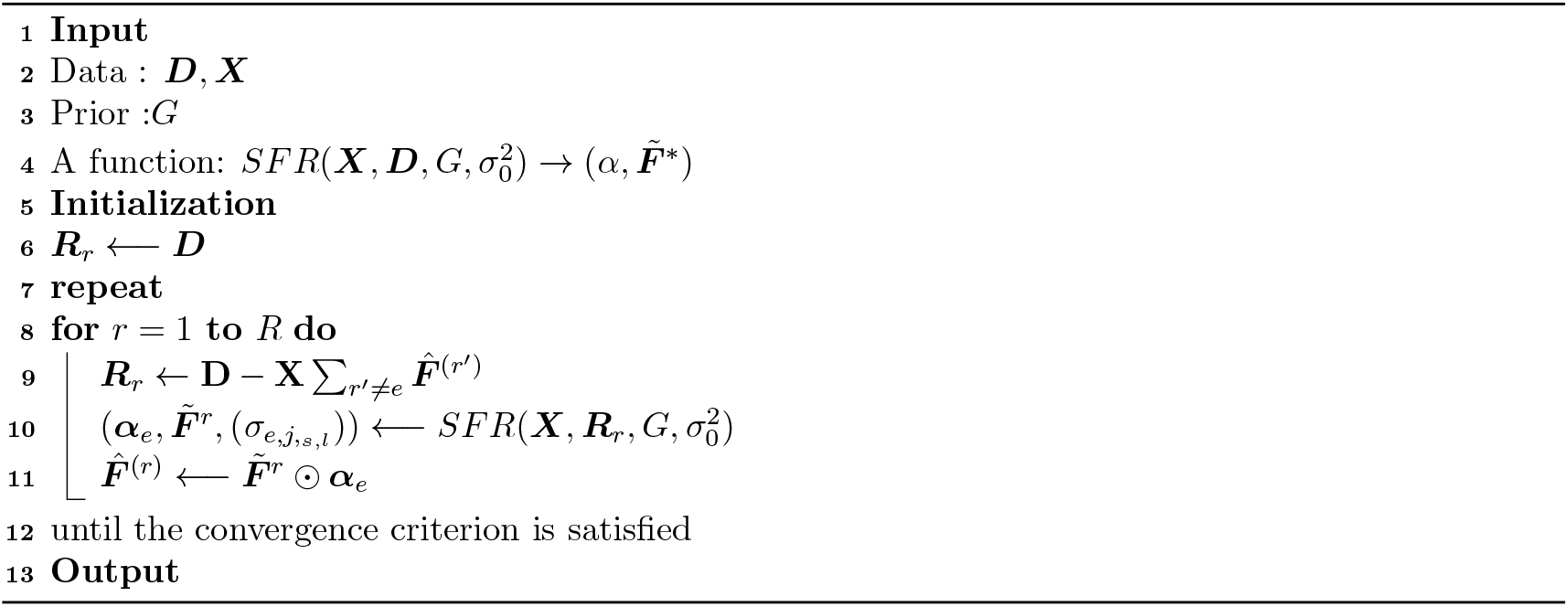

#### Hyperparemeter estimation

Until now 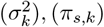, *σ*^2^ were assumed to be known. In practice, 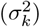 is a thin grid of values that are selected in a data-adaptive way. Briefly put, this grid contains values that range from 0 to ≈ 10 ∗ *M*, where *M* is the largest entry (in absolute value) of 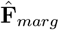, and where 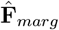 is the matrix corresponding to the marginal association between each covariate and **D** (see [104], and [153] section 1.3.2 for more details). Thus these quantities are not random and are not updated after being selected. On the other hand, (*π*_*s,k*_), *σ*^2^ are considered as random quantities and are estimated by maximizing the ELBO over (*π*_*s,k*_) and *σ*^2^.

Optimizing over *σ*^2^ is straightforward as it simply requires computing the residual sum of squares under variational approximation to then estimate *σ*^2^. Optimizing over (*π*_*s,k*_) is done by maximizing the marginal likelihood of the SFR model over (*π*_*s,k*_). This step is done using a weighted EM algorithm [154]. We provide a detailed description of this step in the Appendix section “Bayesian single functional regression with mixture prior”.

### Derivation of the fSuSiE model fitting algorithms

In this section, we describe our variational approach to fit the sum of single functions model (see equation (13)). Let’s rewrite (13) in the following form form:

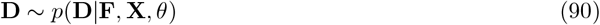

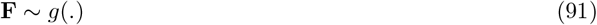

Where **D** is the matrix of the observed data, **X** is the matrix of the observed covariates, **F** is the latent/unobserved variable of interest, *g*∈ 𝒢 is a prior distribution for **F** and *θ* ∈ Θ is the set of the additional/nuisance parameters to be estimated. We adopt an Empirical Bayes (EB) approach, by first estimating our prior *g* and the noise parameters *θ* by maximizing the marginal log-likelihood:

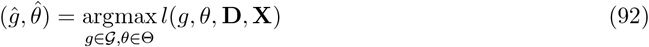

where

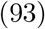

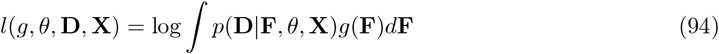

The we compute the posterior distribution for **F** using the estimated prior and the nuisance parameters:

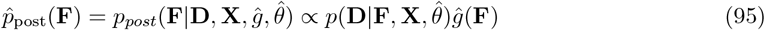

#### Posterior under the SFR model

Assuming independence between wavelet coefficients (standard assumption used in several works [50] [64] [155]), the likelihood of model (12) can be written as

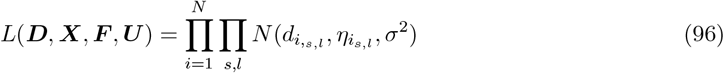

Where *N* (., *η, σ*^2^) corresponds to the density of a Normal distribution with mean *η* and variance *σ*^2^. Here 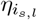 is the linear predictor of 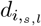 and is equal to 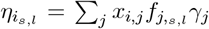. Given that (*π*_*k,s*_) and *σ* are known, the posterior distribution of **F**^(1)^ = **F**^1^ ⊙ ***γ*** under the SFR model is completely determined by the following quantities.

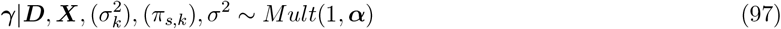

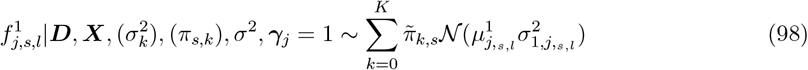

Where ***α*** = (*α*_1_, …, *α*_*J*_) corresponds to the vector of posterior inclusion probability (PIP); 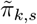 is the posterior mixture weight; 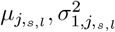 are the posterior mean and posterior variance of 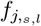 under the normal prior 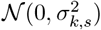. All these quantities can be computed using summary statistics from the regression of each column of **D** on each of the columns **X** (*J* ×*T* regressions). The derivations to obtain these quantities are provided in equation (61). For ease of presentation, we now refer to SFR as a function that takes as input **X, Y**, *σ*, (*σ*_*s,l*_), (*π*_*s,l*_) and returns ***α*, F**^(1)^, (*σ*_1,*s,l*_), formally written as :

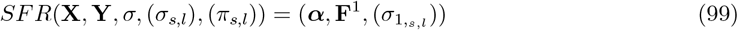

#### K-L divergence and the ELBO

As computing the full posterior of model (13) in an intractable problem, we use a VA. More precisely, we seek at finding a distribution *q* that belongs to a certain class of function 𝒬 (detailed in equation (104)) that minimizes the KL divergence with the true posterior of model (13). For the sake of pedagogy, let’s rewrite the KL divergence in terms of evidence and evidence lower bound

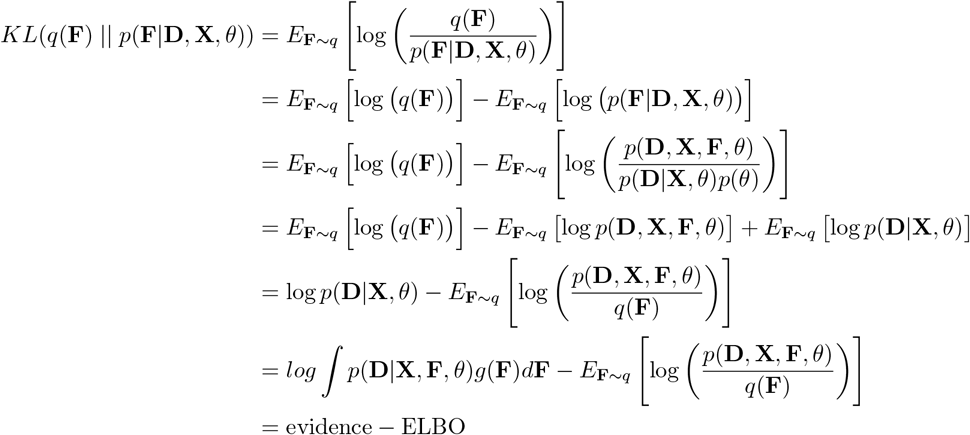

In particular, the ELBO can be written in two different way:

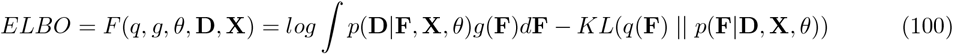

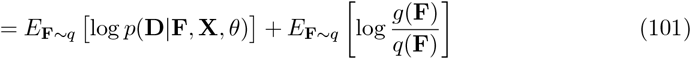

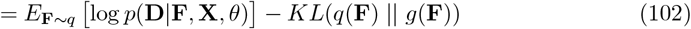

So the ELBO depends on three “parameters” *g*(.), *q*(.), *θ*. Our general optimization problem is the following:

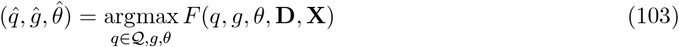

Maximizing the ELBO with respect to (*g*(.), *θ*) corresponds to our EB procedure (as the first terms of eq (100) corresponds to the marginal likelihood); maximizing the ELBO with respect to *q*∈𝒬 is the standard variational Bayes (VB) approach. We optimize (88) in a iterative fashion, by keeping *q*(.) constant and maximize over (*g*(.), *θ*). Then by keeping (*g*(.), *θ*) constant and maximize over *q*(.) (using equation (101)). The EB procedure to maximize *g* is described in Appendix section “Bayesian single functional regression with mixture prior”. Here we solely focus on maximizing the ELBO while keeping *g* and *θ* constant.

#### Form of the variational approximation

As previously mentioned, computing the full posterior of (88) is a (likely) intractable problem. Thus, we restrict the form of the “candidatr” posterior (*q*) to facilitate the optimization. More precisely, we choose a classical mean-field approximation, so 𝒬 is defined as

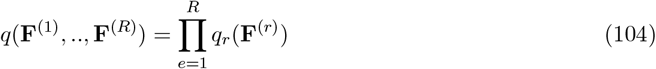

#### Analytical expression for the ELBO

For *q* ∈ 𝒬 the ELBO can be written as

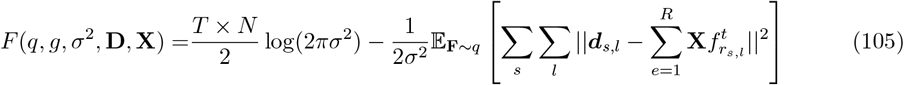

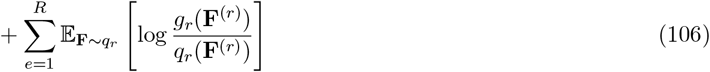

Where ||. || is the euclidean norm and ***g*** = (*g*_1_, …, *g*_*E*_) is a collection of mixture prior. In the line of Wang and colleagues[24] (proposition A.1) given that we restrict *q* to (104) the ELBO becomes

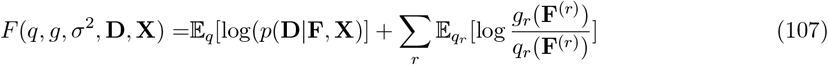

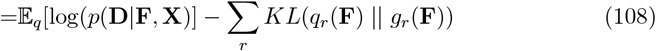

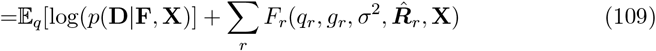

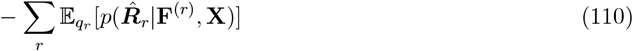

Where 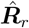corresponds to the expected residual without effect *e*, i.e.:

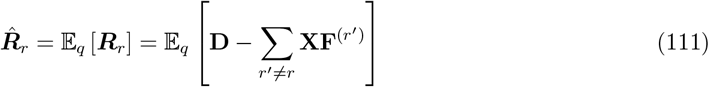

Proof for this Proposition in the univariate case is given in [24].

#### Expected residual sum of squares

Here we focus on the second term of equation (105), which we refer to as the “expected residual sum of squares” (ERSS). To compute this term, we need to compute 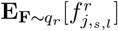 and 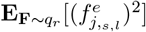, given that *g*_*r*_ is a scale mixture prior, it is straightforward given to compute these quantities using the analytical formula provided in (71) (72). Thus the “expected residual sum of squares” (ERSS) can be written as:

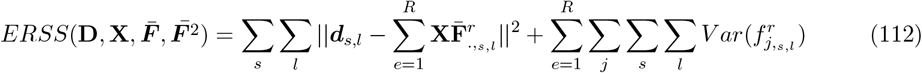

#### ELBO for the e-th single effect

Similar to the SuSiE approach 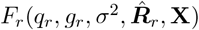 is equal to log *L*(***R***_*r*_, *g*_*r*_, *X, σ*^2^) at the maximum 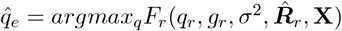, which can be computed as

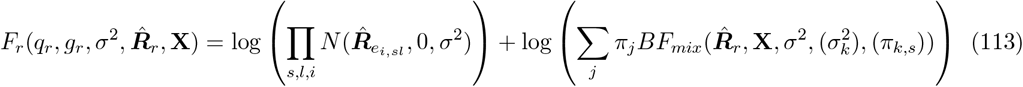

#### K-L divergence for the e-th effect

Following previous derivations, we have that

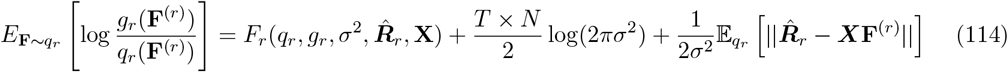

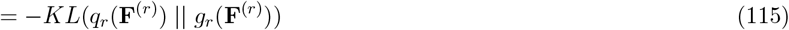

### Summary-statistics formulation of fSuSiE

An fSuSiE analysis of individual-level data (**X, Y**) can be re-expressed as a summary-statistics analysis, analogous to SuSiE–RSS [25] and mvSuSiE–RSS [33]. The required inputs are the per-SNP marginal OLS estimates 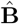 and standard errors 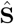 from the individual-level data (both *J* × *T*), the LD matrix **R** = (*n* −1)^−1^**X**^⊤^**X**, and the sample size *n*. We assume the columns of **X** and **Y** are centered and scaled to unit sample variance.

Because the DWT acts linearly on the position axis, the wavelet-coefficient marginal OLS estimates follow from the individual-level ones by

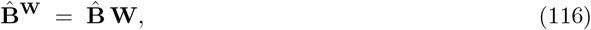

obtained by post-multiplying **X**^⊤^**Y** by **W**. Positionwise standard errors propagate as 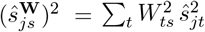; under homoscedasticity across positions this reduces to 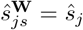 by orthogonality of **W**.

Within the IBSS loop, the partial-residual update for component *r* at wavelet coefficient *s* can be written as

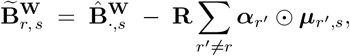

and the ELBO-based residual-variance update reduces to a function of 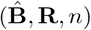 and the internal posterior state. The adaptive-shrinkage mixture prior on wavelet coefficients used in fSuSiE is itself a summary-statistics method in the style of ashr [104], whose posterior updates require only the wavelet-space summaries 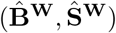 and the current mixture weights, so the fSuSiE inner loop is directly compatible with summary-statistics inputs and the prior requires no modification.

The caveats that apply to SuSiE–RSS apply here, in particular sensitivity to mismatch between in-sample and reference-panel **R**.

## Supplementary figures

**Supplementary Figure 1.**
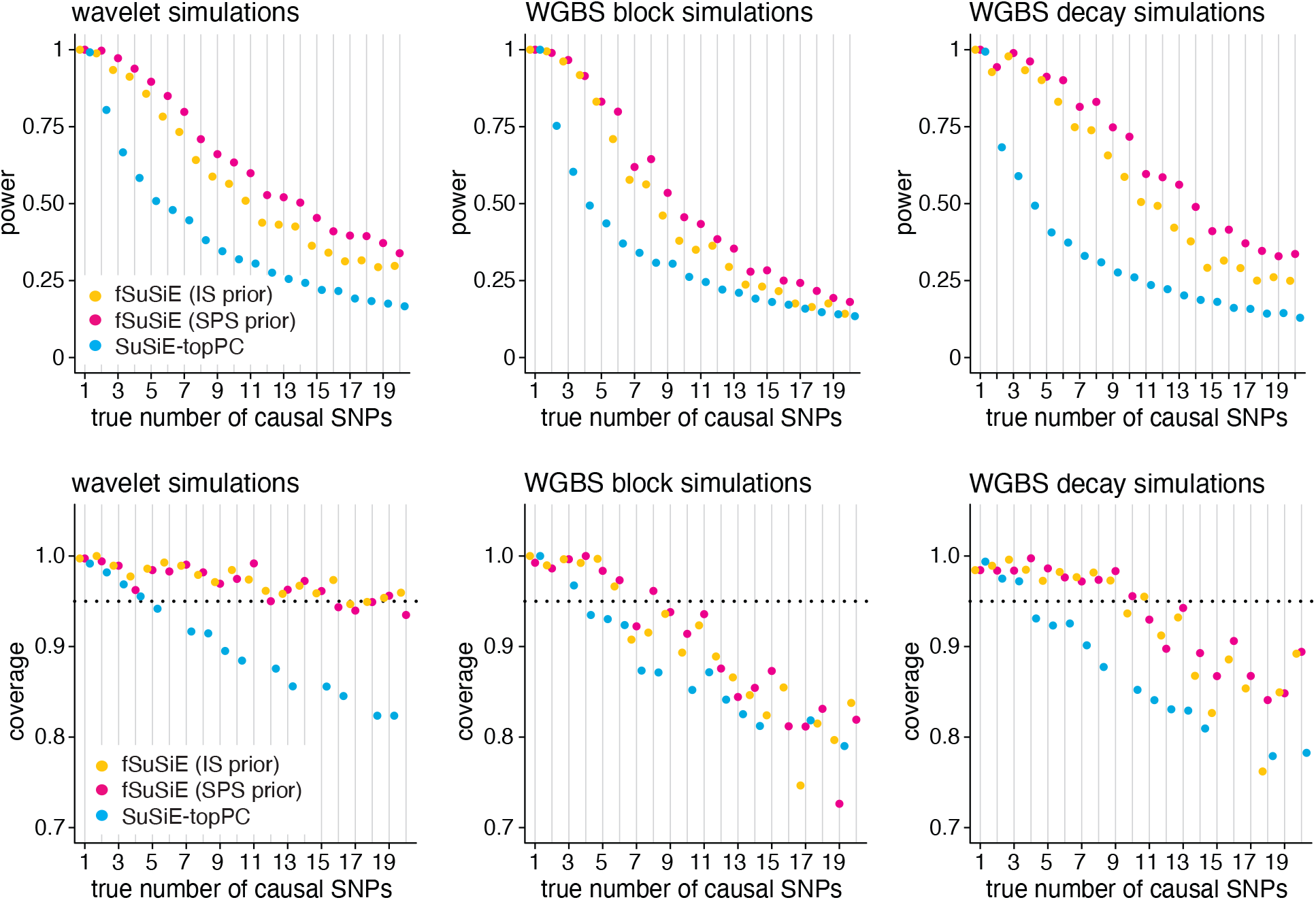
Additional assessment of SuSiE and fSuSiE CSs in simulated data sets, with performance broken down by the true number of causal SNPs. *Power* is the proportion of the true causal SNPs included in at least one CS; *coverage* is the proportion of CSs containing a true causal SNP. The dotted horizontal line shows the target coverage (95%). In all simulation settings, the data sets were simulated with *n* = 100 samples, *T* = 128 CpGs, *J* = 1,500–4,000 SNPs, and 10% total variance explained. Each plot summarizes the results from *n* = 300 simulations.

**Supplementary Figure 2.**
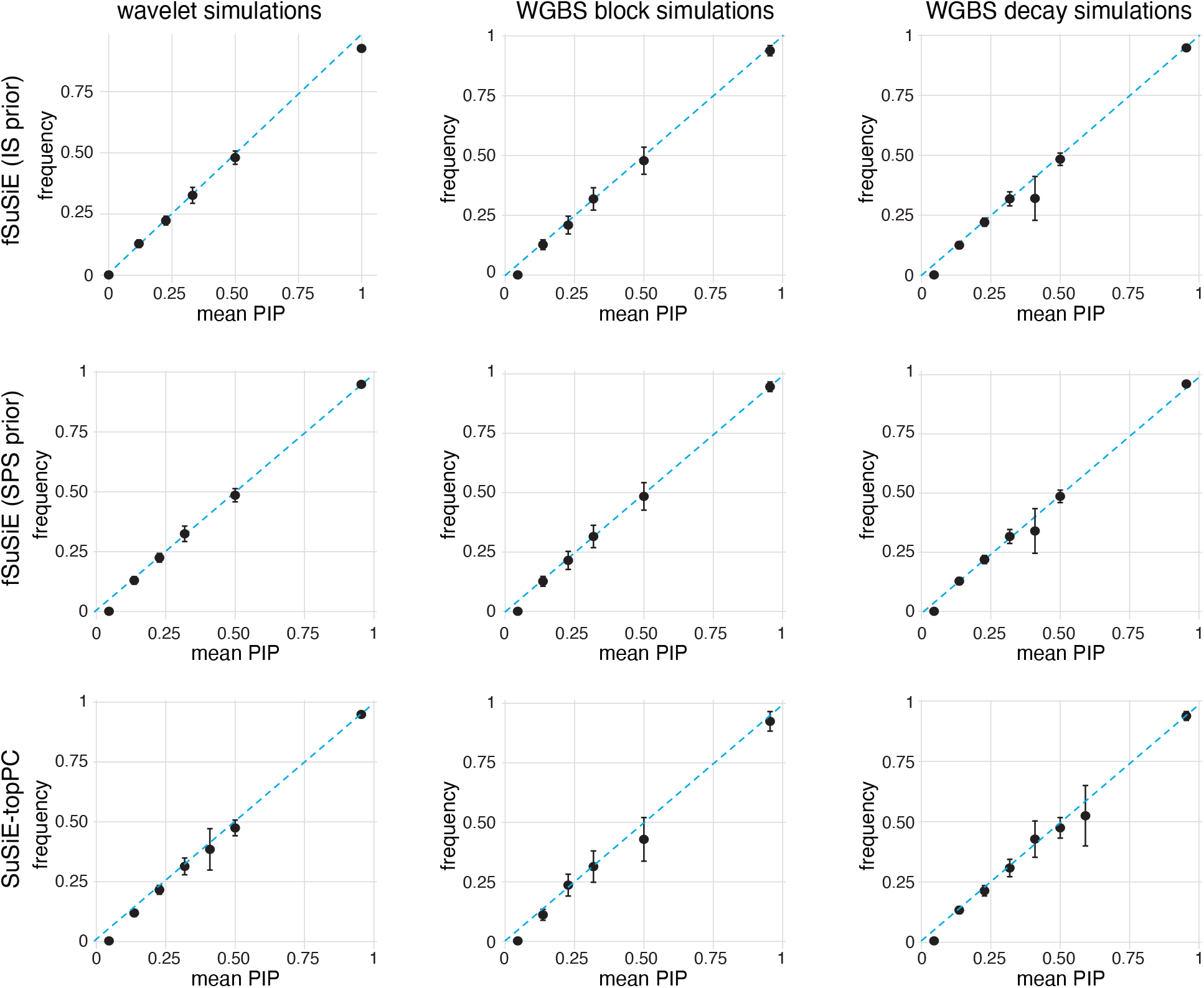
Assessment of PIP calibration in simulations. In each simulation setting, the SNPs from all simulations in that setting (*n* = 100 simulations) were grouped into bins according to their PIP (10 equally spaced bins from 0 to 1). The plots show the average PIP from each bin (x-axis) against the proportion of SNPs in that bin that are causal (y-axis). Bins with fewer than 30 SNPs are not shown in the plots. Error bars depict 2 times the empirical standard error. Note that a well-calibrated method should produce points near the diagonal. In each of the three simulation scenarios, data sets were simulated with *N* = 100 samples, *T* = 128 CpGs, *J* = 1,500–4,000 SNPs, among which 1–20 were causal, and 10% total variance explained.

**Supplementary Figure 3.**
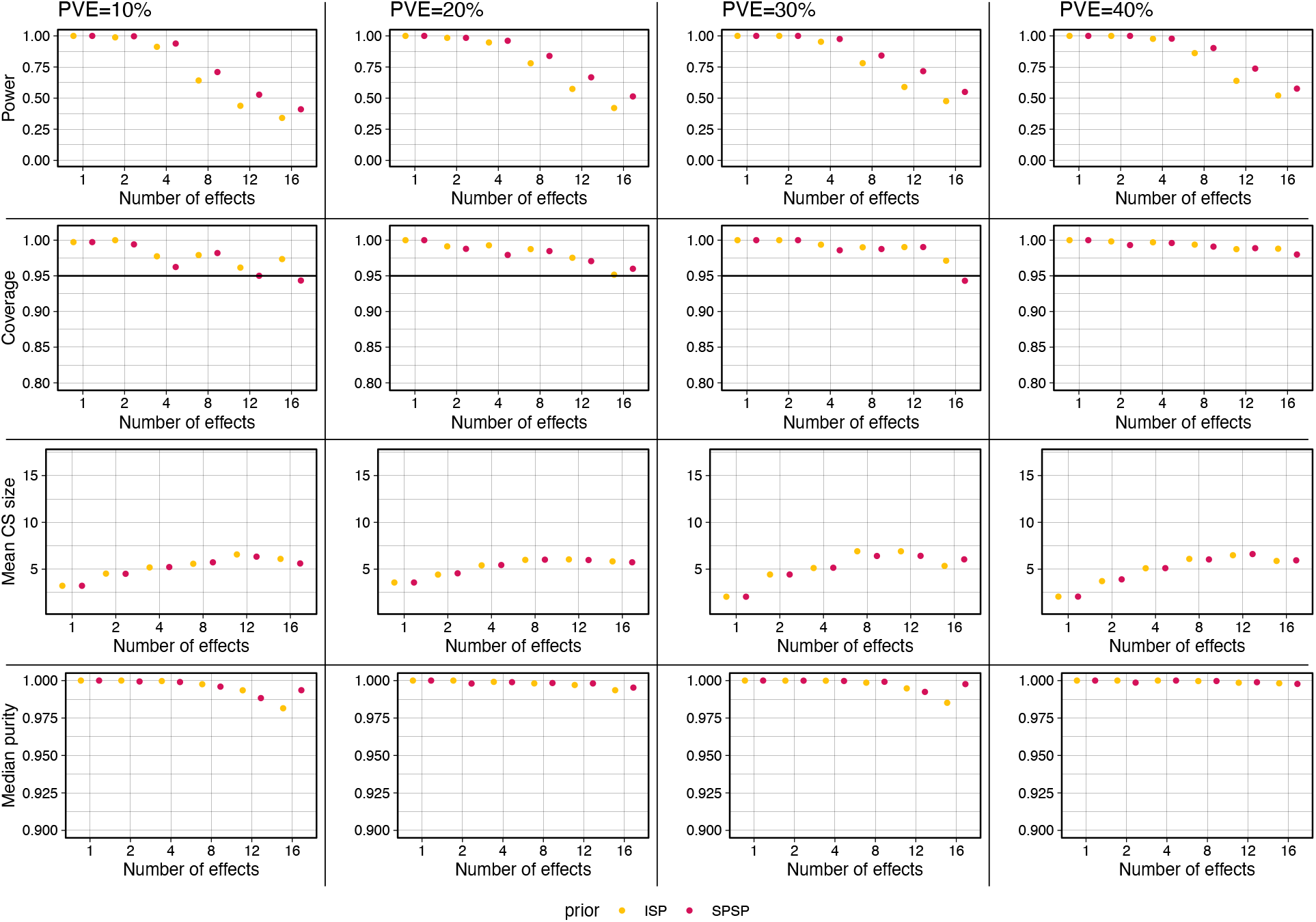
Additional assessment of fSuSiE CSs in wavelet simulations with 128 CpGs. “Purity” is the smallest absolute correlation (Pearson’s *r*) among all pairs of SNPs in a CS [24]. See the Fig. 3 caption for definitions of power, coverage and CS size. ISP = fSuSiE with IS prior; SPSP = fSuSiE with SPS prior. Each plot summarizes the results from *n* = 300 simulations. The data sets were simulated with *N* = 100 samples, and *J* = 1,500–4,000 SNPs.

**Supplementary Figure 4.**
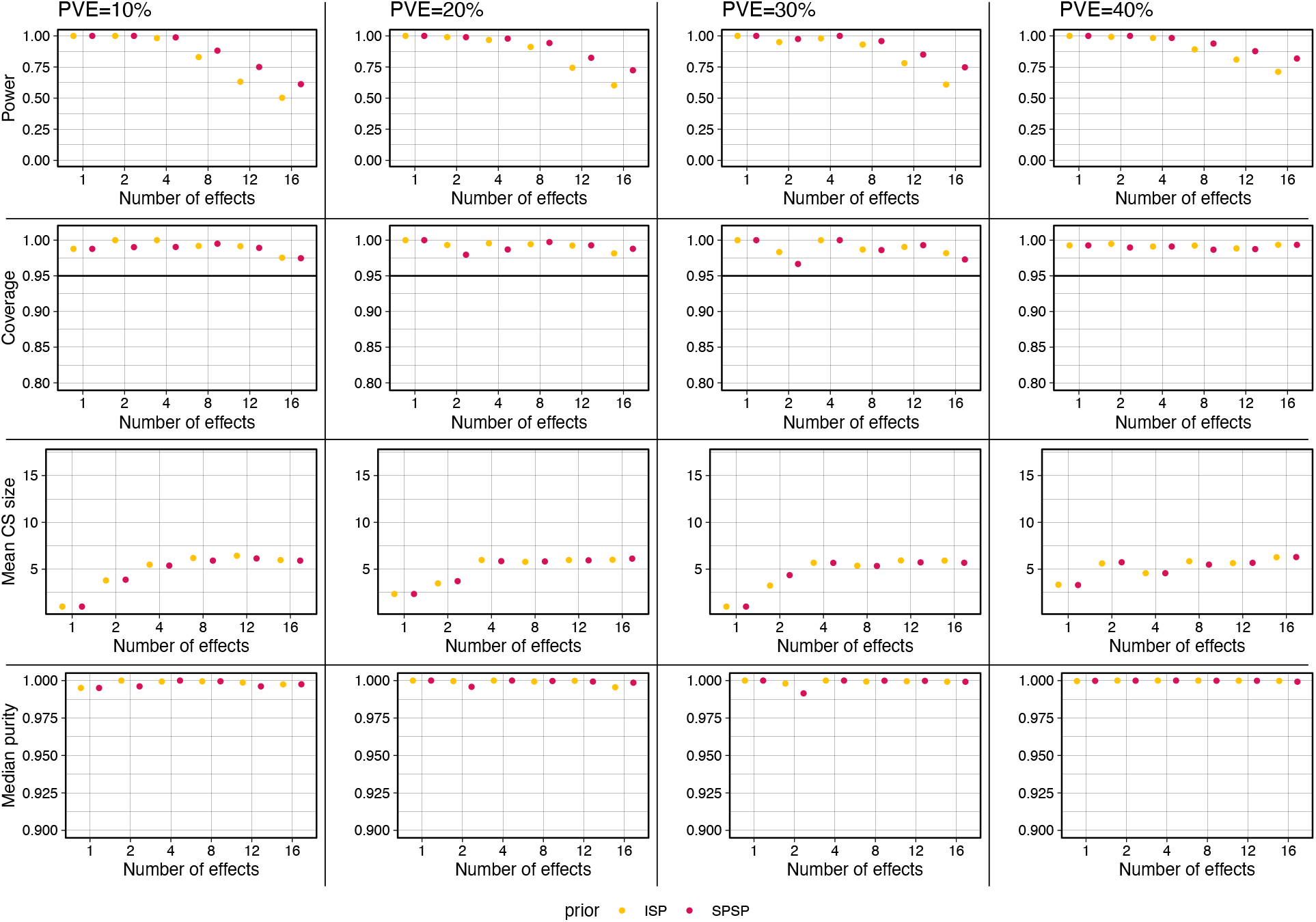
Additional assessment of fSuSiE CSs in wavelet simulations with 512 CpGs. “Purity” is the smallest absolute correlation (Pearson’s *r*) among all pairs of SNPs in a CS [24]. See the Fig. 3 caption for definitions of power, coverage and CS size. ISP = fSuSiE with IS prior; SPSP = fSuSiE with SPS prior. Each plot summarizes the results from *n* = 300 simulations. The data sets were simulated with *N* = 100 samples, and *J* = 1,500–4,000 SNPs.

**Supplementary Figure 5.**
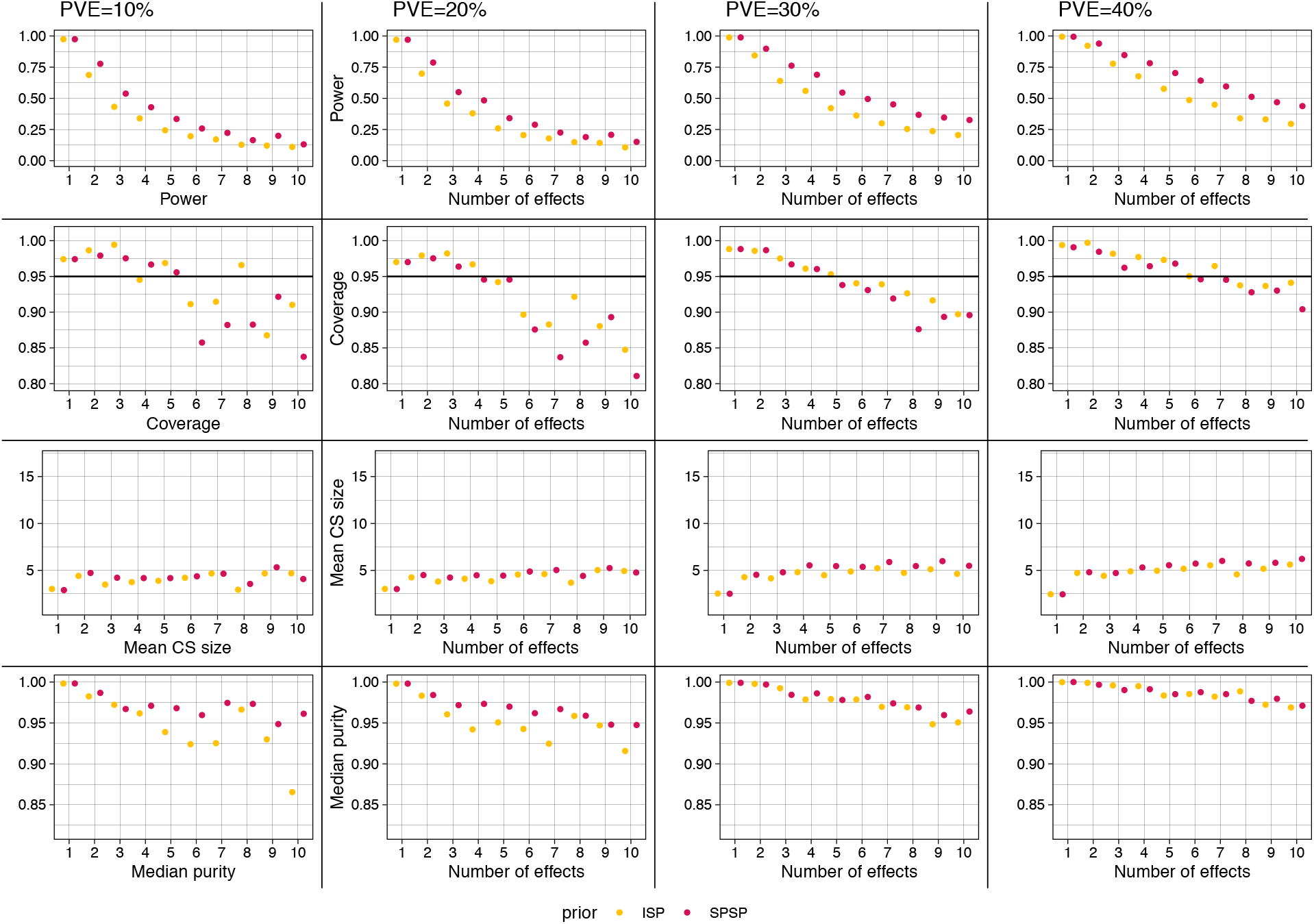
Additional assessment of fSuSiE CSs in wavelet simulations with fractional Gaussian noise, 128 CpGs. “Purity” is the smallest absolute correlation (Pearson’s *r*) among all pairs of SNPs in a CS [24]. See the Fig. 3 caption for definitions of power, coverage and CS size. ISP = fSuSiE with IS prior; SPSP = fSuSiE with SPS prior. Each plot summarizes the results from *n* = 300 simulations. The data sets were simulated with *N* = 100 samples, and *J* = 1,500–4,000 SNPs.

**Supplementary Figure 6.**
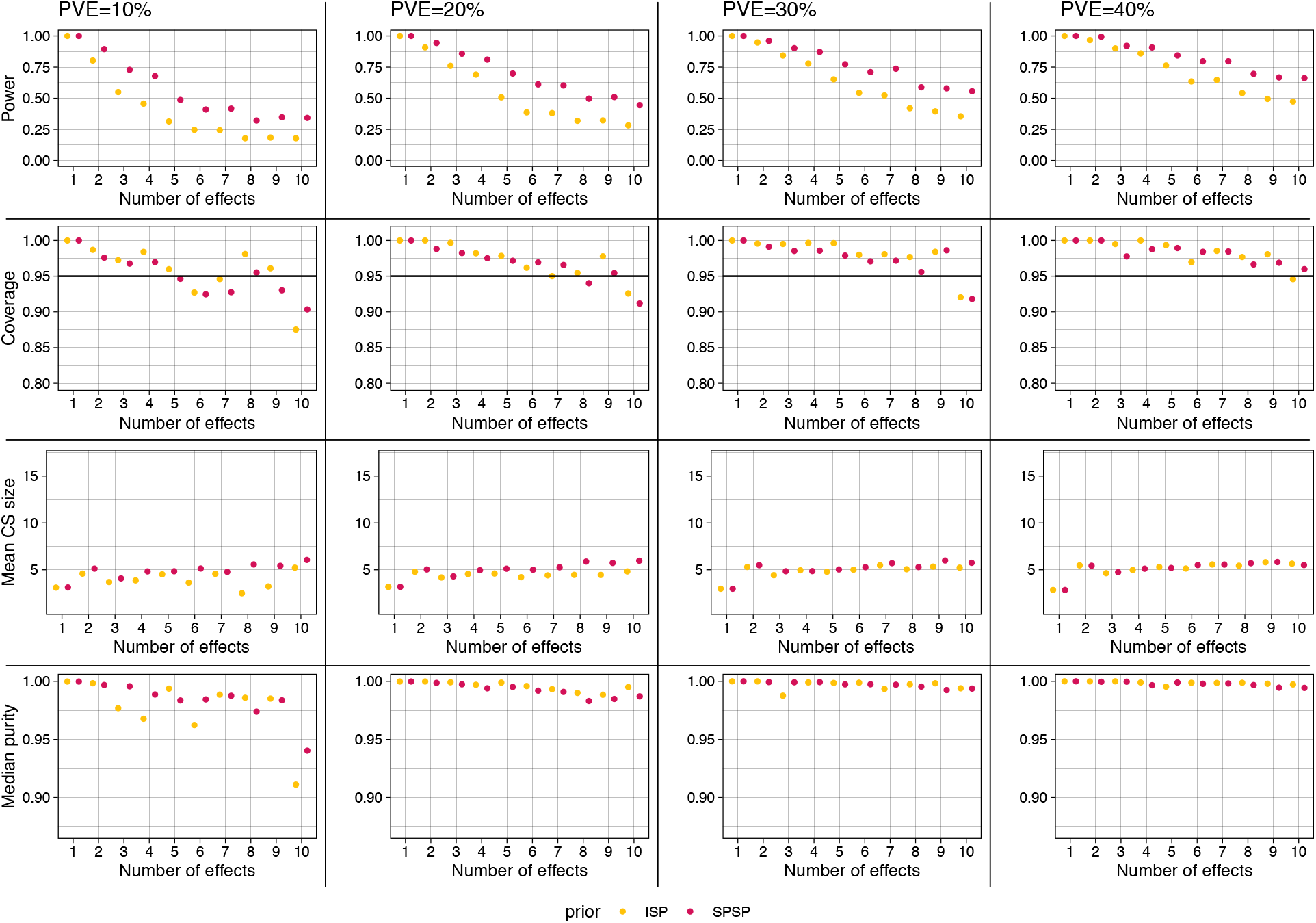
Additional assessment of fSuSiE CSs in wavelet simulations with fractional Gaussian noise, 512 CpGs. “Purity” is the smallest absolute correlation (Pearson’s *r*) among all pairs of SNPs in a CS [24]. See the Fig. 3 caption for definitions of power, coverage and CS size. ISP = fSuSiE with IS prior; SPSP = fSuSiE with SPS prior. Each plot summarizes the results from *n* = 300 simulations. The data sets were simulated with *N* = 100 samples, and *J* = 1,500–4,000 SNPs.

**Supplementary Figure 7.**
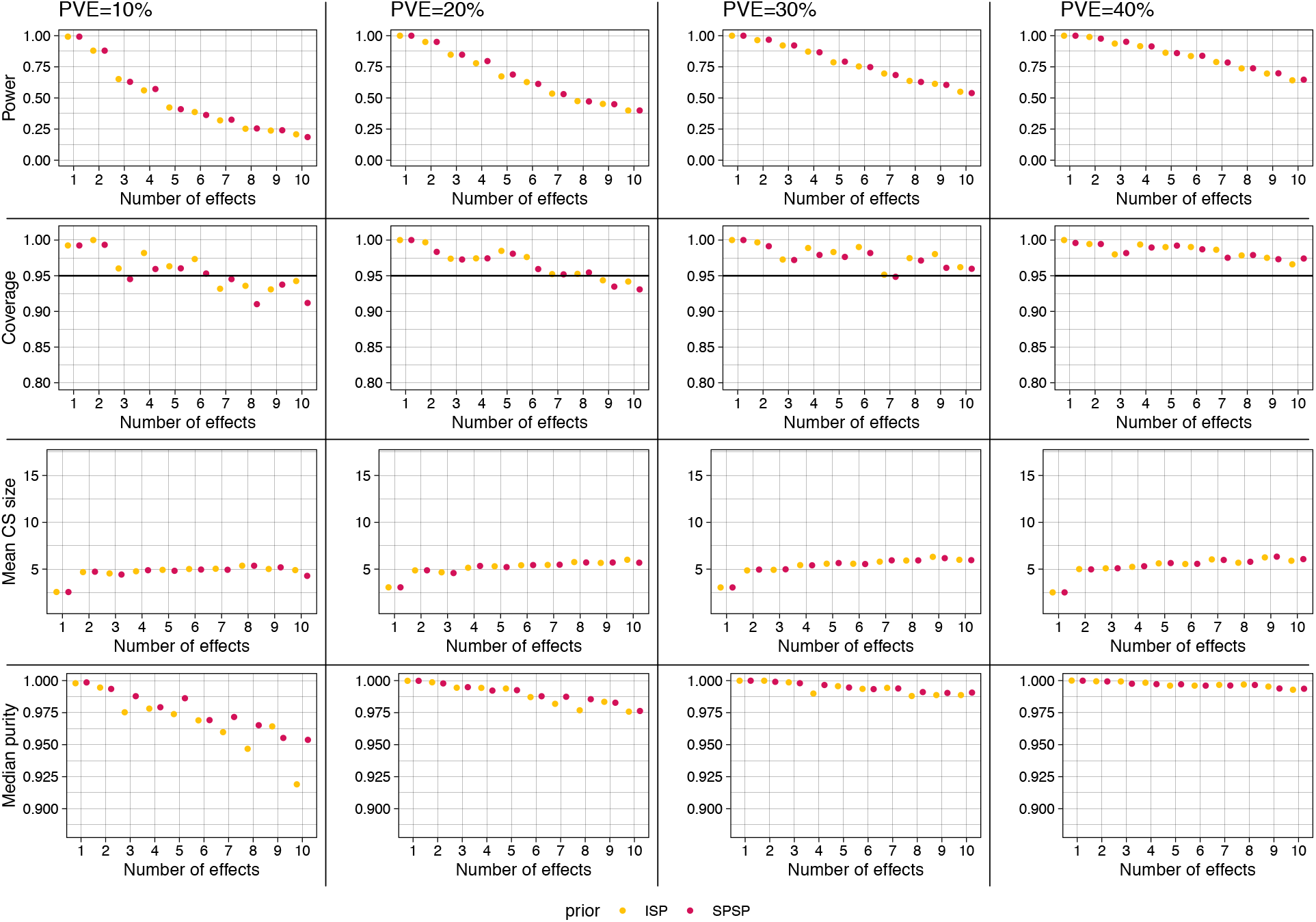
Additional assessment of fSuSiE CSs in wavelet simulations with GP noise, 128 CpGs. “Purity” is the smallest absolute correlation (Pearson’s *r*) among all pairs of SNPs in a CS [24]. See the Fig. 3 caption for definitions of power, coverage and CS size. ISP = fSuSiE with IS prior; SPSP = fSuSiE with SPS prior. Each plot summarizes the results from *n* = 300 simulations. The data sets were simulated with *N* = 100 samples, and *J* = 1,500–4,000 SNPs.

**Supplementary Figure 8.**
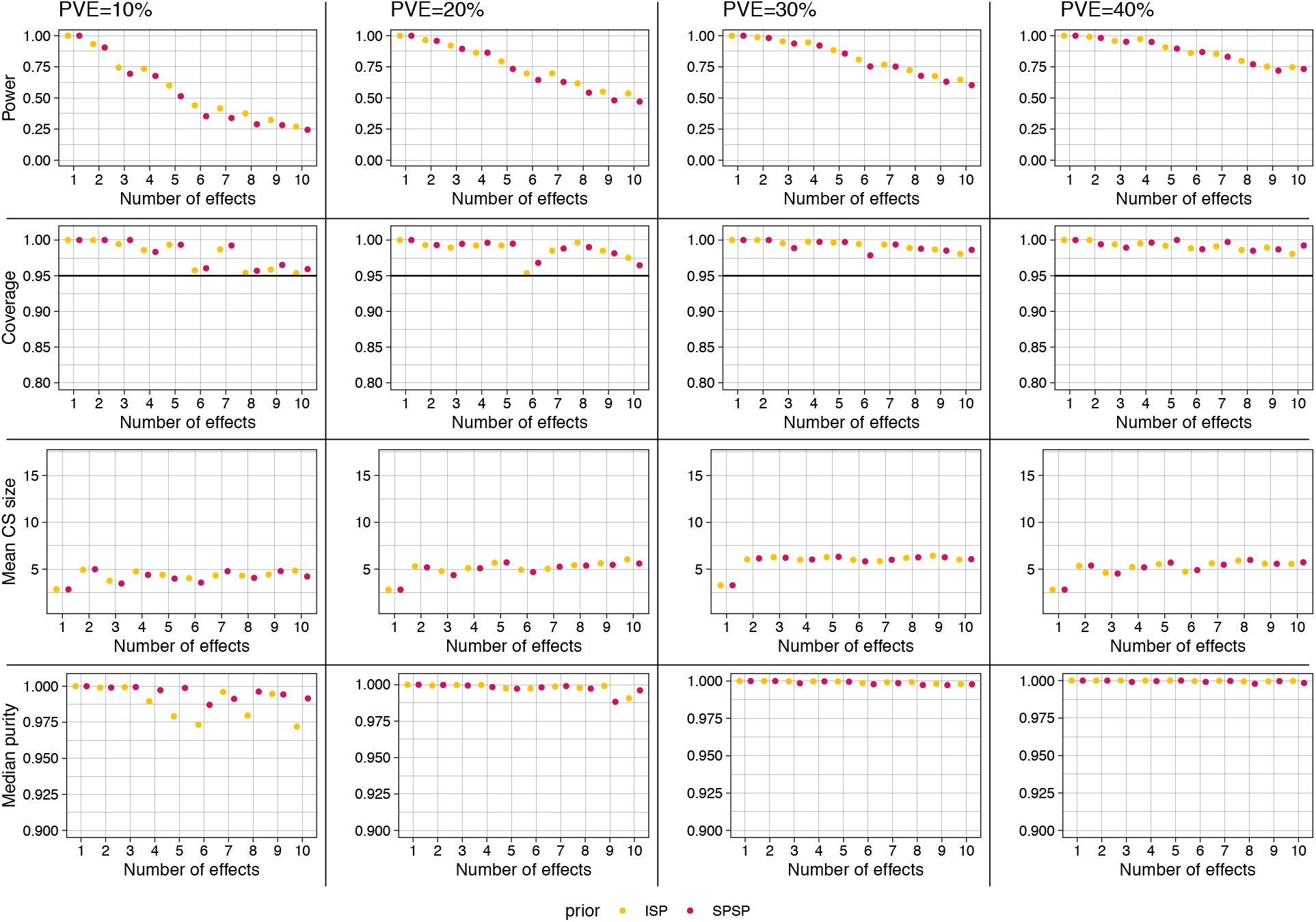
Additional assessment of fSuSiE CSs in wavelet simulations with GP noise, 512 CpGs. “Purity” is the smallest absolute correlation (Pearson’s *r*) among all pairs of SNPs in a CS [24]. See the Fig. 3 caption for definitions of power, coverage and CS size. ISP = fSuSiE with IS prior; SPSP = fSuSiE with SPS prior. Each plot summarizes the results from *n* = 300 simulations. The data sets were simulated with *N* = 100 samples, and *J* = 1,500–4,000 SNPs.

**Supplementary Figure 9.**
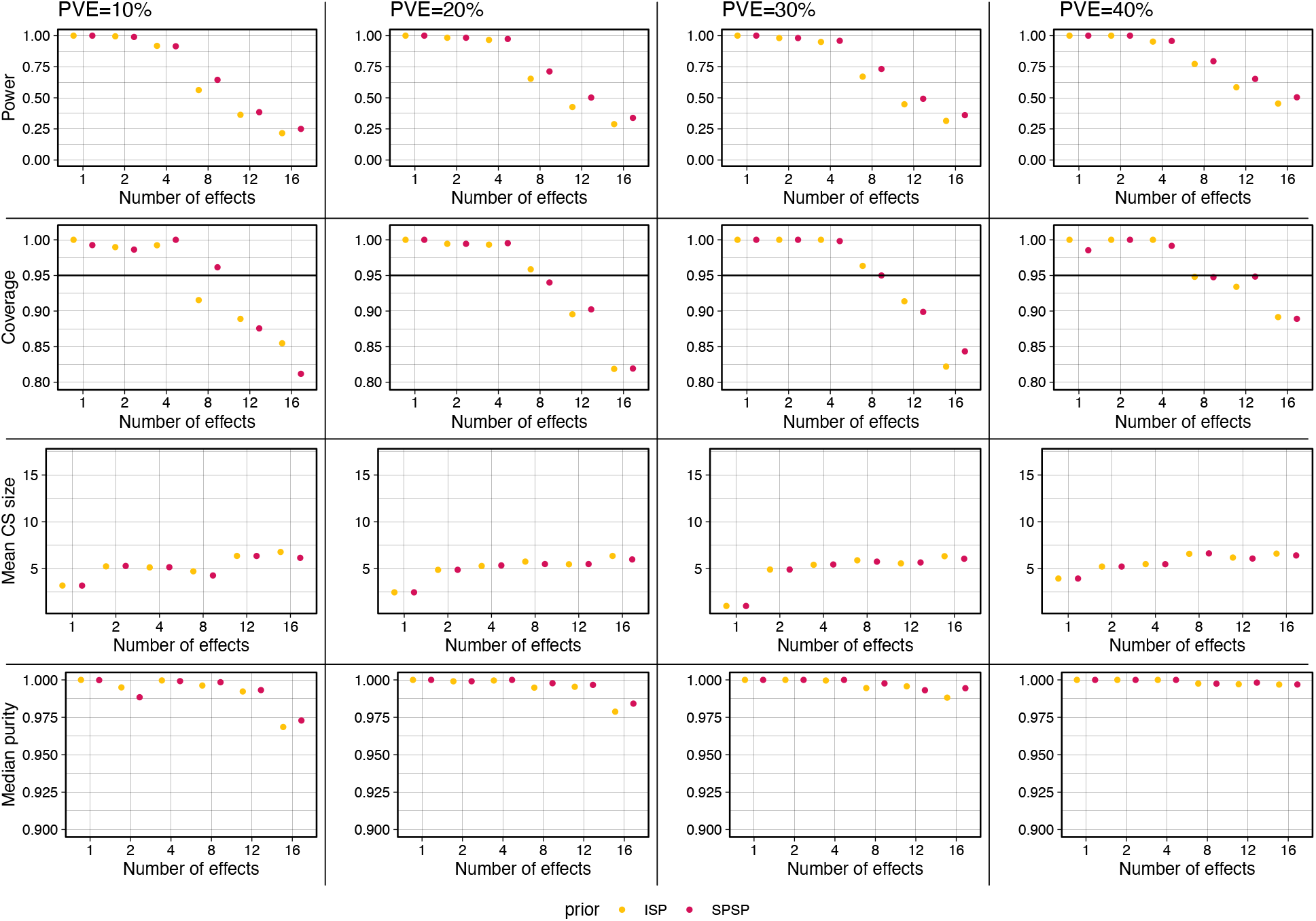
Additional assessment of fSuSiE CSs in WGBS block simulations with 128 CpGs. “Purity” is the smallest absolute correlation (Pearson’s *r*) among all pairs of SNPs in a CS [24]. See the Fig. 3 caption for definitions of power, coverage and CS size. ISP = fSuSiE with IS prior; SPSP = fSuSiE with SPS prior. Each plot summarizes the results from *n* = 300 simulations. The data sets were simulated with *N* = 100 samples, and *J* = 1,500–4,000 SNPs.

**Supplementary Figure 10.**
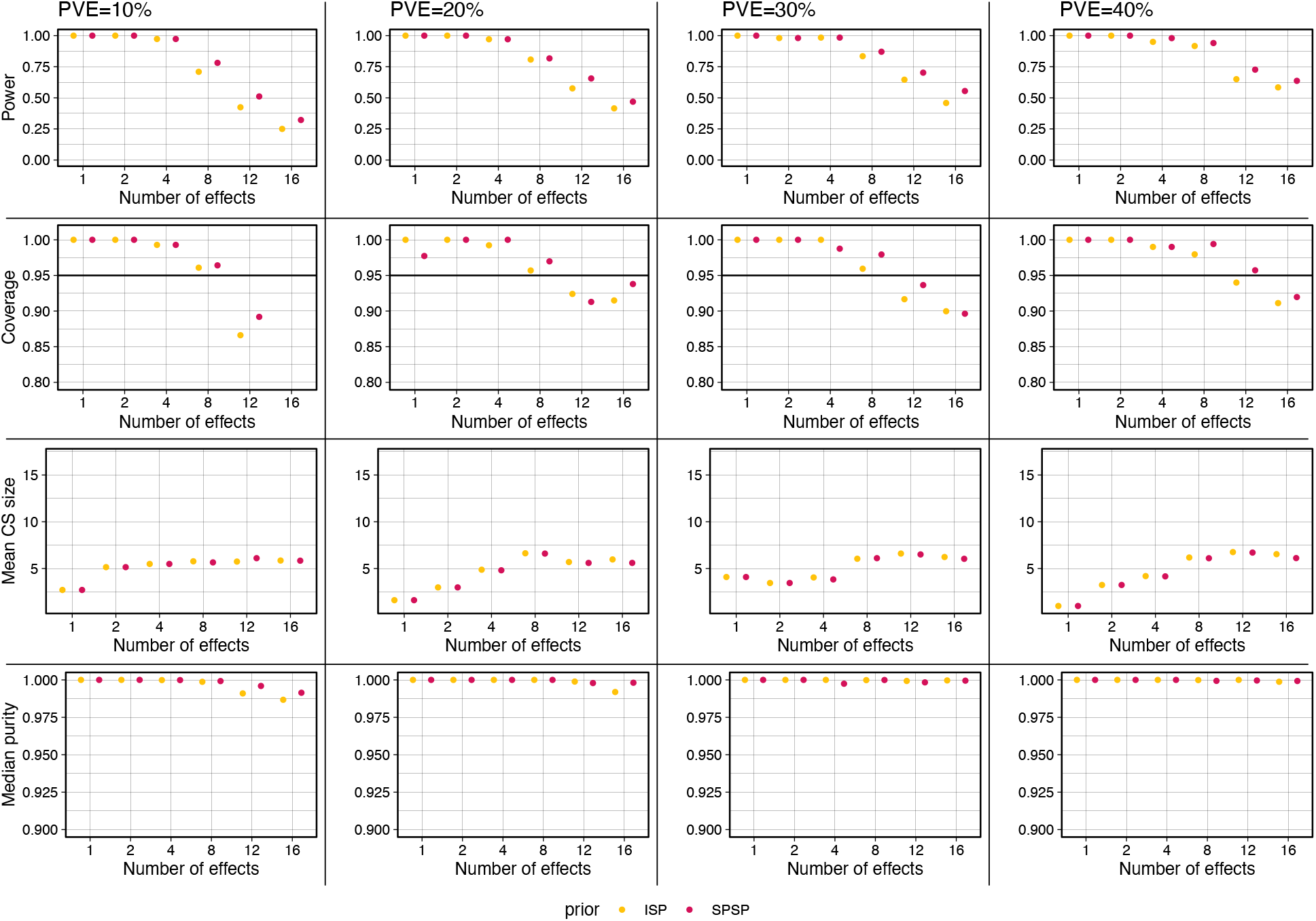
Additional assessment of fSuSiE CSs in WGBS block simulations with 512 CpGs. “Purity” is the smallest absolute correlation (Pearson’s *r*) among all pairs of SNPs in a CS [24]. See the Fig. 3 caption for definitions of power, coverage and CS size. ISP = fSuSiE with IS prior; SPSP = fSuSiE with SPS prior. Each plot summarizes the results from *n* = 300 simulations. The data sets were simulated with *N* = 100 samples, and *J* = 1,500–4,000 SNPs.

**Supplementary Figure 11.**
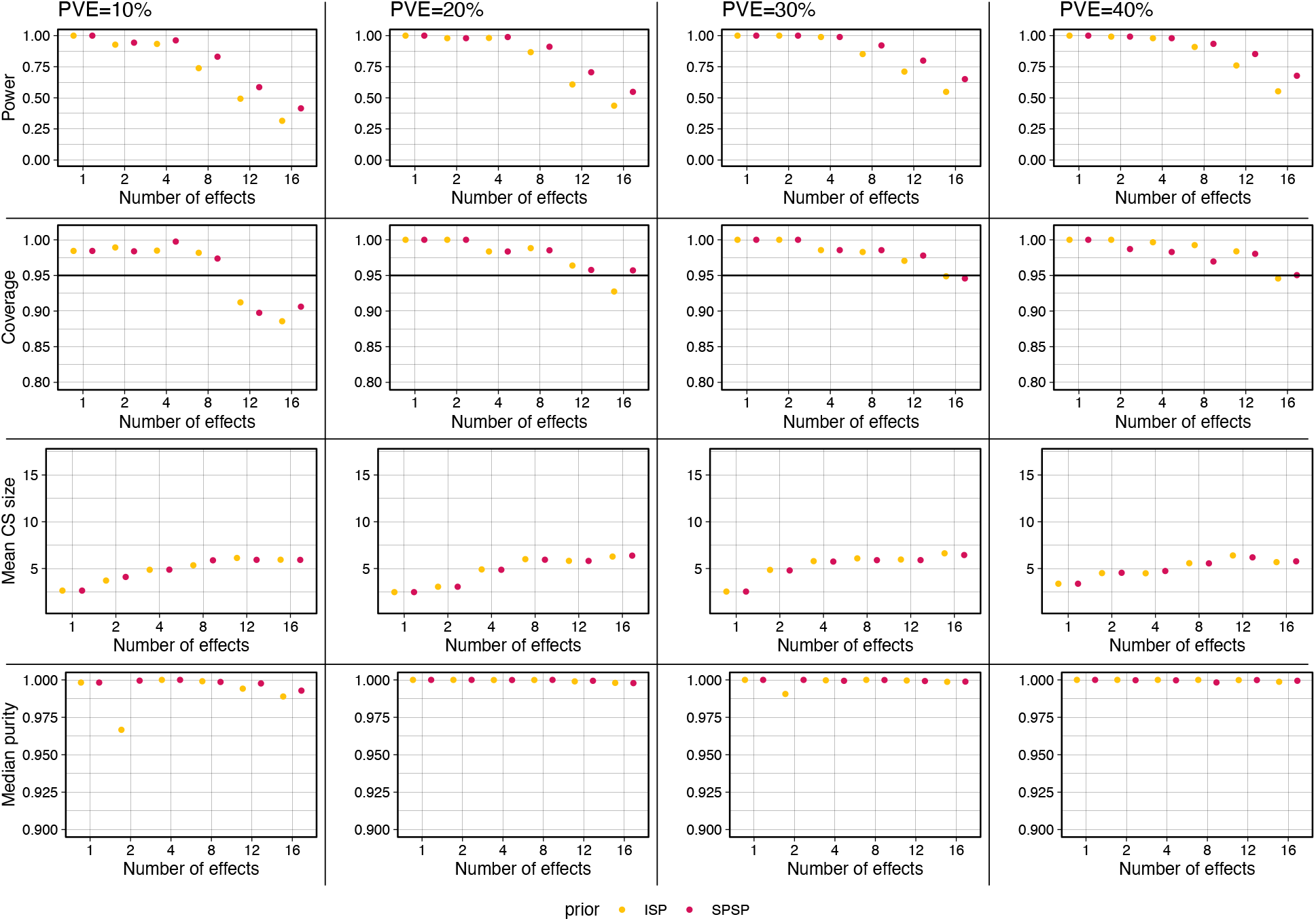
Additional assessment of fSuSiE CSs in WGBS decay simulations with 128 CpGs. “Purity” is the smallest absolute correlation (Pearson’s *r*) among all pairs of SNPs in a CS [24]. See the Fig. 3 caption for definitions of power, coverage and CS size. ISP = fSuSiE with IS prior; SPSP = fSuSiE with SPS prior. Each plot summarizes the results from *n* = 300 simulations. The data sets were simulated with *N* = 100 samples, and *J* = 1,500–4,000 SNPs.

**Supplementary Figure 12.**
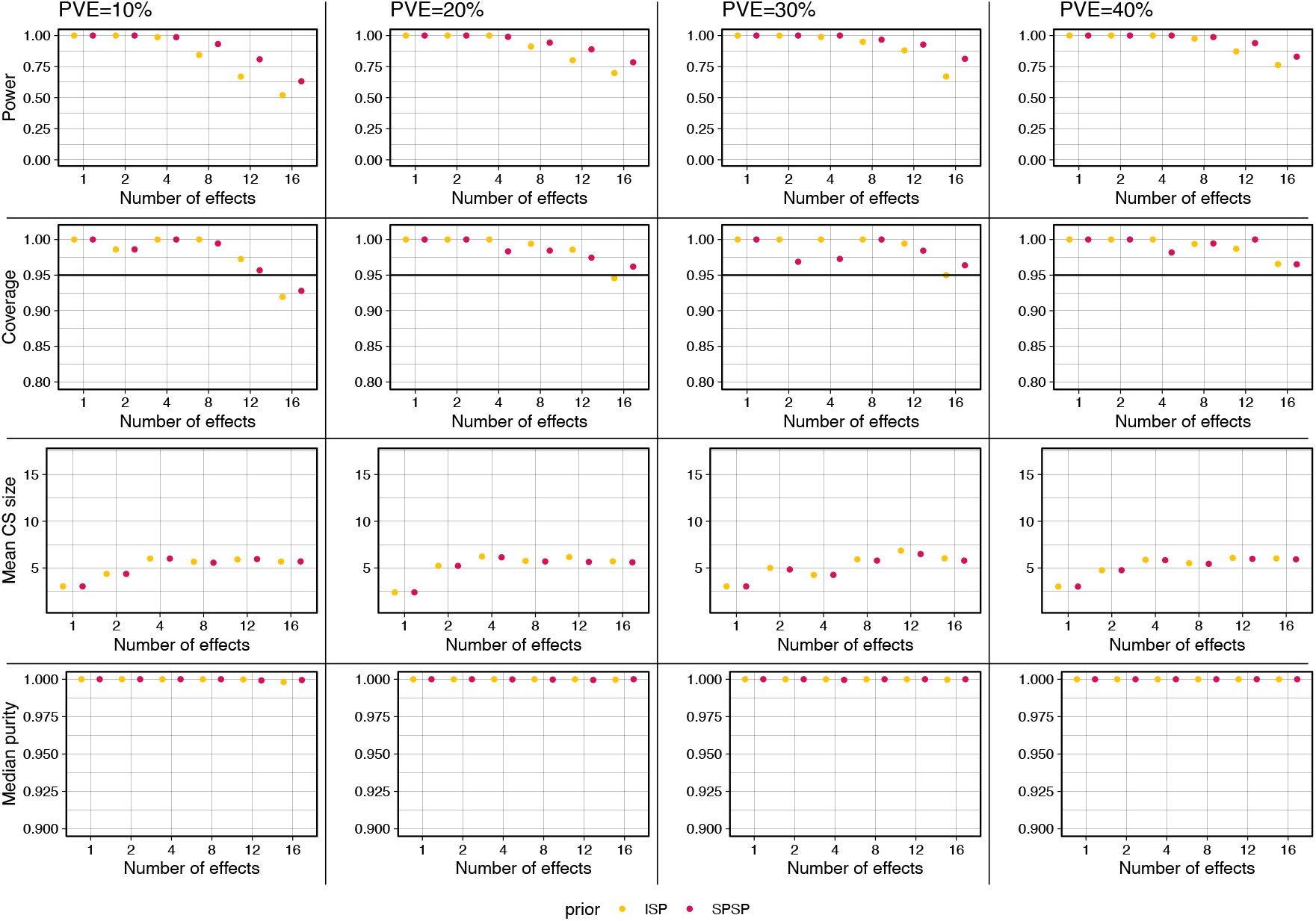
Additional assessment of fSuSiE CSs in WGBS decay simulations with 512 CpGs. “Purity” is the smallest absolute correlation (Pearson’s *r*) among all pairs of SNPs in a CS [24]. See the Fig. 3 caption for definitions of power, coverage and CS size. ISP = fSuSiE with IS prior; SPSP = fSuSiE with SPS prior. Each plot summarizes the results from *n* = 300 simulations. The data sets were simulated with *N* = 100 samples, and *J* = 1,500–4,000 SNPs.

**Supplementary Figure 13.**
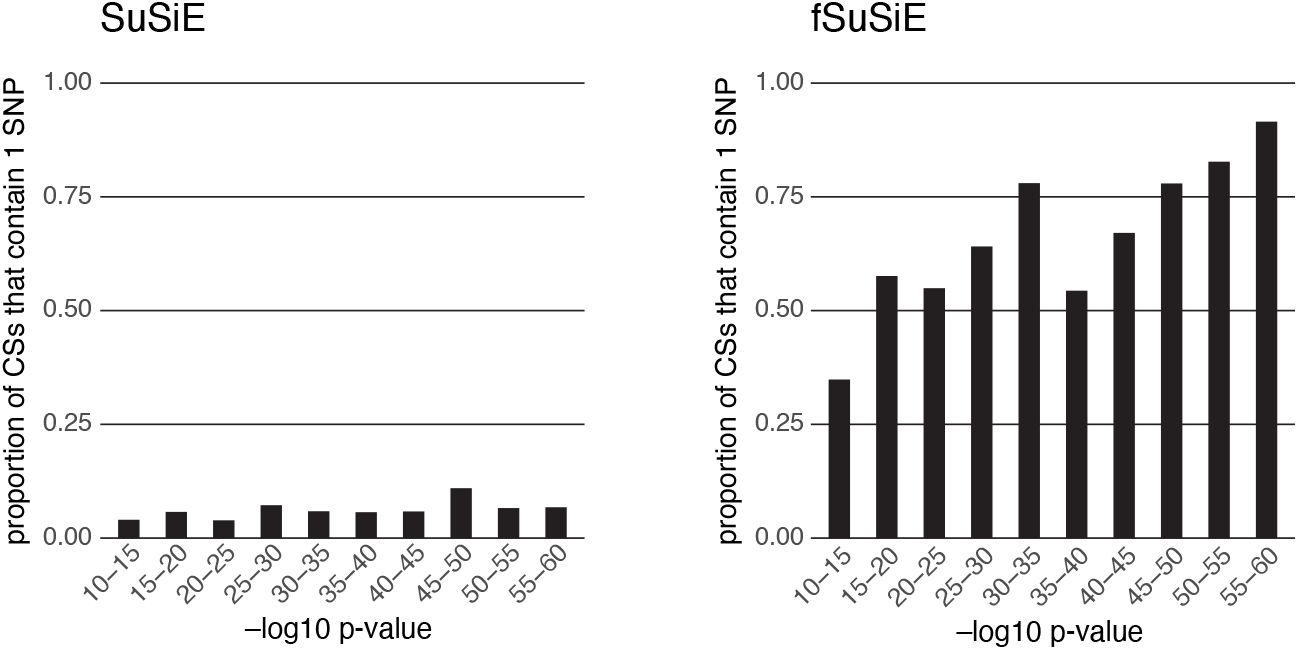
Results of the 2-SNP simulations. “*p*-value” is the smallest *p*-value from association tests between SNP 1 and molecular locations 1–128.

**Supplementary Figure 14.**
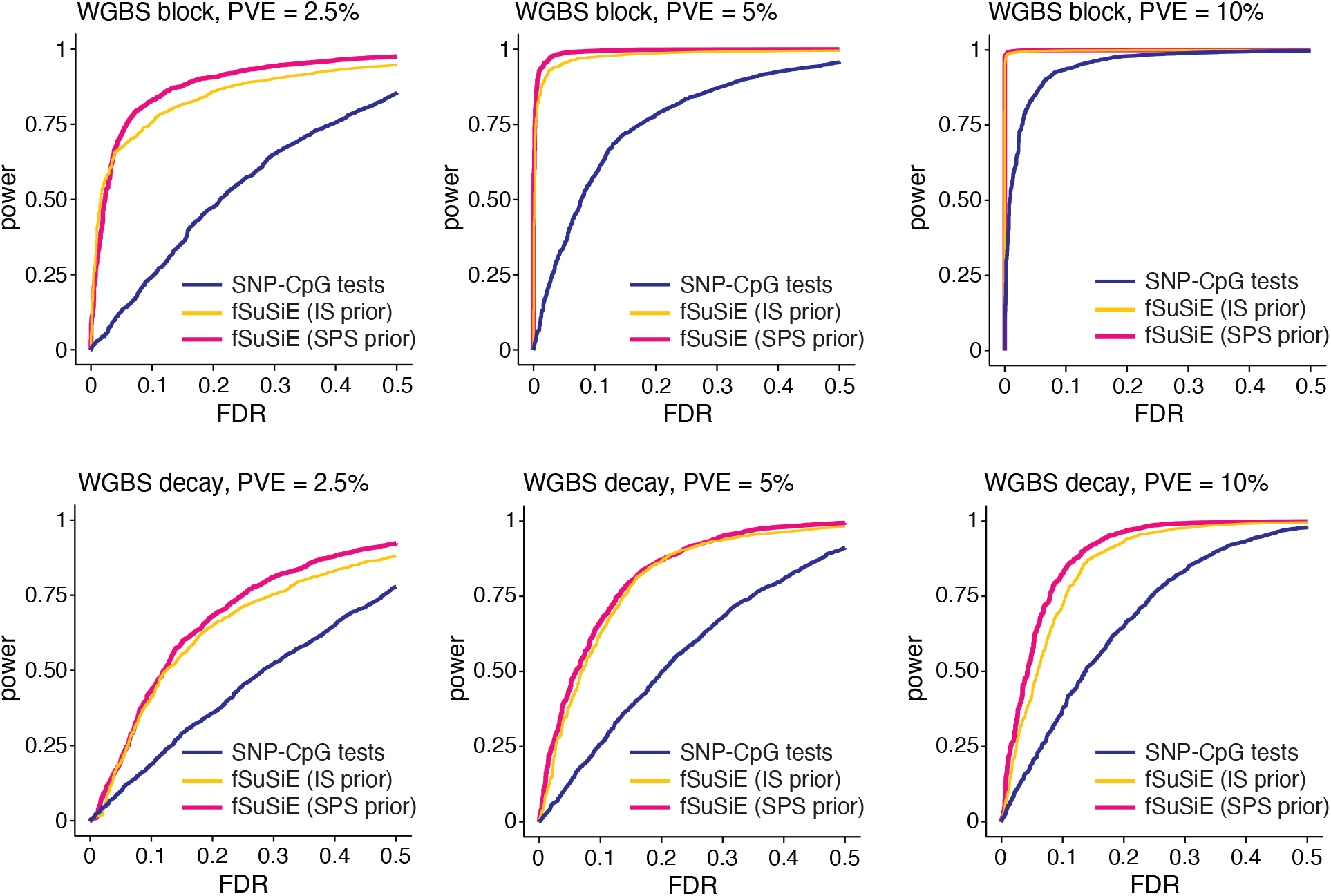
Additional assessment of recovery of affected CpGs in simulated data sets. The power-FDR plots evaluate recovery of affected CpG in which the methylation levels were simulated with different total variance explained by the SNPs (PVE = proportion of variance explained). See the Fig. 3 caption for an explanation of the power-FDR plots. In each of the simulation scenarios, *n* = 300 data sets were simulated with *N* = 100 samples, *T* = 128 CpGs, and *J* = 1,500–4,000 SNPs, among which 1–20 were causal.

**Supplementary Figure 15.**
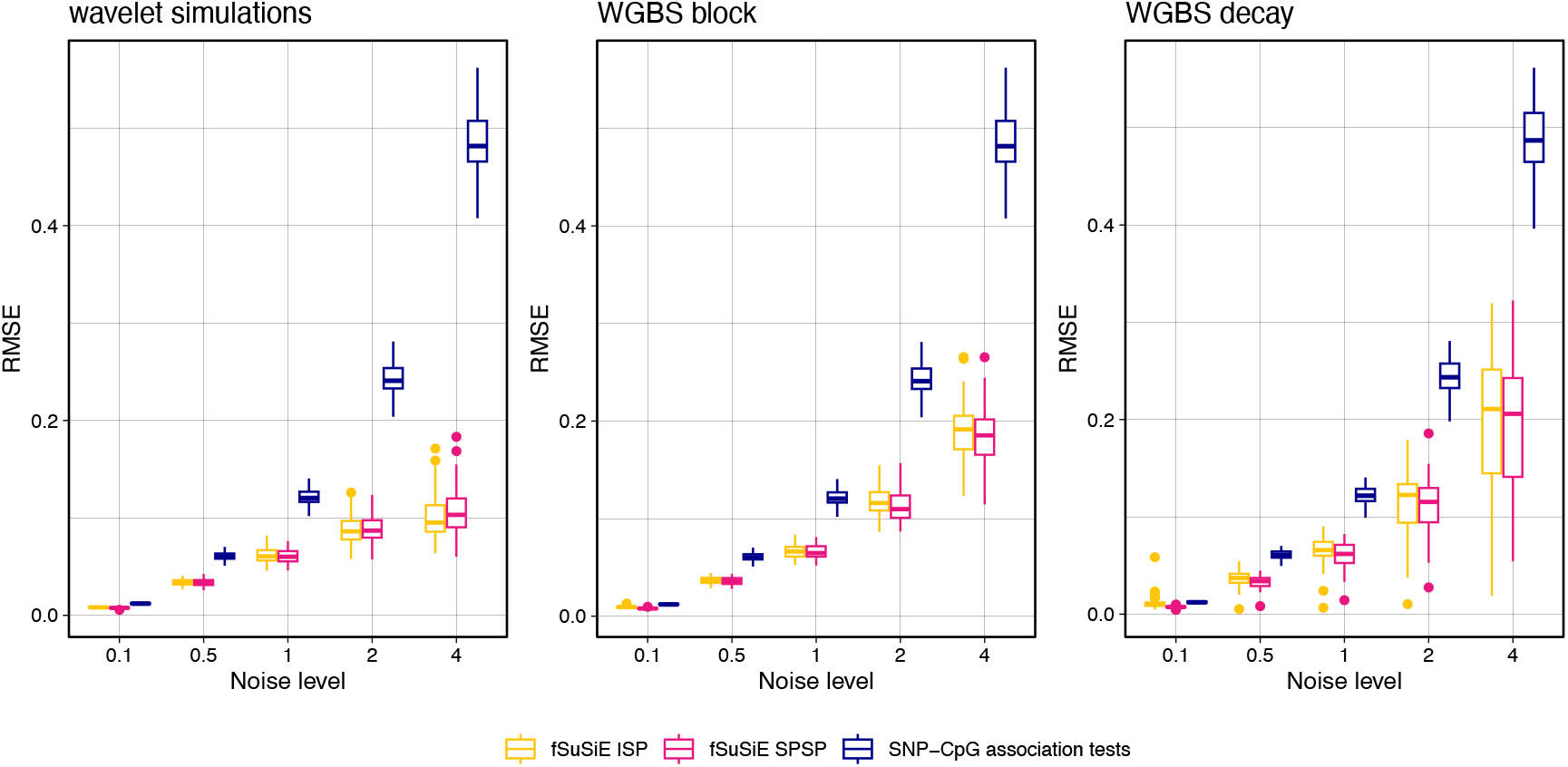
Accuracy of effect estimates in simulations. In addition to assessing recovery of the affected CpGs (Fig. 3D), we also assessed accuracy of the estimated effects of genotype on methylation levels at all CpGs. “Accuracy” in a data set is summarized by the root mean squared error, 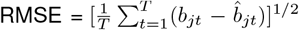, where *T* is the number of CpGs, 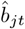 is the estimated effect and *b*_*jt*_ is the true effect (*b*_*jt*_ = 0 at unaffected CpGs). Note that *j* is always 1 in these simulations since there is only 1 SNP. Each box plot summarizes accuracy across *n* = 50 simulations: the box plot whiskers depict 1.5× the interquartile range, the box bounds represent the upper and lower quartiles (25th and 75th percentiles), the center line represents the median (50th percentile), and points represent outliers. The data sets were simulated with *N* = 100 samples, *T* = 128 CpGs and *J* = 1 causal SNP.

**Supplementary Figure 16.**
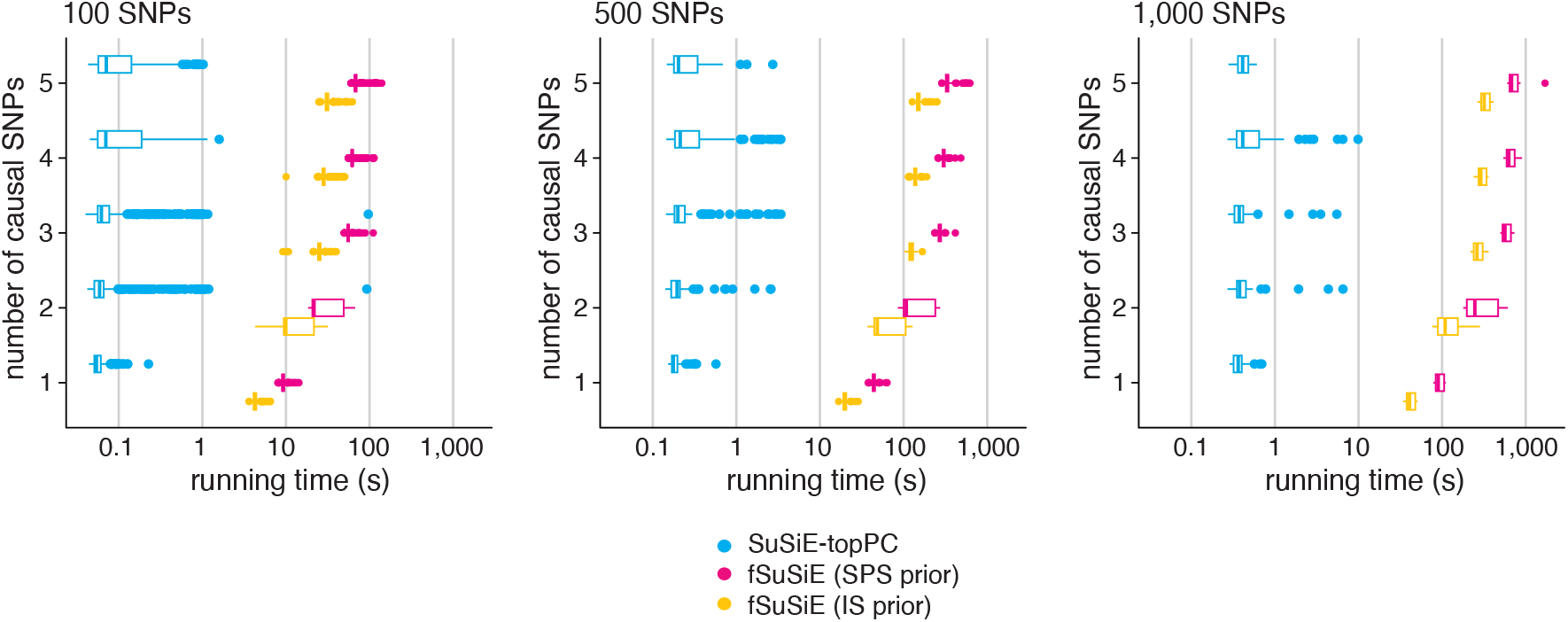
Additional running times in simulations. Running times in simulated data sets with 100, 500 and 1,000 SNPs (each plot summarizes results from *n* = 300 simulations). The box plot whiskers depict 1.5× the interquartile range, the box bounds represent the upper and lower quartiles (25th and 75th percentiles), the center line represents the median (50th percentile), and points represent outliers. The data sets were simulated with *N* = 100 samples, *T* = 128 CpGs, 1–20 causal SNPs, and 10% total variance explained.

**Supplementary Figure 17.**
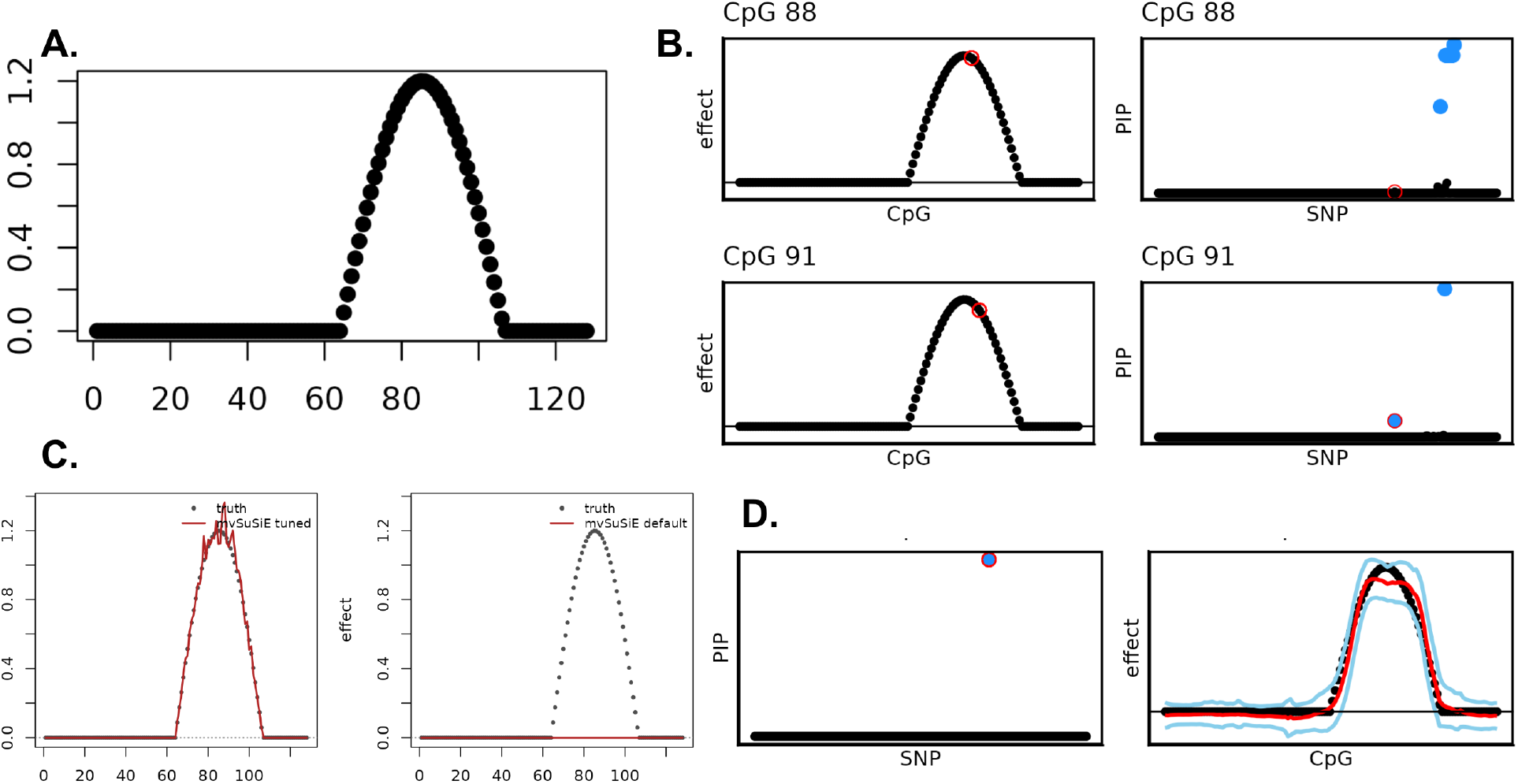
Cmparison of SuSiE, mvSuSiE, and fSuSiE on a single simulated locus with a spatially structured genetic effect. An example simulation replicate with *N* = 100 samples, *T* = 128 positions, and *J* = 991 candidate SNPs; a single causal SNP (index 691) has a cosine-shaped effect spanning 88 of the 128 positions, with independent Gaussian residuals. **(A)** Ground-truth effect of the causal SNP along genomic positions. **(B)** Univariate SuSiE applied independently at each position. Only a small subset of affected positions yields a 95% credible set, and the lead variants differ across adjacent positions even when the underlying effect is similar in amplitude, illustrating the difficulty of aggregating per-position fine-mapping into a single coherent signal. **(C)** mvSuSiE with two prior/residual configurations. *Left:* true effect-size prior and true residual variance, in which case mvSuSiE recovers the causal SNP at PIP = 1. *Right:* canonical MASH prior [33] with a default-initialized residual variance, in which case mvSuSiE returns PIP = 0 and no credible set. **(D)** fSuSiE recovers the causal SNP at PIP = 1 and reconstructs a smooth effect curve along the genome (solid line) with a 95% credible band (shaded region) that closely tracks the ground truth (grey dots), without requiring problem-specific specification of the prior or residual variance.

**Supplementary Figure 18.**
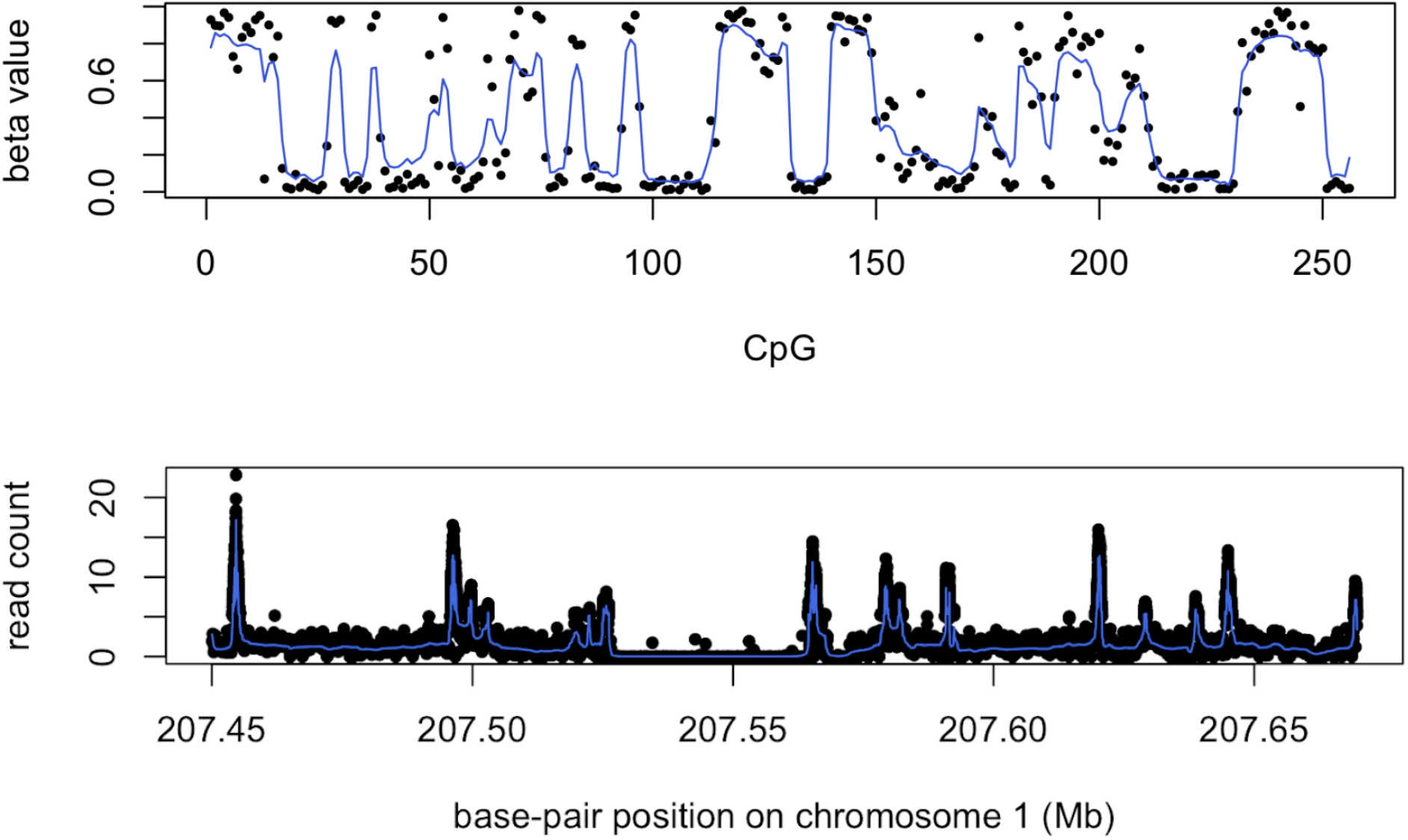
Empirical illustration of spatial structure in molecular trait data from ROSMAP DLPFC. Top: Illumina Infinium HumanMethylation450 BeadChip methylation *β* values across the CASS4 fine-mapping region on chromosome 20 (the same locus highlighted in Fig. 6). Bottom: H3K9ac ChIP-seq read counts across the *CR1*/*CR2* fine-mapping region on chromosome 1 (also highlighted in Fig. 6). The two assays produce signals of different shape and dynamic range, but in both cases positions of high activity (*β* values near 1, or large read counts) tend to cluster along the chromosome. A reproducible walkthrough using the smashr R package on these data is available at https://stephenslab.github.io/fsusie-experiments/smash_meth_ha.html.

**Supplementary Figure 19.**
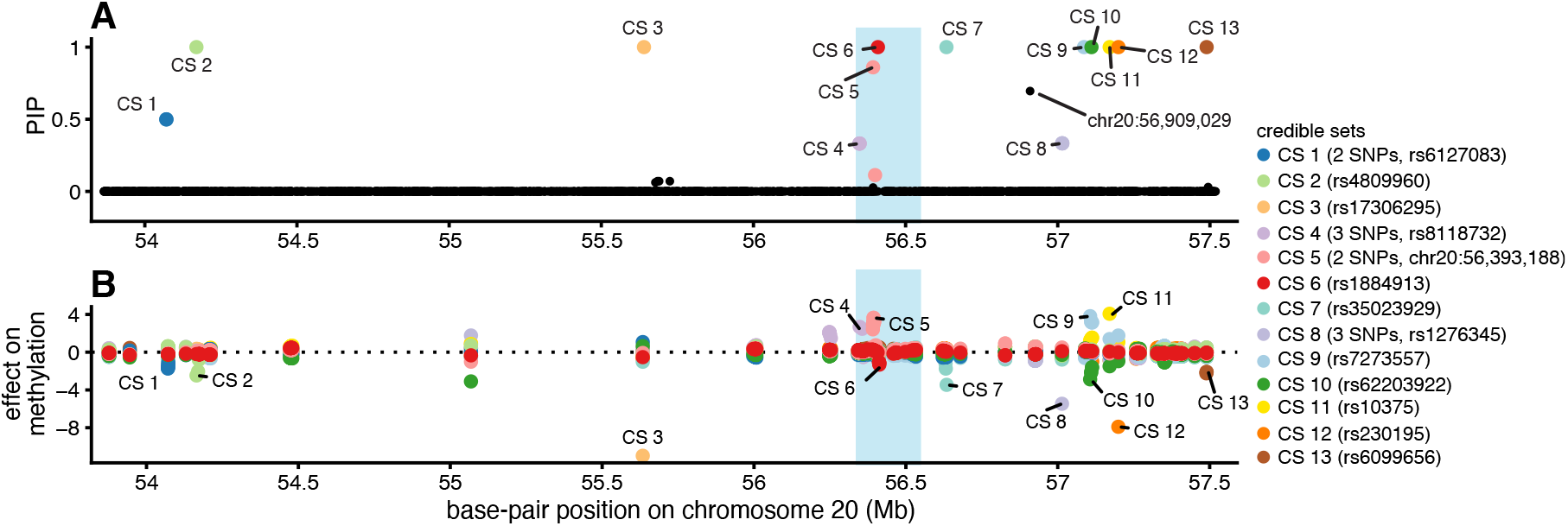
fSuSiE methylation fine-mapping example on chromosome 20. Panel A shows the PIPs for all SNPs analyzed in the TAD (chromosome 20, 53.9––57.5 Mb (20,320 SNPs, 300 CpGs). The PIP is an estimate of the probability that the SNP affects methylation levels at one or more peaks within the TAD. The colors indicate the different mSNPs, and the uncertainty in which SNP is the mSNP is represented by the 95% CS: each SNP in the CS is a candidate mSNP, with probability of being causal given by the PIP. More detail about the CSs is given on the right-hand side: the number of SNPs in the CS (if not given, then it is a single-variant CS); and the sentinel SNP, the SNP with the highest PIP in the CS. Note that a single SNP at base-pair position 56,909,029 with PIP = 0.70 is not included in a CS because the CS was removed by the CS purity filter (Online Methods). Panel B shows the effect of each mSNP on methylation at each of the peaks analyzed in the TAD. See Fig. 4 for an explanation of how to interpret this plot. The results within the blue-shaded region are examined in more detail in Fig. 6.

**Supplementary Figure 20.**
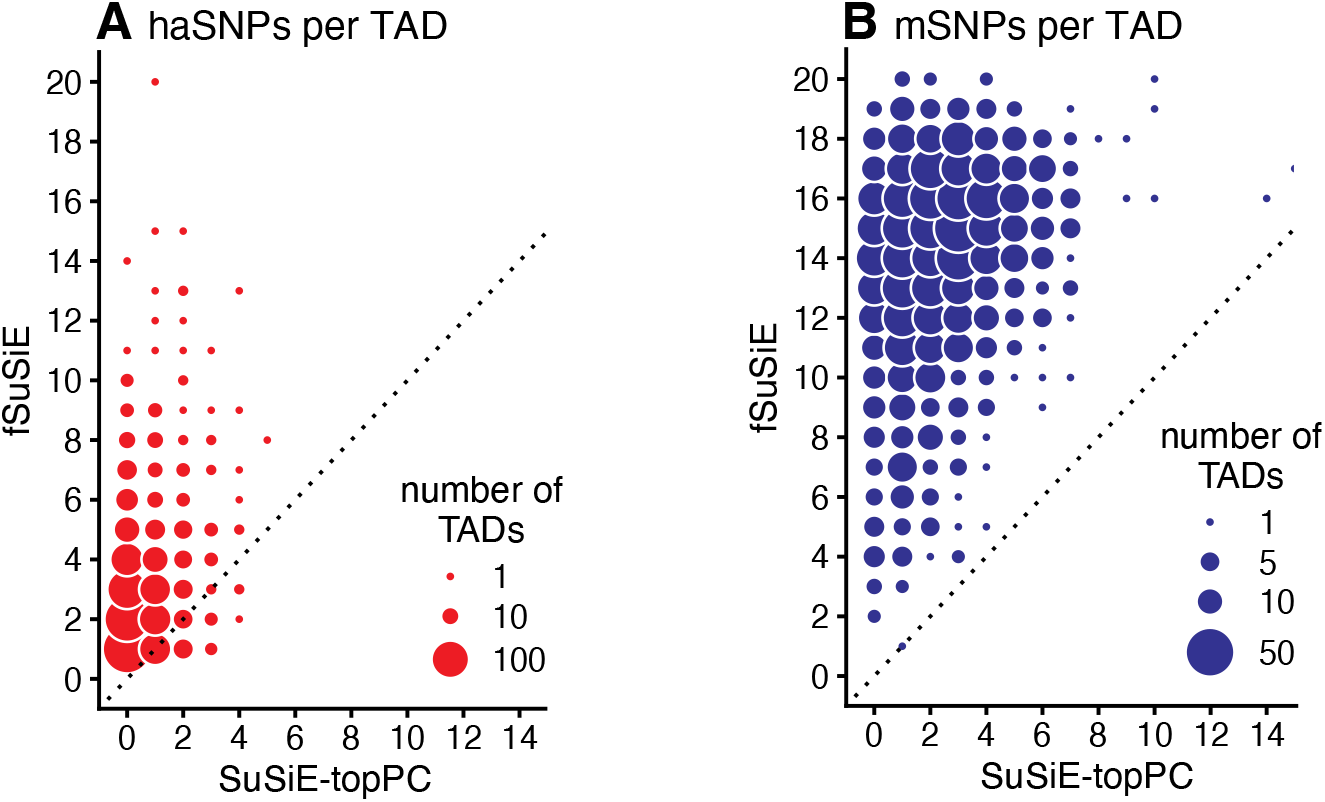
Additional results on fine-mapping of mSNPs and haSNPs in DLPFC. These plots compare the number of haSNPs and mSNPs (i.e., the number of 95% CSs) returned by SuSiE-topPC and fSuSiE in each TAD.

**Supplementary Figure 21.**
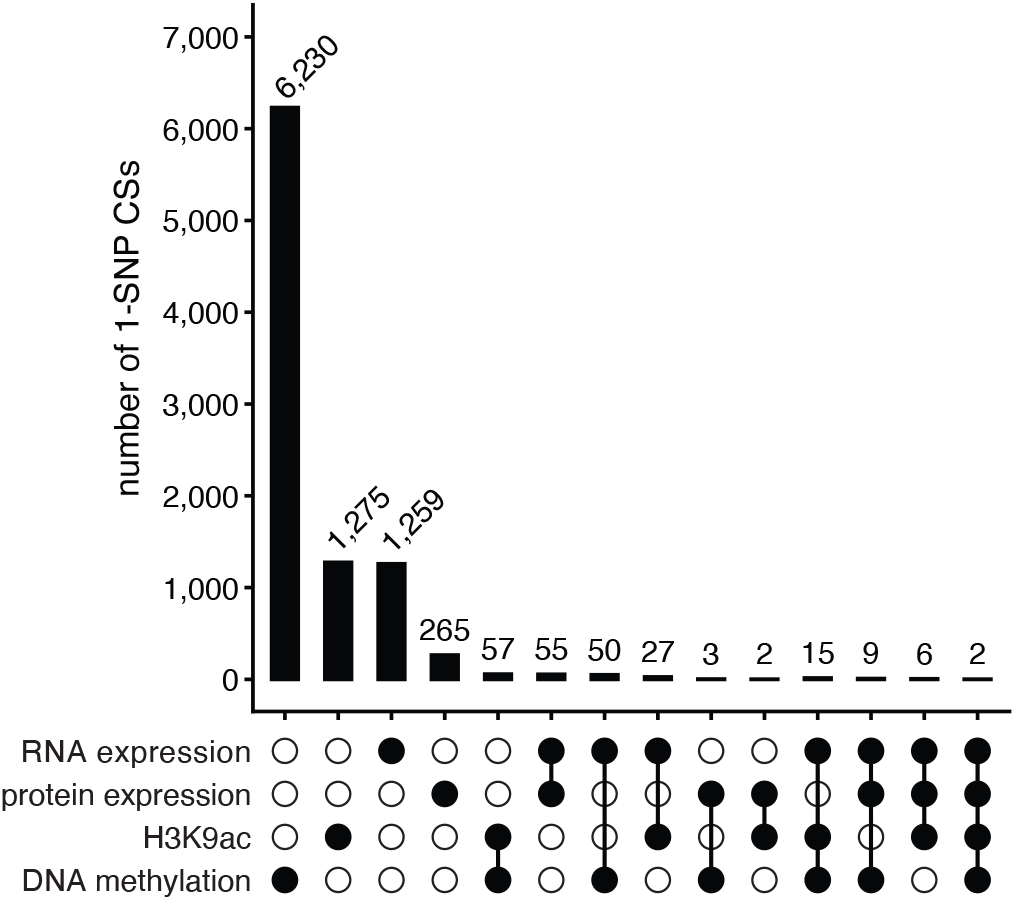
Overlap among molecular trait SNPs in DLPFC. Overlap is determined based on base-pair position of SNPs in the single-variant CSs returned by SuSiE (RNA expression, protein expression) and fSuSiE (H3K9ac, DNA methylation). For reference, the total number of single-variant CSs for each trait is: RNA expression (1,420), protein expression (340), H3K9ac (1,372) and methylation (6,355).

**Supplementary Figure 22.**
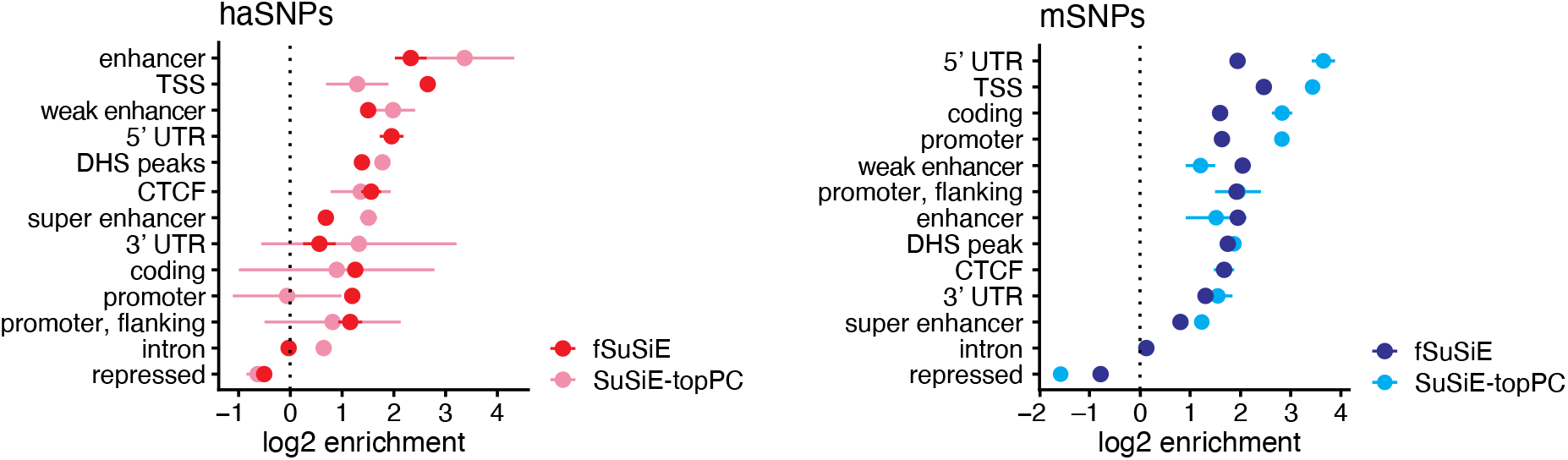
Additional results on functional enrichment of haSNPs and mSNPs. Functional enrichment of the mSNPs and haSNPs is assessed by by functional annotations [80, 81]. See Online Methods for details on how enrichment is quantified. Points and error bars represent means and standard errors estimated from jackknife resampling.

**Supplementary Figure 23.**
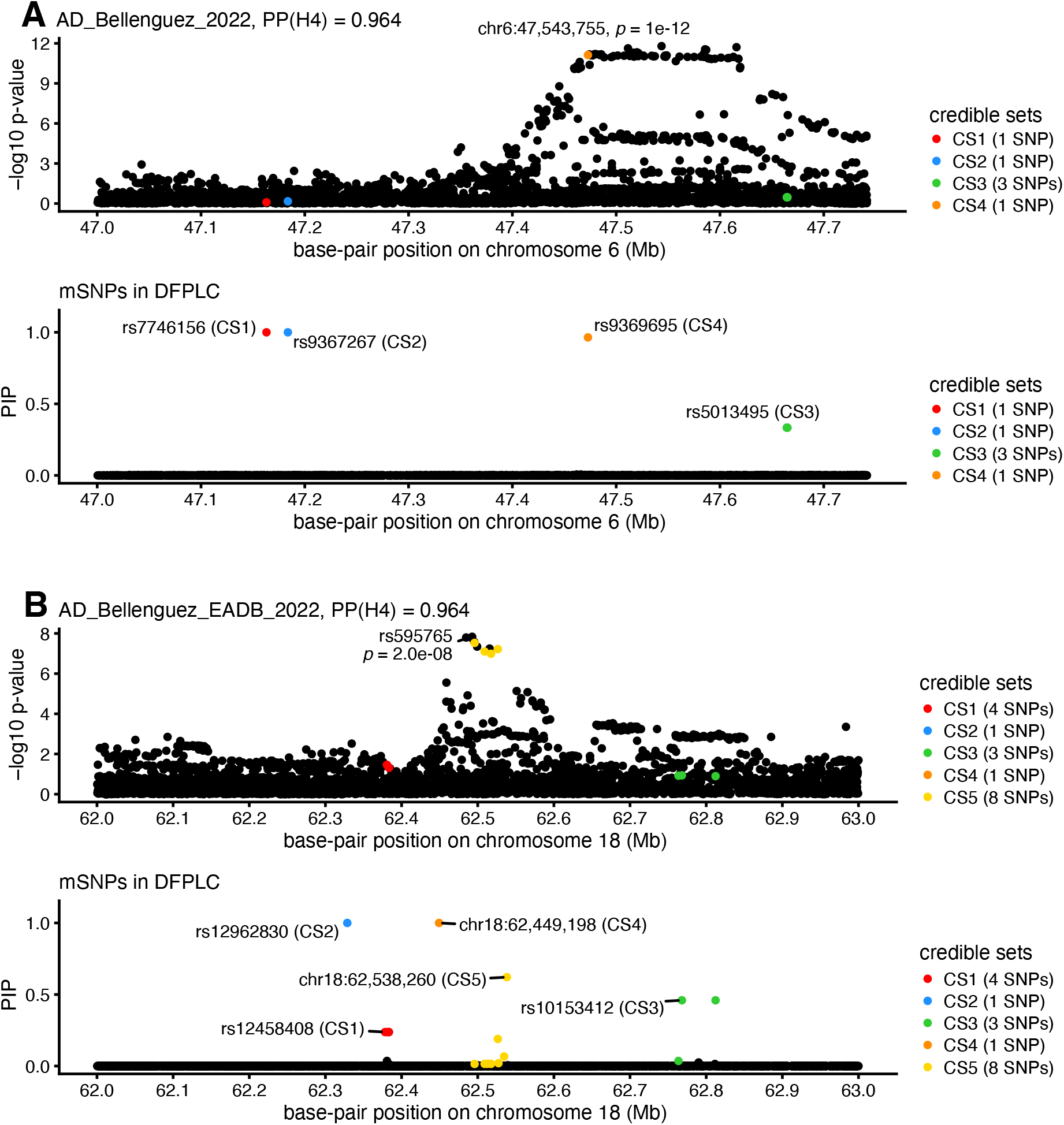
Additional examples of mSNPs colocalizing with AD risk variants [83]. For each CS, the sentinel SNP is labeled. The SNP with the smallest AD association test *p-*value (two-sided *t*-test) is also labeled.

**Supplementary Figure 24.**
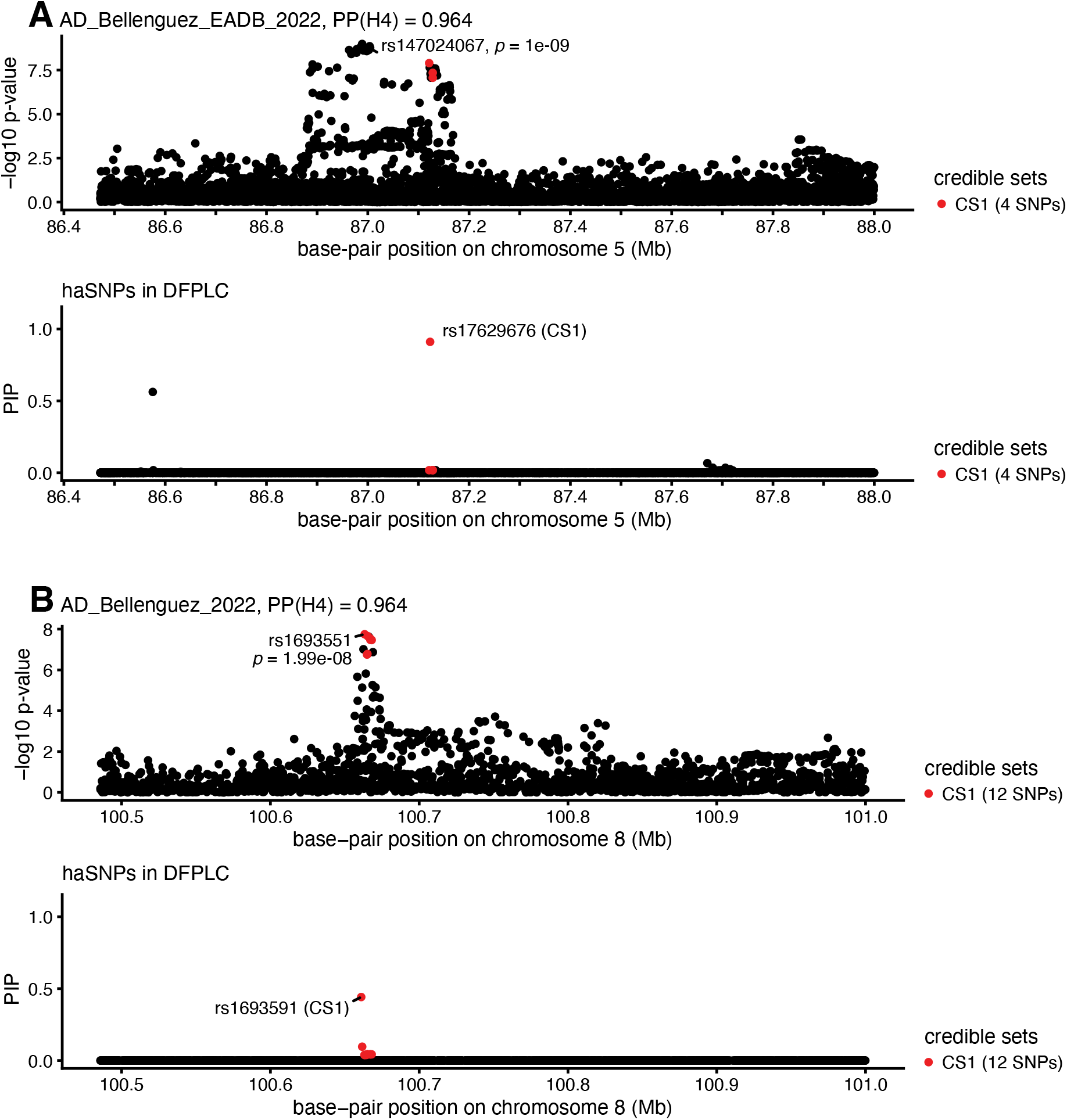
Additional examples of haSNPs colocalizing with AD risk variants [83]. For each CS, the sentinel SNP is labeled. The SNP with the smallest AD association test *p-*value (two-sided *t*-test) is also labeled.

